# Parvalbumin interneurons gate and shape striatal sequences

**DOI:** 10.1101/2025.11.08.687375

**Authors:** Mariana Duhne, Isabelle Gonzalez Montalvo, Chelsea S. Zheng, Lilian Pelattini, Joshua D. Berke

**Affiliations:** Department of Neurology; San Francisco, USA; Weill Institute for Neurosciences; San Francisco, USA; Department of Psychiatry and Behavioral Sciences, University of California, San Francisco; San Francisco, USA

## Abstract

Impulsive and uncontrolled behaviors are associated with loss of parvalbumin-expressing striatal interneurons (PV+). To understand this microcircuit mechanism we compared spiking of five classes of identified neurons in sensorimotor striatum, as rats waited for a cue, then performed brief actions. During waiting PV+ increased firing, and suppression of PV+ increased premature movements, supporting a role in action restraint. Once underway, each action was accompanied by a rapid striatal firing sequence, including PV+ and direct and indirect pathway spiny projection neurons (SPNs). Nearby PV+ and SPNs showed millisecond-level synchrony, and PV+ firing inhibited SPNs ∼2ms later. Suppressing PV+ immediately changed which SPNs participated in sequences, and slowed movements. PV+ thus provide both broad restraint and precise sculpting of striatal output to achieve fluid, appropriately timed behavior.

## Main Text

How the components of striatal circuits work together to influence behavior, and how this coordination is disrupted in neurological and psychiatric disorders, is unknown. Numerically the striatum is dominated by GABAergic spiny projection neurons (SPNs). Of these about half express dopamine D1 receptors (D1+ SPNs) and form the “direct” pathway, while the others express D2 and adenosine A2a receptors (D2+ SPNs) and form the “indirect” pathway(*1*). Interneurons are relatively rare (∼5% in rodents) yet show high evolutionary conservation(*2*) and are critical regulators of striatal function. Cholinergic (ChAT+) interneurons influence many striatal circuit elements including local dopamine release(*3,4*), and are implicated in dyskinesias and depression(*5*). The remaining interneurons are GABAergic. While these have diverse properties(*6*), they are primarily either fast-spiking cells that express parvalbumin (PV+), or low-threshold-spiking cells that express somatostatin (SST+)(*7–9*). Deficits in striatal PV+ cells are associated with uncontrolled movements in Tourette Syndrome(*10*) and dystonia in Huntington’s Disease(*11*), and aberrant striatal PV+ activity is associated with excessive stereotyped movement in mouse models of autism spectrum disorder(*12*).

PV+ interneurons make synapses onto SPN cell bodies and proximal dendrites, and in brain slices and anesthetized rats have been shown to provide fast and powerful inhibition of SPNs(*13–15*). This influence has been suggested to be essential for normal behavioral control, in part by organizing SPN firing into functional cell assemblies(*16*). However testing such hypotheses requires recording from identified PV+ and SPNs in behaving animals with high temporal resolution, which has not been previously achieved. In the absence of positive identification, many studies (including from our group) have used the duration of extracellularly recorded spikes to distinguish presumed PV+ fast-spiking interneurons (“pFSIs”) from presumed projection cells(*17–20*). For cortex this classification approach is supported by evidence that some brief (narrow-waveform) units make fast inhibitory connections to nearby neurons(*21*). In the case of striatum however, prior studies have generally not found the expected rapid interactions between pFSIs and pSPNs(*18, 22*) and report that pFSIs have only limited consistency in their behavior-related activity(*23*).

Here we compare the spiking of five major neuronal classes within sensorimotor striatum, each recorded during the same behavioral task and positively identified using optogenetic tagging. Surprisingly, we report that identified striatal PV+ do not correspond to the commonly observed narrow-waveform striatal units. Furthermore we find that PV+ have highly distinct and consistent firing patterns, and contribute to both the suppression of impulsive actions and to the moment-by-moment temporal organization of SPNs during action execution.

## Results

### Optogenetically identified PV+ are not the expected narrow-waveform units

To identify neurons we used cell-type-specific expression of the excitatory opsin ChrimsonR(*24*). We infused an adeno-associated virus (AAV5-Syn-FLEX-rc[ChrimsonR-tdTomato]) into the sensorimotor (dorsal-lateral) striatum (DLS) of Cre-expressing rats (PV-Cre, SST-Cre, ChAT-Cre, D1-Cre, or A2a-Cre) (Fig. 1A). We then implanted chronic assemblies consisting of a circular array of tetrodes (4-wire microelectrodes) surrounding an optic fiber(*25–27*)(Fig. 1B). We recorded neuronal spiking (Fig. 1C), while applying brief pulses of red light through the fiber. Neurons that rapidly increased spiking (Fig. 1D) were identified as ChrimsonR-expressing, and hence either PV+, SST+, ChAT+, D1+, or D2+/A2a+ depending on rat line (see Methods for full criteria).

**Fig. 1.**
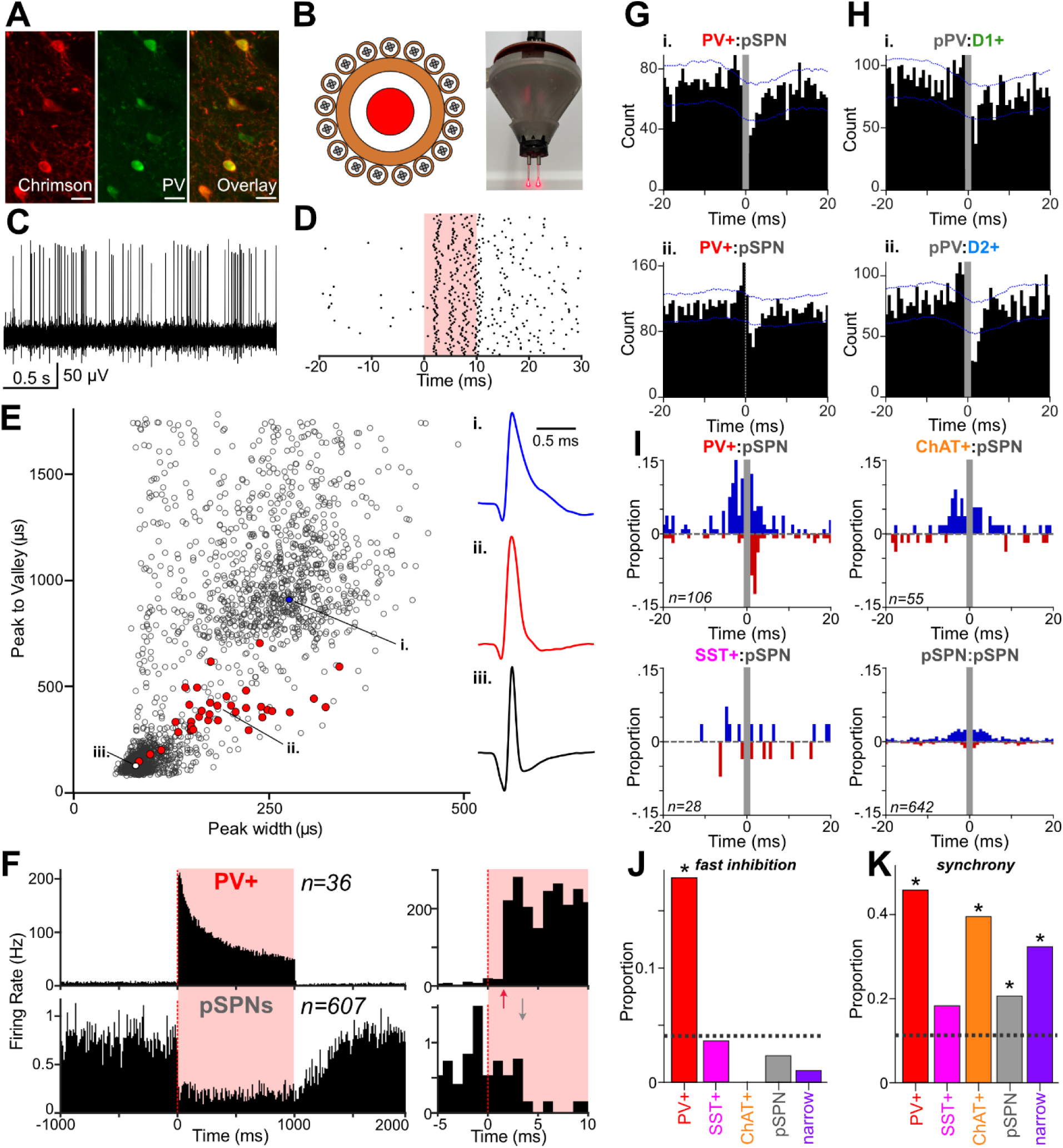
Striatal PV+ rapidly inhibit and synchronize with SPNs. **(A)** Images showing the colocalization (overlay image, right) of ChrimsonR-tdTomato expression (red, left) with PV+ staining (green, middle). Scale bars, 20 µm. **(B)** Optogenetic tagging approach. Left, schematic of tetrodes surrounding an optic fiber; right, complete assembly for bilateral implant. **(C)** Filtered signal from one tetrode wire showing spiking of a single PV+ unit. **(D)** Raster plot showing the response of a PV+ unit to 10 ms laser pulses (red). **(E)** Left, measurements of average spike waveform duration for all DLS units (n=1,880) recorded in PV-Cre rats with Cre-dependent ChrimsonR (n=4). Opto-tagged (PV+; n=36) are shown in red. Right, average waveforms of three example units (i, ii, iii). **(F)** Left, mean firing rates of PV+ (top, n=36) and pSPNs recorded in the same sessions (bottom, n=607) aligned on 1 s epochs of PV+ opto-stimulation. Data is shown in 1 ms bins. Right, close-up of the first few ms after laser onset. PV+ firing increase begins ∼2 ms after light onset, pSPN firing decrease begins ∼4 ms after laser onset. **(G)** Session-wide cross-correlograms for example PV:pSPN pairs recorded on the same tetrode (i) or a different tetrode (ii). Plots are aligned on the PV+ spikes. Bin duration = 0.75 ms. Grey bar around zero in (i) indicates epoch (-0.75 to +0.75 ms) when cross-correlogram is not calculated for neuron pairs on the same tetrode, as overlapping spikes may not be correctly sorted. Blue dotted lines indicate 99% confidence intervals. **(H)** Example cross-correlograms for a pPV:D1+ pair (i) and pPV :D2+ pair (ii). Same format as G. **(I)** Proportions of all cross-correlograms (constructed as in G) that reached significance - i.e. more (blue) or less (red) pSPN firing than expected by chance, for each 0.75ms time bin for pairs of neurons recorded from the same tetrode. Zero time is the firing of identified interneurons (PV+, SST+, ChAT+, top three rows) or other pSPNs (bottom row). **(J)** For each DLS unit subtype, the proportions of cross-correlograms with pSPNs showing a significant dip in the time range 0.75-3.75ms (consistent with fast inhibition). Dashed line shows proportion expected by chance (4 bins tested each with α=0.01, so chance ∼3.9%). Asterisks indicate proportion significantly exceeds chance (binomial test, PV+:pSPN p=1.11 ×10^−7^). **(K)** For each DLS unit subtype, the proportions of cross-correlograms with pSPNs showing a significant peak in the time range -5ms to +5ms (i.e. synchronous spiking). Dashed line shows proportion expected by chance (12 bins tested each with α=0.01, so chance ∼11.36%). Asterisks indicate that proportion significantly exceeds chance (binomial tests, PV+; pSPN pairs p=6.65×10^−14^, ChAT+:pSPN p=2.52×10^−7^; pSPN:pSPN 5.74×10^−11^; narrow:pSPN 8.97×10^−8^).

Across all electrophysiological data sets we consistently replicated prior observations that extracellularly recorded striatal units mostly fall into two clusters based on waveform duration (Fig. 1E; Fig. S1). As expected for SPNs, virtually all identified D1+ and D2+ units (D1+ 327/351, 93%; D2+ 389/408, 95%) fell into the largest cluster, and showed sporadic rather than continuous (tonic) firing (Fig. S1). Following prior assumptions, we further expected that identified PV+ neurons would fall squarely into the dense cluster with the narrowest waveforms. Instead, PV+ formed a largely distinct, rare cluster (Fig. 1E). All PV+ fired tonically in awake rats, though less regularly than identified ChAT+ (Fig. S1). SST+ did not consistently show tonic firing (Fig. S1) despite their cell-autonomous activity *ex vivo*(*28*).

### PV+ inhibit and synchronize with SPNs

With these positively identified PV+ we revisited the question of how PV+ influence SPNs in awake behaving animals. First, we confirmed that evoking PV+ spikes via optogenetic stimulation can very rapidly inhibit pSPN firing(*29,30*). In rats expressing ChrimsonR in PV+, one second illumination evoked strong firing of PV+ and a corresponding reduction in average pSPN firing (Fig. 1F, left). The onset of this reduction was ∼2ms after the observed increase in PV+ firing (Fig. 1F, right; red and grey arrows). Next, we examined the influence of spontaneous PV+ spikes, using session-wide cross-correlograms. PV+ spikes were often followed by reduced pSPN firing ∼2ms later (Fig. 1G, I). This short delay is consistent with monosynaptic fast inhibition(*12,31*), although our extracellular recordings cannot establish monosynaptic connections with certainty(*32*). This candidate fast inhibition was predominantly seen for PV+:pSPN pairs recorded from the same tetrode, that are likely within tens of µm from each other(*33*)(Fig. 1Gi, I). We did observe similar cross-correlogram dips for some pairs recorded from different tetrodes within the same hemisphere (Fig. 1Gii), but these interactions were rare (Fig. S2A). For identified SST+ and ChAT+ interneurons no comparable fast influence over pSPNs was apparent (Fig. 1I, J), although we did observe candidate examples of more prolonged SST+ inhibition of pSPNs (Fig. S2B). This overall pattern fits anatomical and physiological data from slices: while PV+ provide short-range perisomatic synapses onto SPNs, SST+ tend to make connections at longer distances (up to mm) (*7*) onto SPN distal dendrites(*31*) and thus have a less direct relationship to SPN spiking(*34*).

In slices, PV+ provide GABAergic feedforward inhibition to both D1+ and D2+ SPNs(*29*). While we did not have simultaneously tagged PV+ and D1+/D2+ recordings, we used PV+ properties newly identified in this study to estimate which untagged cells in our D1+ and D2+ recordings are likely PV+ (“pPV”; see Methods and Figs. S1,S3). We found clear examples of inhibition for both pPV:D1+ and pPV:D2+ pairs (Fig. 1Hi, ii), and these showed the same fast (∼2ms), brief timing as for PV+:pSPN pairs (Fig. S6B). We conclude that PV+ provide robust fast inhibition in behaving rats, helping determine the precise timing of both D1+ and D2+ SPN spikes.

The correlation analyses also revealed cell-type-specific, millisecond-timescale synchronization of nearby striatal neurons. PV+:pSPN pairs often (though not always) showed narrow cross-correlogram peaks (e.g. Fig. 1Gii). The timing of these peaks was variable, even when examining PV+:pSPN pairs from the same PV+ neuron (Fig. S2D). In aggregate, this variability produced a hump of positive synchrony in the approximate time range -5ms to +3ms (Fig. 1I). Plausibly, this aggregate pattern reflects common excitatory input to PV+ and nearby SPNs, together with feedforward inhibition reducing SPN spiking after PV+, resulting in an asymmetrical synchrony hump. Pairs of nearby pSPNs themselves, along with ChAT+:pSPN pairs, also showed synchrony on a similar millisecond timescale (Fig. 1I). The proportion of pSPN:pSPN pairs showing significant synchrony was relatively low (Fig. 1K), though this partly reflects the modest statistical power caused by the lower firing rates of SPNs. Interestingly, at longer distances (different striatal tetrodes, within the same hemisphere) SST+: pSPN pairs appeared to be anticorrelated on millisecond timescales (Fig. S2A) - a phenomenon that merits further investigation. Overall, striatal neurons show cell-type-specific patterns of millisecond-scale inhibition and synchrony during behavior, with PV+ having a distinct role providing fast inhibition to nearby SPNs.

### PV+ increase firing as rats wait

Rats were trained to perform a probabilistic reward (“bandit”) task that we have previously used to study striatal dopamine signals(*25,35,36*) and other neuronal populations(*27,37*). Within an operant chamber, freely moving rats poked and maintained their noses in a central port, waited for an auditory *Go!* cue, then made a brief lateral movement either leftward or rightward to an adjacent side port (Fig. 2A, top). Side port entry triggered sucrose pellet reward delivery, with varying probability (see Methods). We focused our analyses on the periods of waiting (the “hold” period before the *Go!* cue), action initiation (*Go!* cue to Center Out) and action execution (Center Out to Side In).

**Fig. 2.**
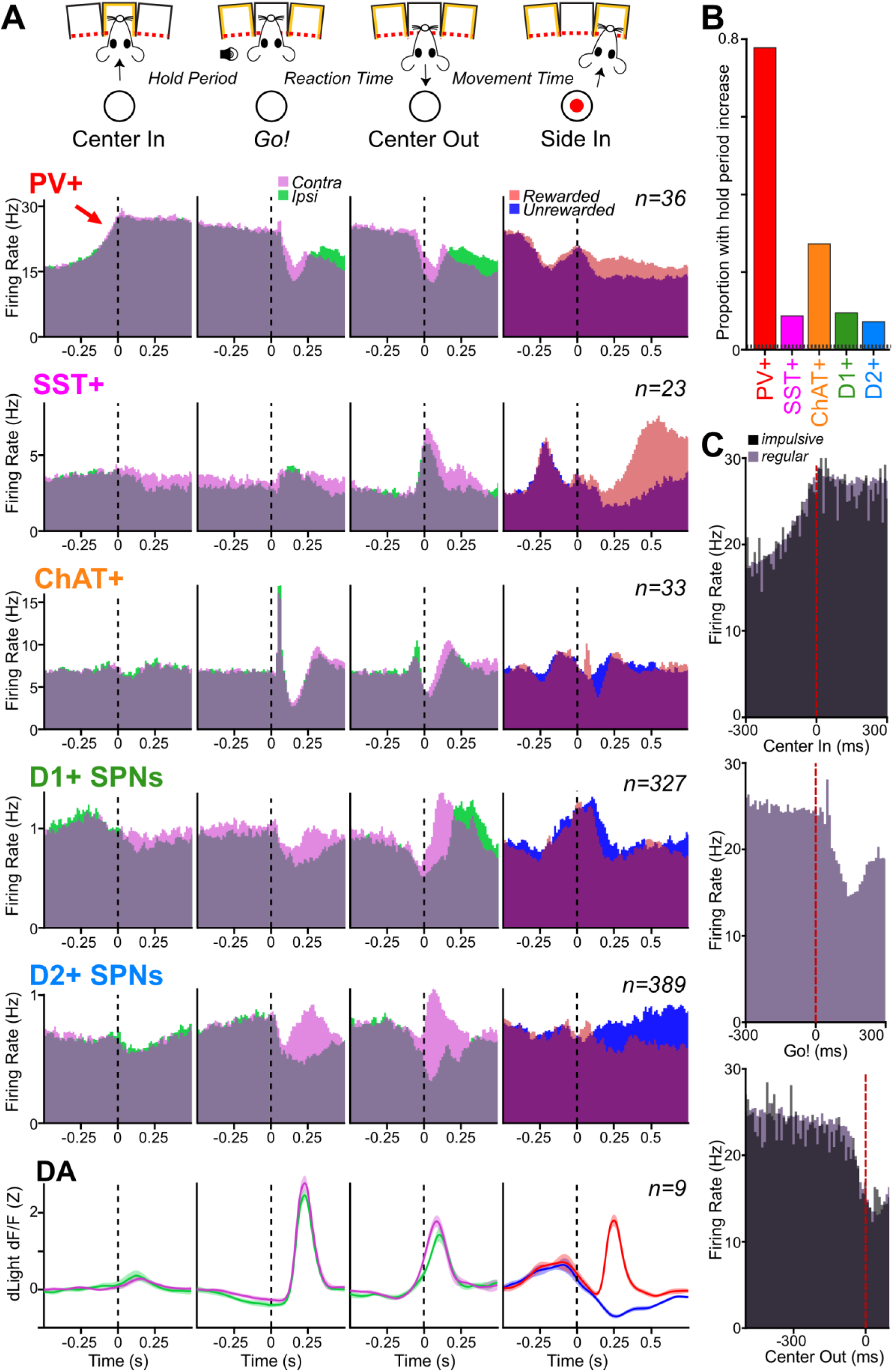
Striatal PV+ increase firing as rats wait to move. **(A)** Top, schematic of selected behavioral task events; middle rows, corresponding mean peri-event firing rates for neurons of each identified type; bottom, event-aligned dopamine signals (fiber photometry for dLight1.3b, +-SEM; recorded in a separate set of rats^29^). For Center In, *Go!* Cue, and Center Out alignments, trials are divided *Contra / Ipsi* (movement away from / towards the recorded hemisphere; magenta/green). For Side In alignment, trials are divided into *Rewarded / Unrewarded* (red / blue). **(B)** Proportion of identified cells of each type showing significantly elevated firing during the hold period (-500 to 0 ms relative to *Go!* cue) compared to shuffle controls (10,000 shuffles, one-tailed test, α=0.001, dashed line). Each of the tagged subpopulations significantly exceeded chance (binomial test, p<0.05). PV+ modulation was different from all other populations; ChAT+ cells were significantly different from D1+ and D2+; no other comparisons were significant (q<0.05, FDR-corrected Fisher’s tests). **(C)** Comparison of mean peri-event firing rates of PV+ on regular (mauve) and impulsive (black) trials for Center In, *Go!* Cue, and Center Out alignments.

On average, each DLS interneuron class showed activity around behavioral events that was very distinct to other interneurons, to D1+ and D2+ SPN firing, and to DLS dopamine release dynamics(*26, 36*)(Figs. 2A, S3). In particular, DLS PV+ increased firing as rats entered the center port (Fig. 2A, red arrow), and this elevated mean level was maintained during the hold period. This increase was quite consistent for individual PV+ neurons (28/36, 78%; see Appendix for details on each individual identified PV+ neuron) but not for other cell types (Fig. 1B). The *Go!* cue was followed in quick succession by a burst-pause in DLS ChAT+ firing(*26*), a sharp drop in mean PV+ firing, a pulse of dopamine release, and a movement-locked increase in mean SST+ firing (Fig. 2A).

The elevated firing of DLS PV+ as rats hold their position suggests a role preparing and/or restraining actions. To assess this, we first examined “impulsive” trials in which rats failed to wait until the *Go!* cue (12.8% of trial starts in PV+ recording sessions). In regular trials, PV+ reduced their collective activity starting around ∼65ms after the *Go!* cue and preceding Center Out (Fig. 2C). In impulsive trials, despite the absence of the *Go!* cue PV+ activity also dropped sharply before Center Out (Fig. 2C), consistent with a restraint role.

### PV+ suppression immediately impacts action restraint, selection and execution

To further test PV+ functions we turned to precisely timed optogenetic suppression. We again infused virus for Cre-dependent opsin expression in the DLS of PV-Cre rats (n=11), this time using the inhibitory opsin stGtACR1(*38*)(Fig. 3A). In a subset of rats (n=2) we combined this opsin expression with electrophysiology to verify effective inhibition of PV+. Short epochs of green laser light (300ms, 505nm, 1mW) strongly and rapidly suppressed the activity of a cell subpopulation (Fig. 3B). As expected, the opto-suppressed cells had waveforms, firing statistics and behavioral correlates that closely resembled the PV+ identified through excitatory optogenetic tagging (not shown). Within the bandit task we first provided this optogenetic suppression of PV+ bilaterally during the hold period (for 500ms, starting 250ms after Center In). This increased the rate of impulsive trials (Fig. 3C), again supporting a role for PV+ in action restraint.

**Fig. 3.**
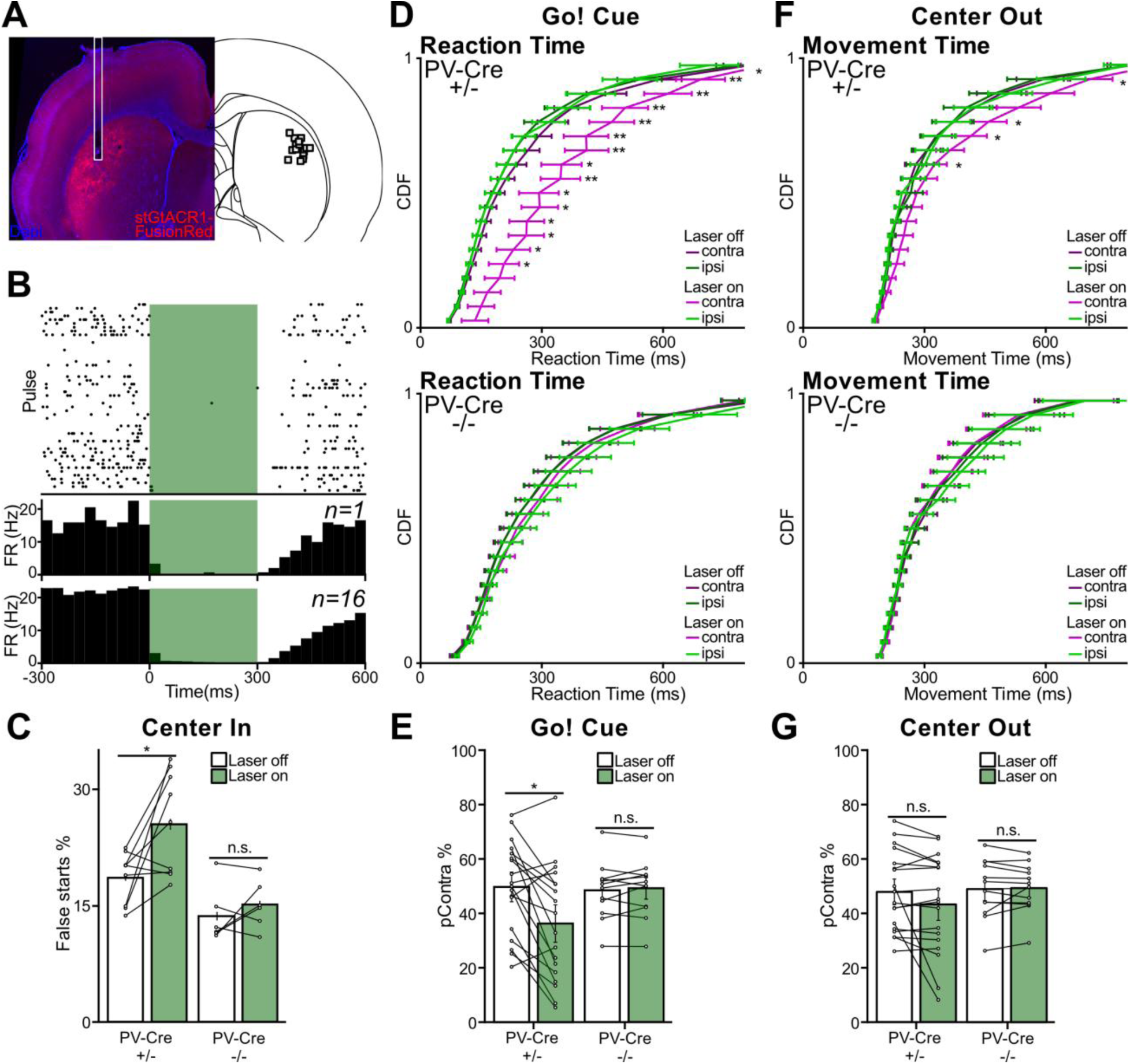
Suppression of PV+ increases premature actions, delays cue-evoked actions. **(A)** Left, example of stGtACR1-FusionRed expression in DLS, with white outline indicating optic fiber position. Right, range of tip positions for all fibers in DLS (shown on the same side for simplicity). **(B)** Spike raster (top), histogram (middle) for one example unit and average histogram (bottom) for all recorded neurons rapidly suppressed by illumination (300ms, 505 nm, 1mW at fiber tip). **(C)** Effect of bilateral illumination (500ms, starting at Center In) on the rate of premature action initiation (false starts), in Cre-positive (hemizygous, +/-, n=9) and negative (-/-, n=6) rats from the same PV-Cre colony. Error bars, SEM. Wilcoxon rank-sum tests, two tailed: Cre-positive p=0.039, Cre-negative p=0.4375. **(D)** Cumulative density functions (CDF) of reaction times, with and without unilateral illumination of DLS (300ms, starting at *Go!* cue). Top panel, Cre-positive rats; lower panel, Cre-negatives. Suppressing PV+ via stGtACR1 selectively slowed movement initiation for *Contra* trials (magenta), in Cre+ but not Cre-animals. Asterisks show significant difference for bins within-rats (one-tailed Wilcoxon signed-rank test at each quantile, with FDR correction q < 0.05. Considering each DLS in each individual rat, the *Contra* reaction time CDF showed a significant change (Laser ON vs OFF, KS test p<0.05) in 8/18 Cre+, but 0/12 Cre-hemispheres. **(E)** PV+ suppression at *Go!* cue biases choice away from *Contra.* Wilcoxon rank-sum tests: p=0.0038 for Cre+ and p=0.4697 for Cre-. **(F)** Effects of PV+ suppression at Center Out on movement times (same format as D). Movement execution was selectively slower for *Contra* trials (magenta). Among individual hemispheres tested, 6/18 Cre+, but 0/12 Cre-hemispheres showed a significant change in movement time distribution for *Contra* trials (KS test, p<0.05). **(G)** Same as F, except showing that laser illumination at Center Out has no significant effect on *Contra / Ipsi* choice.

To assess whether the functional contribution of PV+ is limited to withholding movement, we also suppressed PV+ later in the trial: for 300ms beginning at either the time of *Go!* cue, or at Center Out, once movement has already begun. Unilateral PV+ suppression slowed reaction times (movement initiation) selectively for contraversive actions (“*Contra*”; i.e. those made in the direction away from the manipulated DLS hemisphere; Fig. 3D). This manipulation also introduced a directional bias, making *Contra* choices less likely than ipsiversive movements (“*Ipsi*”; Fig. 3E). Unilateral PV+ suppression at Center Out selectively slowed ongoing *Contra* movement execution (Fig. 3F) without affecting choice (Fig. 3G). No optogenetic effects on behavior were seen in control rats from the same colony, that lack Cre recombinase (Fig. 3C-G).

### PV+ participate in fast movement sequences

To determine how PV+ suppression may interfere with movement performance, we looked more closely at the firing of individual identified neurons. Across a range of behaviors striatal neurons have been reported to fire in sequential order(*20,39,40*). Similarly, in our task subsets of both D1+ and D2+ SPNs fired in very rapid sequences (Fig. 4A, B). These sequences were apparent in simultaneously recorded populations of tagged and untagged striatal neurons, and distinguished between *Contra* and *Ipsi* actions (Fig. S4, 5). Sequences were predominantly locked to the movements themselves (Center Out), rather than the *Go!* cue that evoked the movement (Fig. S5). For each neuron we measured the time of peak firing within the movement sequence (“peak time”) and the duration of this firing (“peak width”, measured half-way between zero and the maximum rate). Peak widths varied between neurons, but were often very brief (example, Fig. 4A, purple) indicating precise firing at a specific part of the sequence. Peak times were more similar for pairs of neurons recorded from the same tetrode, compared to more separated pairs (Fig. 4D). In other words, nearby neurons are more likely to jointly participate in a functional cell assembly, that forms and disappears as the sequence progresses.

**Fig. 4.**
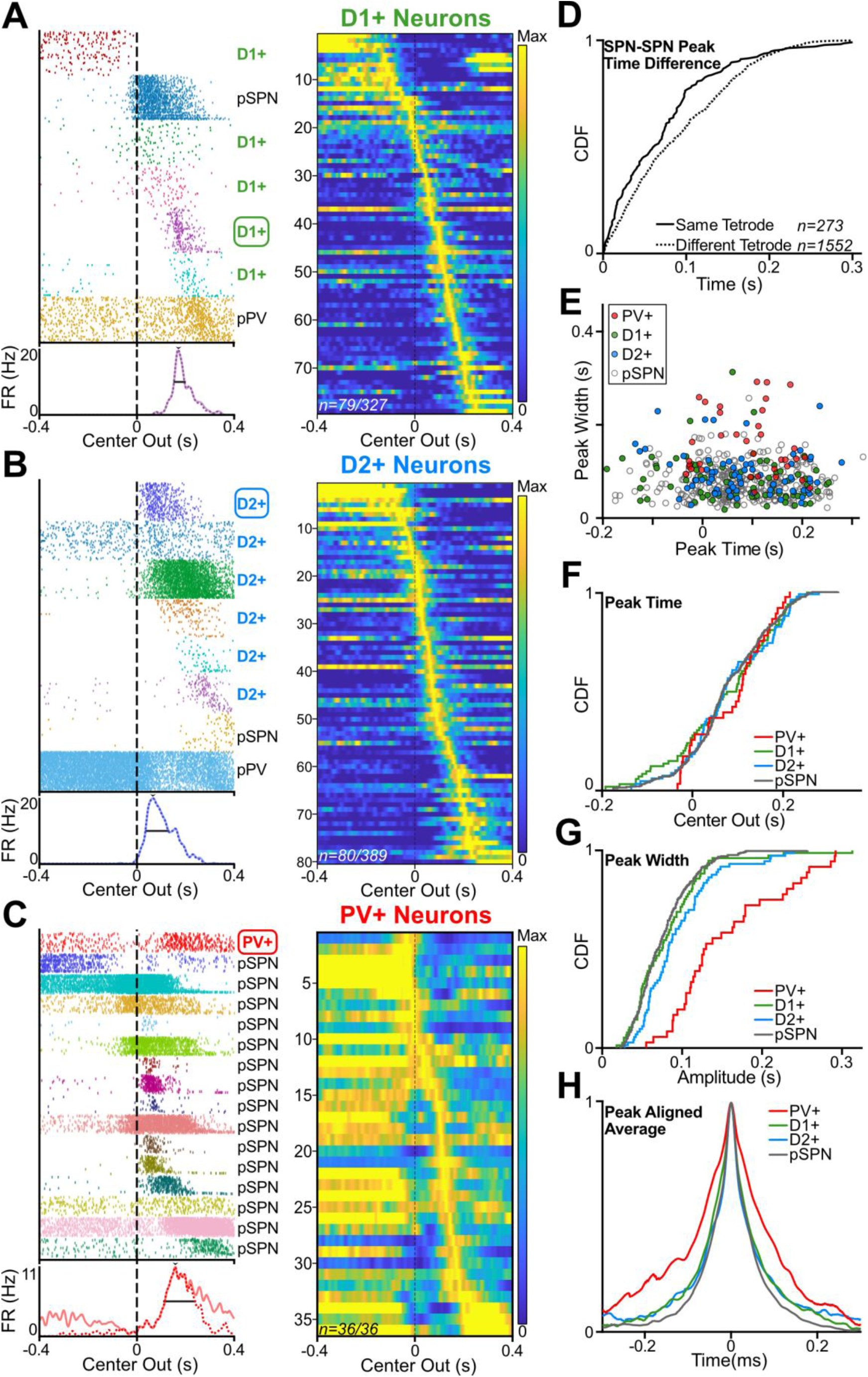
Joint firing sequences of D1+, D2+ and PV+ during actions. **(A)** Left, example of a *Contra* sequence from a single recording session, showing spike rasters for simultaneously recorded D1+ and pSPNs. For each unit, trials are ordered by movement time. For clarity of individual spikes, only 50 *Contra* trials are shown for each cell (from 183 *Contra /* 122 *Ipsi* trials in this session). Bottom, peri-event histogram for the indicated neuron (circled, purple) from the sequence. Horizontal lines mark peak width (at half-maximum; note peak width is overestimated by not compensating for variable movement duration). The histogram is plotted twice, without (solid) and with (dotted) removal of spikes that fell outside the designated epoch for movement-related activity (70 ms after *Go!* cue until Side-In). Right, activity of D1+ SPNs during *Contra* movements, sorted by the time of their peak firing rate. Included are the subpopulation (79/327, 24.2%) that reached a minimum firing rate of >3Hz, at some point from 200ms before Center Out until Side In. **(B)** As A, but for D2+ SPNs (80/389, 20.6%). **(C)** As A,B but for PV+ (all 36 tagged cells met the >3Hz inclusion criterion). **(D)** Cumulative density plot of the difference in *Contra* peak time for pairs of neurons, either recorded on the same tetrode (solid, n=273) or different tetrodes within the same hemisphere (dotted, n=15,524). Included are all D1+, D2+, and presumed SPNs that met the 3Hz criterion. Data were not divided by SPN subtype because relatively few pairs on the same tetrode included identified SPNs. **(E)** Scatter plot comparing *Contra* peak times and half-width durations for PV+, D1+, D2+ and pSPNs. **(F)** Cumulative density plots of *Contra* peak times. There were no significant differences between subpopulations (pairwise KS tests, all p > 0.83 after Bonferroni correction for multiple comparisons). **(G)** Cumulative density plots of *Contra* peak widths. PV+ had significantly different widths to each SPN subpopulation (pairwise KS tests and Bonferroni corrected for multiple comparisons PV+ to D1+ p=7.11×10^−6^; PV+ to D2+ p=9.15×10^−4^; PV+ to pSPNs p=6.97×10^−10^) but SPNs subpopulations did not differ (all p >0.79). **(H)** Averaged, normalized, peak-aligned *Contra* firing rates, showing broader average peaks of PV+ compared to SPNs.

Unexpectedly, PV+ also participated in these fast action sequences (Fig. 4C). As with SPNs each PV+ reached peak firing at a distinct moment, and the firing times of SPNs and PV+ neurons all tiled the full movement duration (Fig. 4E, F). However each individual PV+ was active for a longer duration compared to SPNs (median peak widths: D1+, 71ms; D2+, 83ms; pSPNs, 68ms; and PV+, 130ms; Fig. 4G,H). Furthermore, the great majority of individual PV+ participated at some point during each movement sequence, compared to only a minority of SPNs (Fig. 4A-C, S5, S6). The greater participation and more extended firing of PV+ compared to SPNs are consistent with prior evidence that PV+ receive a broader range of cortical inputs, and are especially sensitive to these inputs(*41,42*).

### PV+ influence which SPNs participate in sequences

To assess how this sequential PV+ activity contributes to ongoing sequential SPN activity, in another set of PV-Cre rats (n=4) we placed bilateral optrodes in DLS, with unilateral infusion of virus for Cre-dependent expression of the inhibitory opsin HcKCR1(*43*). During task performance we delivered green light (514 nm, 2mW, 300ms) starting at Center Out, to either the virus side or the opposite control side (in separate sessions, in 30% of trials). Identified PV+ neurons (see methods) were effectively and rapidly inhibited by light during task performance (Fig. 5Ai, B).

**Fig. 5.**
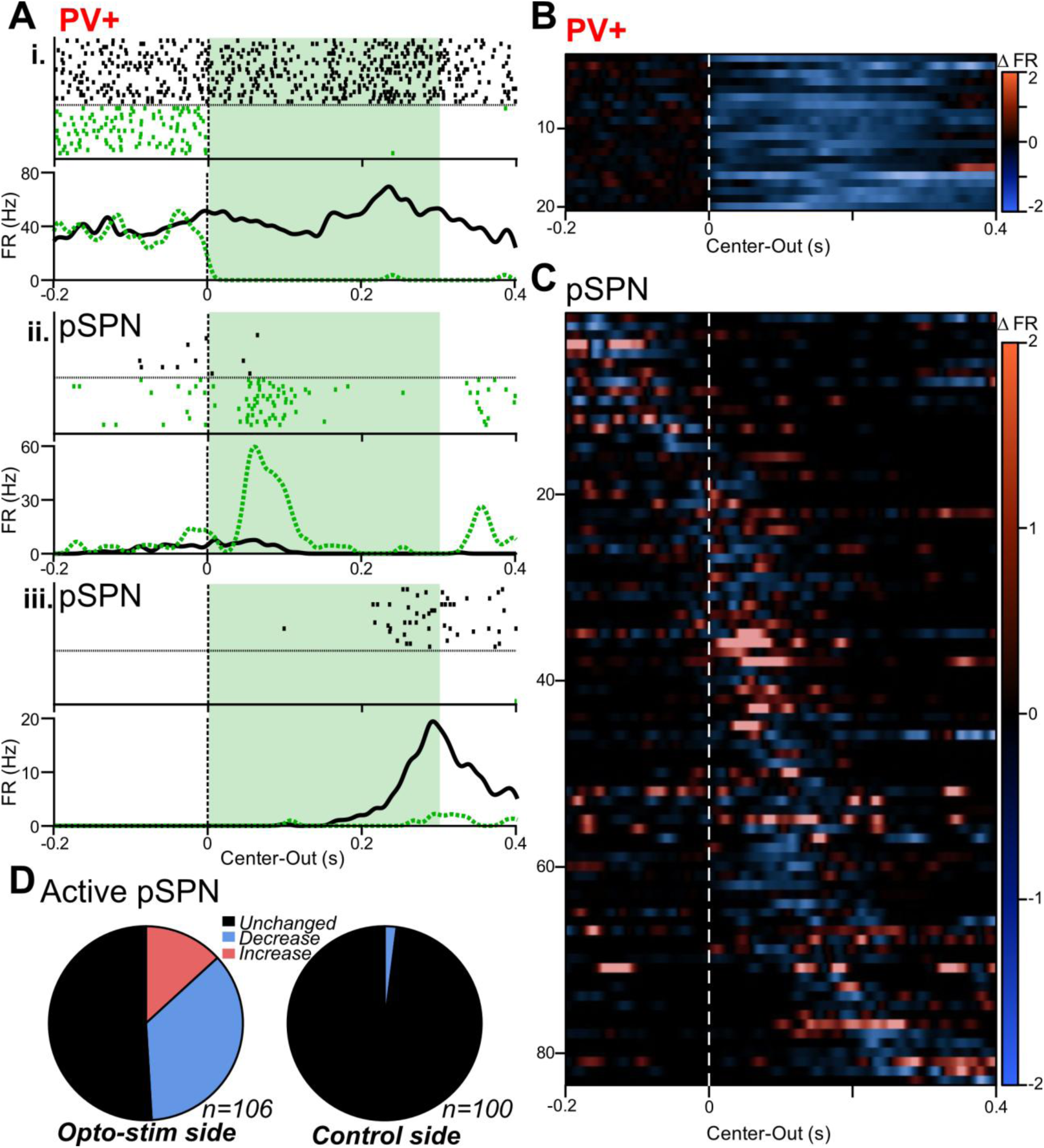
Striatal PV+ shape SPN movement sequences. **(A)** Spike raster and PETH of example PV+ (i) and pSPNs (ii,iii), aligned to Center Out in control (black) and laser ON (green) trials. Examples are taken from a PV-Cre rat with Cre-dependent expression of HcKCR1 in DLS. Illumination was provided on a randomly selected 30% of the trials (300 ms, 2mW at the fiber tip). **(B)** Change in normalized firing rate of all inhibition-identified PV+ neurons aligned to Center Out. **(C)** Change in normalized firing rate of pSPNs active during the laser-activation period (n=106/486 pSPNs for virus-infused sides and n=100/375 for control sides). **(D)** Proportions of active pSPNs with unchanged (gray), decreased (blue) and increased (red) firing rates during the laser period, comparing on the virus-infused (left) and control (right) sides. Significant changes in pSPN firing rate were determined using trial-based shuffle tests (10,000 shuffles, one sided α=0.01). On the virus infused side, the proportion of cells with increased and decreased FR were significantly different from chance (two-sided exact binomial tests; increased: p<1×10^−6^; decreased: p<1×10^−6^). For the control side, neither proportion was significant (increases: p=1; decreased: p=0.264). The proportion of cells with increased or decreased FR was also different between virus-infused and control sides (two-sided Fisher exact tests; increased: p=8.3×10^−5^; decreased: p<10^−6^).

Loss of PV+ firing during movement execution had strong but selective effects on pSPN firing. Few if any pSPNs showed sustained enhancement of firing during the 300ms illumination period (Fig. 5C; S7). Instead, some cells that were quiescent in regular trials acquired reliable, transient movement-related activity (example, Fig. 5Aii), while others with clear movement-related firing in regular trials stopped firing (example, Fig. 5Aiii). Light-evoked decreases in pSPN firing were more common (37/106) than increases (14/106), and there were no significant changes on the control side (Fig. 5D; S7). Overall, SPNs continued to fire in sequences during PV+ suppression, but with altered SPN composition, demonstrating the critical role of PV+ in shaping ongoing striatal output.

## Discussion

Identifying striatal neurons in behaving animals has been a long-sought goal(*44*). In the absence of positive identification reasonable assumptions have been made based on properties including waveform and firing statistics. In the case of ChAT+ interneurons these assumptions - long waveforms and regular tonic firing - have turned out to be largely, though not consistently, accurate(*26*). Here we found that for the PV+ fast-spiking interneurons such assumptions are not safe. As expected, PV+ indeed have briefer extracellular waveforms than SPNs(*8,14,45*)(Fig. S1), and fire at higher rates. Yet PV+ do not map well onto the prominent cluster of particularly narrow waveform units consistently seen across many studies. Although we did not positively identify these common narrow units, they do not correspond to any of the five cell types we recorded that together comprise ∼99% of rodent striatal neurons(*46*). They most likely correspond to a population of axons, as has been recently reported for the narrowest units in cortex(*47*). One specific possibility is axons from globus pallidus, particularly the “Arkypallidal” neurons that project heavily to striatum(*48*). On average, the firing patterns of striatal narrow units resemble those we reported for Arkypallidal cells in the same behavioral task(*37*), although both narrows and Arkypallidals show very diverse activity. Overall, prior studies examining striatal presumed FSIs likely sampled from a variable mixture of actual PV+, other GABAergic interneurons, and axons, depending on the details of recording conditions and classification criteria.

To a first approximation the striatum is an inhibitory network(*49*). SPNs receive GABAergic inputs both from interneurons and from each other(*50*), and throughout striatum this local GABAergic circuitry is important for restraining mistimed or maladaptive behaviors. Interfering with ventral striatal PV+ increases impulsive approaches(*51*), while interfering with PV+ in dorsal-medial striatum increases compulsive grooming(*52*). In sensorimotor striatum, blockade of glutamatergic inputs to PV+ can evoke dyskinesias(*53*) and interfering with GABA-A transmission results in tic-like jerks(*54*). Such abnormal movements are blocked by local infusion of glutamate antagonists(*55*), suggesting they result from local microcircuits misprocessing cortical and/or thalamic inputs(*56*). However, another study that selectively interfered with DLS PV+ found no sign of altered movements, and proposed that these cells are more involved in governing plasticity than online movement control(*29*).

The ongoing need for DLS PV+ may be particularly apparent when performing highly trained, kinematically refined movements. After thousands of trials of bandit task training our rats perform the *Contra* and *Ipsi* movements very quickly, and momentary suppression of DLS PV+ immediately interfered with this expert performance. This fits with prior results that blockade of DLS GABA-A receptors (or DLS activity more generally) causes previously trained actions to become slower and more variable(*54, 57*), and that chemogenetic suppression of DLS PV+ causes behavioral control to switch out of a well-trained, habitual mode(*58*).

Compared to other striatal subregions the DLS is enriched in PV+ cells(*59*), likely reflecting a particular DLS need to produce temporally precise SPN sequences. While sequential activity has been reported in multiple subregions(*20,57,60*), the DLS shows a quicker tempo of neural activity(*20,36,61*), and the fast SPN sequences we observed during brief (∼200-300ms) movements are similar to those reported for pSPNs during highly practiced brief birdsong syllables(*62*). Striatal sequences have been argued to encode the passage of time(*39*), serving as a temporal framework for reinforcement learning(*63,64*). Ensuring that each DLS SPN fires only briefly at a specific moment within sequences may therefore be important to support kinematics that are precise and repeatable over short timescales.

Firing of both PV+ and SPNs requires external excitatory afferents(*7,65*), and DLS PV+ respond more strongly and consistently to these inputs compared to nearby SPNs(*29*), or to SST+ or ChAT+(*40*). An SPN that receives synchronous glutamatergic inputs may respond quickly enough to spike, but fast feedforward inhibition via perisomatic PV+ synapses can veto SPN responses to weaker or less temporally organized input(*14,66*). In this way PV+ may help enforce coincidence detection(*67*) so that SPNs are sensitive to highly specific cortical and/or thalamic states. PV+ mediated inhibition is brief and subject to short-term depression(*68, 69*), so does not necessarily prevent those SPNs that receive the “right” input from increasing their firing rate and forming functional ensembles(*16,70*).

We found that PV+ spikes are frequently followed by reduced spiking of nearby SPNs ∼2ms later, but PV+ have a more complex influence over SPNs than simple inhibition(*69*). The chloride conductance associated with GABA-A input can also facilitate SPN spiking, in particular when it just precedes excitatory input(*71,72*). As a result, in some large-scale striatal simulations removing PV+ actually raises overall SPN firing rates(*70*). The ability of PV+ input to move SPNs closer to firing threshold may help explain why suppressing PV+ activity caused more SPNs to lose, rather than gain, movement-related activity.

An unanticipated feature of our data was millisecond-precision synchrony between nearby striatal neurons, including both SPN:SPN pairs and PV+:SPN pairs recorded from the same tetrode. Measured on slower time scales using calcium imaging, SPNs tend to be co-active within local zones at a spatial scale of ∼100µm or less(*73–75*). Our results indicate that more precise synchrony of striatal neurons is also a local phenomenon, which may account for why prior studies have typically reported that fine timescale synchrony in striatum is very rare(*18*) or absent(*76*). Synchrony can reflect common excitatory input, and individual cortical axons synapse onto both PV+ and nearby non-PV+ neurons(*77*). However, cortical axons often extend across large (mm-scale) zones of striatum(*78*). The local spatial scale of SPN synchrony may therefore be driven by the relatively compact and dense axon field of PV+ neurons. Multiple active SPNs under the influence of the same PV+ are more likely to fire together, as the inhibitory current wanes(*71,79*). Consistent with this, one prior investigation did report correlated firing between (unidentified) striatal cells(*80*), and found that this correlation was reduced in mouse models of Huntington’s Disease which have a deficit in PV+ interneurons(*81*). Synchrony between SPNs may boost the impact of their convergent projections to downstream basal ganglia structures, which consist of tonically active neurons sensitive to the fine timing of inputs(*82*). Dopamine loss increases PV+ connectivity to D2+ SPNs, and in simulations this enhances D2+ SPN synchrony(*83*), potentially driving aberrant network dynamics and motor symptoms in Parkinson’s Disease.

For investigating local circuit mechanisms our *in vivo* extracellular recordings and analyses have notable limitations. For example, reaching significance in short-timescale cross-correlations requires considerable numbers of spikes, and most SPNs fire at low rates. As a result, we very likely underestimate the overall incidence of fast intrastriatal interactions. We also cannot easily test how these interactions evolve during behavior(*32*), given how briefly DLS SPNs typically fire during movements. Striatal information processing is further influenced by synapses between interneurons(*84*), but we obtained too few pairs of simultaneously recorded, identified interneurons to adequately assess such interactions. Nonetheless, our demonstration that identified PV+ increase firing during action restraint and shape striatal output is a substantial step towards understanding why, in clinical syndromes, loss of PV+ can release inappropriate actions, and why those actions should have an aberrant character.

## Acknowledgments

We thank Charles Wilson, Massimo Scanziani, Vikaas Sohal, Vijay Mohan K Namboodiri, Howard Fields, Ali Mohebi, Dennis Burke and members of the Berke Lab for comments on earlier versions of our results, and Nasim Elyasi for technical support.

## Author contributions

Conceptualization: J.B. and M.D. Data Curation: M.D., I.G.M., C.Z., L.P. Formal Analysis: M.D. Funding Acquisition: J.B., M.D. Investigation: M.D., I.G.M. Software: M.D. Supervision: J.B. Visualization: M.D., I.G.M. Writing – original draft: J.B. and M.D. Writing - Review and Editing: J.B. and M.D.

## Data, code, and materials availability

Data set including spike times, and behavior, for identified neurons together with analysis code, are available at: https://doi.org/10.5281/zenodo.22134941(85).

## Materials and Methods

### Animals

All animal procedures were approved by the University of California San Francisco Institutional Committee on the Use and Care of Animals. We maintained colonies of rats of five distinct genotypes: D1-Cre and A2a-Cre(*88*), PV-Cre(*89*), SST-Cre (LE-SSt.*Cre^em1sage^*, Inotiv) and ChAT-Cre(*90*), all generated on a Long Evans background and maintained by backcrossing to wild type Long-Evans rats (Charles River). Male (400-550 g) and female (300-400 g) rats, 6-12 months old on a reverse 12:12 light:dark cycle were tested during the dark phase. Rats were mildly food deprived, receiving 4g / 100g of body weight of standard laboratory chow in addition to food pellets received during task performance, and body weight was frequently monitored to stay between 85-90% of baseline. No sample size pre-calculations were performed.

### Behavior

Bandit task pretraining and testing were performed in computer-controlled Med Associates operant chambers (25 cm × 30 cm at widest point) each with a five-hole nose-poke wall, as previously described(*25,26,35,36*). The hold period before *Go!* cue was 500–1500 ms (uniform distribution). Left/right reward probabilities were each 10, 50, or 90%, held constant for block lengths of 35–45 trials and independent and randomly selected for each block. Rats were trained to complete >75% of trials without procedural errors before implant surgery.

### Optogenetic Tagging

43 rats (D1-Cre n=13; A2a-Cre n=8; PV-Cre n=6; SST-Cre n=8; and ChAT-Cre n=8) were infused with 1μl of AAV5-Syn-FLEX-rc[ChrimsonR-tdTomato] (Addgene, 62723-AAV5) into DLS (AP 0, ML ±4.0, DV 4.0mm). During the same surgery we implanted custom designed assemblies, consisting of 2 sets of 16 tetrodes each (constructed from 12.5 μm nichrome wire, Sandvik) inserted into a polyimide tube which slides around a 200 μm tapered optic fiber (Doric Lenses). Some assemblies targeted DLS bilaterally others DLS and DMS (data not shown). Only male rats were used for electrophysiological recordings due to the large size of the recording drive. The tetrodes were initially placed 500 μm above the fiber tip. Additionally, 5 bone screws (Fine Science Tools, catalog # 19010, blunted) were placed in contact with the brain surface. Two of them recorded frontal ECoG from each hemisphere (AP 5, ML ±2 mm), one was placed 1 mm posterior to bregma to serve as reference, and two were placed in the posterior lateral skull to serve as ground. During recording sessions, wideband (1–9000 Hz) brain signals were sampled (30,000 samples/s) using a custom headstage with 2 x 64-channel Intan RHD2164 digital amplifier chips(*37*). After bandit task performance, rats were then left for 40-60 min in the recording chamber to record a sleep session, with white noise constantly playing at 45 dB. Finally, optic fibers were connected for light delivery and the laser stimulation session was recorded. Which included 1 s pulses at 2mW measured at the fiber tip to test PV+ mediated inhibition of pSPNs. Tetrodes were lowered by at least 80 μm between sessions to avoid repeated recording from the same units, up to 500 μm below the fiber tip or until no light responsive units were detected over multiple sessions. 6 SST-Cre, 6 D1-Cre and 9 A2a-Cre rats in which we did not record any identified neurons were not analyzed further. The ChAT+ optotagging data and dopamine dLight1.3b photometry data have been previously published(*26*), and the photometry methods have been previously described in detail(*36*). In brief, dLight1.3b fluorescence at 470nm and 405nm (alternating every 10ms) was obtained through 200µm core fibers placed in DLS (AP 0, ML 4.0, DV 4.0mm), and the two signals were rescaled to each other via a least-square fit; signals shown are the averaged, Z-scored fractional fluorescence (dF/F) defined as the (470 - fit_405) / 405_fit.

### Optogenetic Inhibition

PV-Cre +/- rats (n=5 males, n=4 females) and PV-Cre -/- littermates (n=3 males, n=3 females) were bilaterally infused with 1μl of AAV5-hSyn1-SIO-stGtACR1-FusionRed (Addgene, 105678-AAV5) into DLS (males: AP 0, ML ±3.8, DV 4.2mm, females: AP 0, ML ±3.6, DV 4.0mm). During the same surgery we implanted an optic fiber in each hemisphere (males: AP 0, ML ±4.0, DV 4.0 mm, females: AP 0, ML ±3.8, DV 3.8 mm) for light delivery (200 µm core, 0.5 NA; Newdoon). 3 weeks were allowed for expression before any testing. During behavior light was delivered through split patch cables coupled to a 505 nm LED (Thorlabs) or a 505nm Laser Diode (Thorlabs) via a rotary joint. Light output was set to 1 mW at the fiber tip. Inhibition trials (30 %) were randomly interleaved with control trials (no illumination). Each session involved illumination beginning at one event: either Center In (bilateral, 500 ms), *Go!* cue (unilateral, 300 ms) or Center Out (unilateral, 300 ms). For each combination of rat, condition and hemisphere at least three sessions were acquired and data pooled. The session type was randomized before start. Light onset and pulse duration were controlled by the same LabView program used to control the operant chambers using an Arroyo Instruments laser controller as before(*35*). After all sessions were recorded, brains were extracted and processed to assess placement and expression; 11 additional PV-Cre +/- rats were excluded due to no viral expression detected. Reaction times (RT) and movement times (MT) across animals were compared using Vincentized averages to preserve within-subject distribution(*91*). The RT or MT of individual animals were sorted and binned into 20 equal sized quantile bins. Paired Wilcoxon signed-rank tests (right-tailed) compared Vincentized RT and MT between conditions at each quantile. P-values were corrected for multiple comparisons across bins using the Benjamini–Hochberg false discovery rate (q < 0.05). To test the effect of light on PV+ neurons we injected the virus under the same conditions but implanted a combined recording/optogenetic drive as described above (n=2).

To record the effect of PV+ inhibition on SPN firing during movement PV-Cre +/- rats (n=4, all males) were unilaterally infused with 1μl of AAV5-hSyn1-DIO-HcKCR1- mCherry (Vector Builder, HcKCR1-mCherry sequence from Addgene plasmid 177336) (*43*) into DLS (AP 0, ML ±3.8, DV 4.2mm). During the same surgery we implanted a drive for recording, same as for tagging. 3 weeks were allowed for expression before any testing. During behavior light was delivered through split patch cables coupled to a 514 nm LED (Thorlabs) via rotary joint. Light output was set to 2 mW at the fiber tip. Inhibition trials (30 %) were randomly interleaved with control trials (no illumination). Each session involved illumination beginning at Center Out (unilateral, 300 ms). For each position two sessions were recorded alternating the virus injected and control hemispheres.

### Histology

To confirm expression of ChrimsonR, FusionRed and mCherry in PV+ neurons we performed immunohistological staining. After recordings complete animals were anesthetized with isoflurane, then perfused with PBS 1X (Sigma-Aldrich P4417) solution followed by formaldehyde solution 4% (Sigma-Aldrich, F8775). 50-100 µm slices were blocked using a PBS solution containing 5% normal goat serum and 0.4% Triton x-100 (Sigma-Aldrich, 93443), then incubated with primary guinea pig anti-PV (AB_572259, Immunostar 1:1000), and mouse anti CD11b (AB_322901, BioRad 1:1000) antibodies followed by secondary goat anti guinea pig antibody conjugated with Alexa fluor 488 (AB_2534117, Invitrogen, 1:500) and goat anti-mouse antibody conjugated with Alexa Fluor 647 (AB_2866490, Invitrogen, 1:500). To determine recording and stimulation placement, slices with tracks were imaged using a Keyence BZ-X800 microscope with a 2 x objective, images were also assessed for tdTomato, Fusion Red or mCherry expression. We found expression of the fluorescent proteins to be 96.7% specific to PV+ immuno labeled cells and the efficiency of the targeting was 80.8% (n=426 cells from n=10 animals).

### Classification

All electrophysiological data analyses were performed in MATLAB (Mathworks, Inc.; Natick, MA). Spikes were sorted into individual units using the MountainSort algorithm(*92*) followed by careful manual inspection. To determine if units were light responsive, we evaluated their response to light pulse trains delivered at the end of the session(*93*). Trains of different widths: 2, 5 and 10 ms and frequencies: 1, 2, 5 and 10 Hz were used. For a unit to be identified as a light responsive PV+, ChAT+ or SST+, it needed to fire with a spike latency after laser stimulation significantly shorter (Wilcoxon rank sum test, p < 0.01) than the spike latency following randomly selected times within the same session. We also required that spikes appear within 10 ms of laser onset, in at least 30% of trials, for at least one stimulation condition(*26,37*). Due to the low firing rate of D1+ and D2+ cells, to determine their light responsiveness we instead used the stimulus associated latency test (based on Jensen-Shannon divergence) (*93*). To address the possibility that laser-evoked spikes were fired by a different neuron and included with the analyzed cell due to a spike sorting error, we also required that the waveforms of spikes occurring <10 ms following laser stimulation have a Pearson correlation coefficient >0.9 compared to the average waveform during bandit task performance^37^.Waveform and firing features were also used to classify neuronal subpopulations. Peak width was defined as the full-width-at-half-maximum of the most prominent negative-voltage component of the averaged spike waveform. Peak-to-valley time was the interval between the time of this peak and the time of the most positive voltage after this peak, within the 2 ms total duration of the spike analysis window.

Presumed spiny projection neurons (pSPNs) were selected by having a peak width greater than 90 µs, peak to valley duration greater than 560 µs, firing rate lower than 15 Hz and proportion of time in ISIs > 2 s greater than 50%. Light responsive units in D1-Cre and A2a-Cre animals that did not meet these pSPN criteria were excluded from analyses (See Fig S1). Narrow units were classified as units with peak width < 135 µs and peak to valley duration < 230 µs. To select presumed PV neurons (pPV), we first identified neurons with peak width > 100 µs, peak to valley duration between 200 and 700 µs, CV > 0.8 and proportion of time (ISI > 2s) below 15 %. Among these, neurons that increased their firing starting before Center in and sustained this increase until Center Out were classified as pPV. In animals expressing inhibitory opsins the same waveform and firing criteria described above were used to select pPV interneurons.

Among those, cells that were very inhibited within 10 ms and for the duration of the whole light pulse (300 ms) by light pulses were labeled as PV+.

### Analysis of behavior-related activity

We characterized the response of units to events by building peri-event time histograms (PETHs). For all events we analyzed a 1 s window in 25 ms overlapping (12.5 ms shifts) time bins. To determine significant modulation during the hold period we used shuffle tests, comparing the mean firing rate during the 500 ms before the *Go!* Cue (single window) to 10,000 epochs of 500 ms duration sampled randomly from the same session. We used a threshold of p<0.001, one-tailed. To determine if the proportion of modulated units exceeded chance we performed a binomial test (p < 0.01). Group differences were assessed with pairwise Fisher’s exact tests (FDR-corrected, q < 0.05).

To examine activity during movement sequences, for each unit we took spikes occurring in the interval from 70 ms after the *Go!* cue until Side In and convolved them with a Gaussian kernel (σ =7 ms), then averaged across *Contra* or *Ipsi* trials. Units that did not have a minimum of 3Hz average firing during any point in this analysis window were excluded. Distributions of peak time and peak width (measured halfway between 0 and peak firing rate) were compared using the Kolmogorov-Smirnov Test and corrected for multiple comparisons using the Benjamini–Hochberg FDR procedure (q<0.05).

The activity during PV+ inhibition was calculated as above. Laser on or laser off firing rate was normalized to each unit’s maximum trial-averaged firing rate for all *Contra* or *Ipsi* choice trials. The change in firing rate (ΔFR) was calculated by subtracting the normalized average firing rate during laser off trials from the normalized average firing rate during laser on trials. For pSPNs only units with at least 20 spikes within 300 ms after Center Out were considered active.

### Correlations

Spike-time correlations were quantified using cross-correlograms (CCGs) computed in 0.75 ms bins over a ±20 ms window. Only pairs with at least 100 spikes in this CCG window were considered further. For each neuron pair, we created 1000 surrogate data sets by jittering each spike time over a uniform ±5 ms window, preserving firing rates over longer timescales(*32*). At each time lag we labelled the CCG as significant if the observed spike count fell above 99% or below 99% of the surrogate distribution. For pairs recorded in the same tetrode, the CCG was not calculated in the window from -0.75 to 0.75 ms around the reference spikes, since our sorting procedures are not reliable for directly overlapping spikes. CCGs can be prone to artefacts when comparing units recorded from the same electrodes(*94*). To diminish possible problems from imperfect spike sorting, we excluded pairs recorded on the same tetrode that had similar waveforms (waveform correlation > 0.8 for pSPN:pSPN pairs, 0.9 for all others). To assess whether specific subtypes of unit pairs show synchrony or putative inhibition at the population level, each neuron pair was treated as a Bernoulli trial. “Success” meant at least one significant CCG bin in the test window: within ±5 ms for peaks (synchrony) and from 0-4 ms for dips (inhibition). The probability that a pair would show any significant bin by chance (p₀) was estimated as the family-wise false-positive rate, given the per-bin alpha of 0.01 and the total number of bins tested. The number of significant pairs observed across the dataset was then compared to this null expectation using a binomial test.

