## Supplementary material for "Parvalbumin interneurons gate and shape striatal sequences": S1, S2, S3, S4, S5, S6, S7

Supplementary Material
Duhne et al.,2026

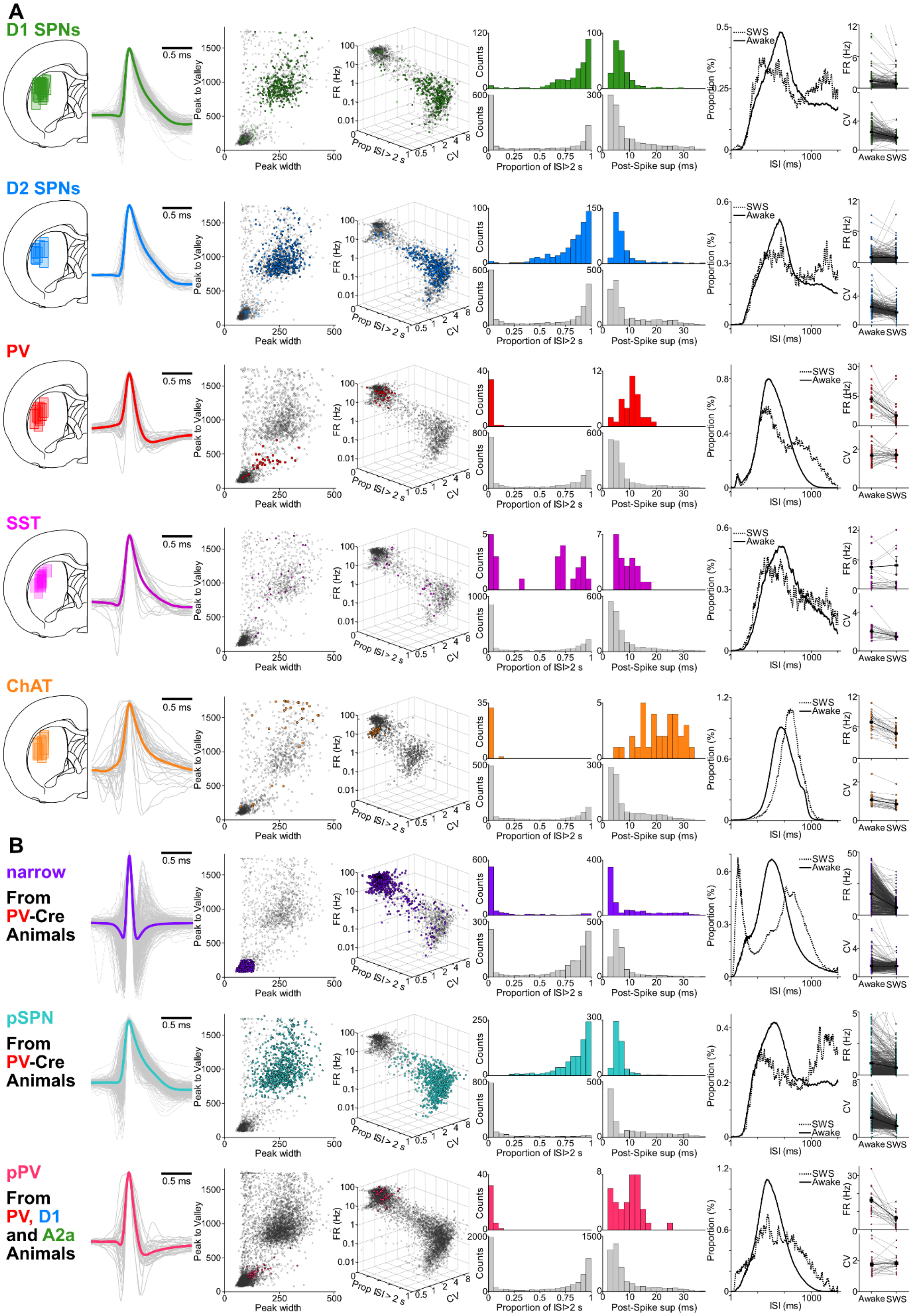

**Fig. S1. Spiking properties of neuron subpopulations.** (**A**) Each row provides information on a distinct opto-tagged neuron type. Columns from left: Schematic of estimated recording areas (rectangles) within DLS, shown on a single atlas hemisphere (Paxinos). For each group: D1 13 rats, 17 fibers, 109 Sessions, 327 D1+, 351 Light responsive units, 1639 total units. D2 8 rats, 8 fibers, 82 Sessions, 389 D2+, 408 Light responsive units, 2247 total units. PV 4 rats, 7 fibers, 51 Sessions, 36 PV+, 1880 total units. ChAT 8 rats, 8 fibers, 32 Sessions, 33 ChAT+, 929 total units. SST 8 rats, 10 fibers, 41 Sessions, 23 SST+, 1676 total units. Mean normalized spike waveforms for each tagged unit (grey) with color line showing grand average; scatter plots of waveform duration (same format as Fig. 1C; tagged cells marked in color). Colored unfilled circles indicate non SPN light responsive units in D1 and D2 recordings. Scatter plot of firing pattern measures, firing rate in Hz, Proportion of time in inter-spike-intervals(ISI) longer than 2 s (Prop ISI>2s) and ISI variation coeficient; Histograms of Prop (ISI>2s) and post-spike suppression for tagged (top) and untagged (bottom) cells, recorded during bandit task performance; ISI distribution in Awake and slow wave sleep (SWS) epochs; Top, mean FR and bottom mean CV for individual units of each population in awake and SWS epochs.

(**B**) Properties of unidentified striatal subpopulations (same format as A). Top, "narrow" units (from the same PV-Cre recordings as A) defined as those with waveform peak width less than 135 μs and peak-valley duration less than 230μs. Next row, untagged cells recorded in PV-Cre rats and presumed to be SPNs (pSPNs) based on waveform duration, high variation coefficient, and tonicity (see methods). Bottom, untagged cells recorded in D1-Cre, A2a-Cre and PV-Cre rats and now presumed to be PV+ ("pPV") based on waveform duration and elevated firing during the hold period.

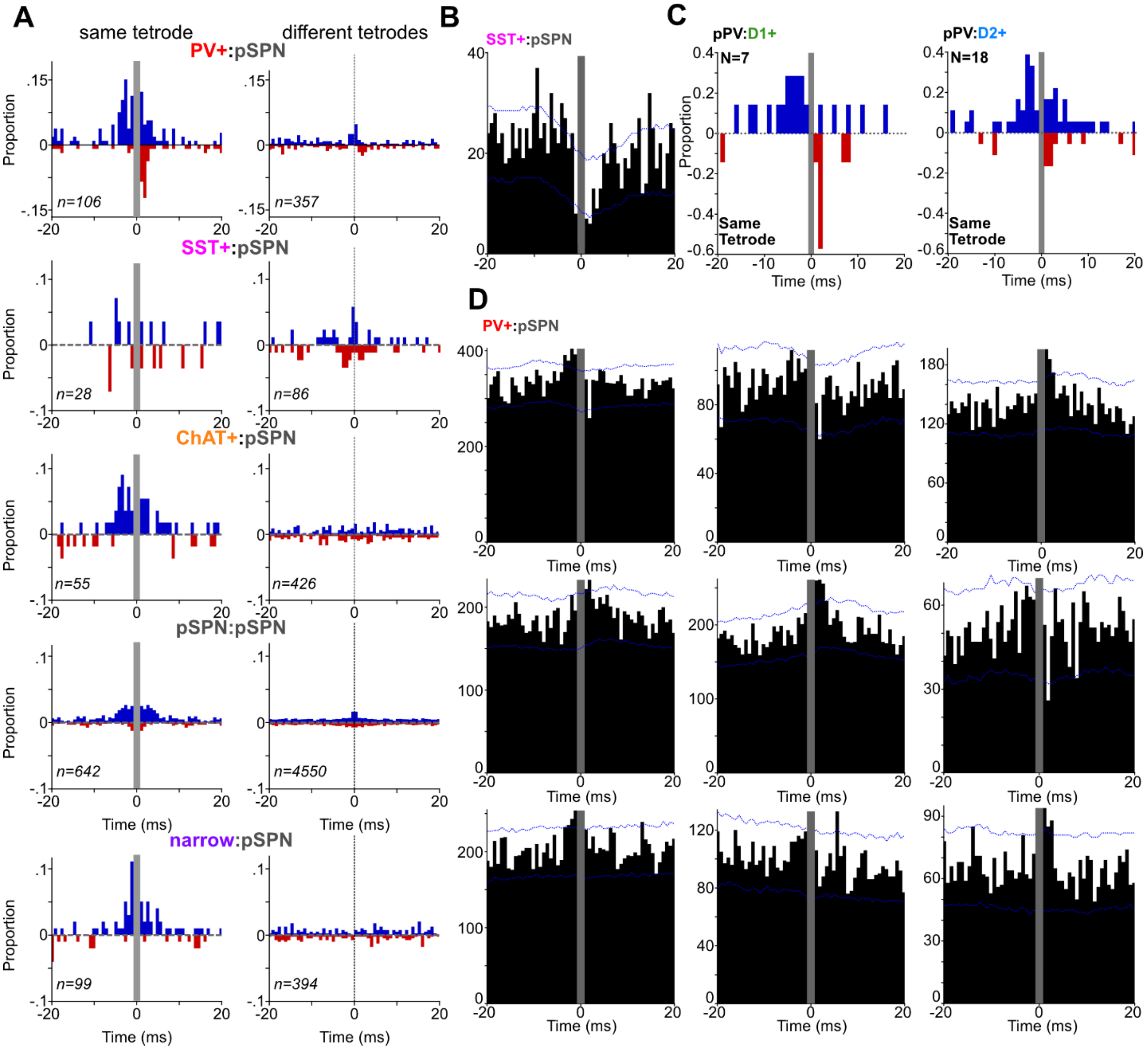


**Fig. S2.** (**A**)Proportion of pairs with significant positive (blue) and negative (red) CC as in Figure 1I but for pairs recorded in the same (left) and different (right) tetrodes. (**B**) Rare example of an SST:pSPN pair recorded from the same tetrode showing possible fast inhibition. Same format at Fig. 4Bi. (**C**) Proportions of pPV:D1+ pairs (top) and pPV:D2+ pairs (bottom) with significant cross-correlograms at each time bin. Same format as Fig. 4C. (**D**) Example of cross-correlograms between a single PV+ and a set of pSPNs recorded simultaneously from the same tetrode. Same format as Fig. 4B. Presence and timing of peaks and dips varies between pairs even from the same PV+.

Supplementary Material
Duhne et al.,2026

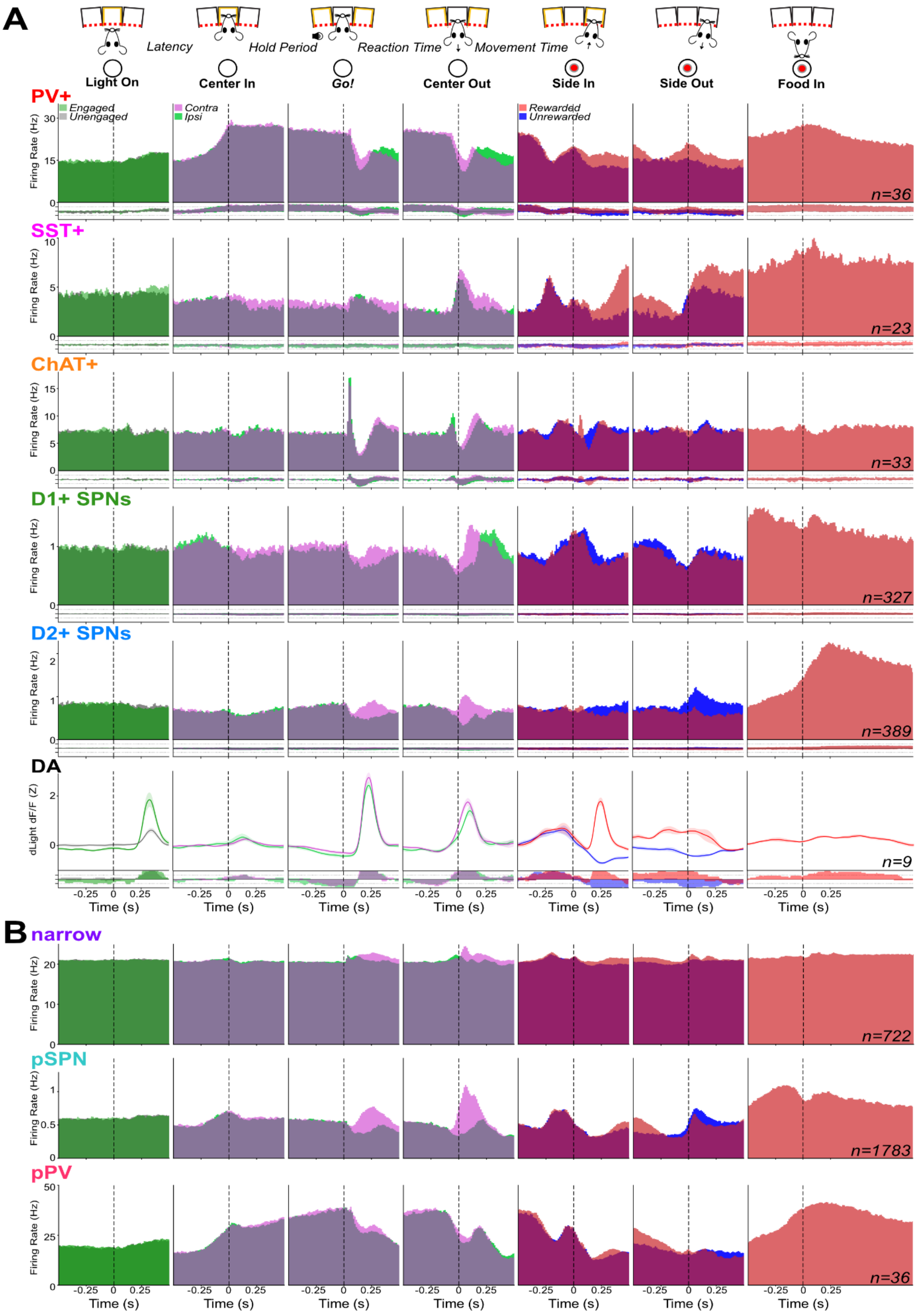

**Fig. S3. Extended view of event-aligned spiking.** (**A**) Same format as Fig. 2A, but showing all task events. Bottom proportion of significantly modulated units. (**B**). Mean event-aligned activity of untagged "narrow", pPV and pSPN units, as in Fig. 2A.

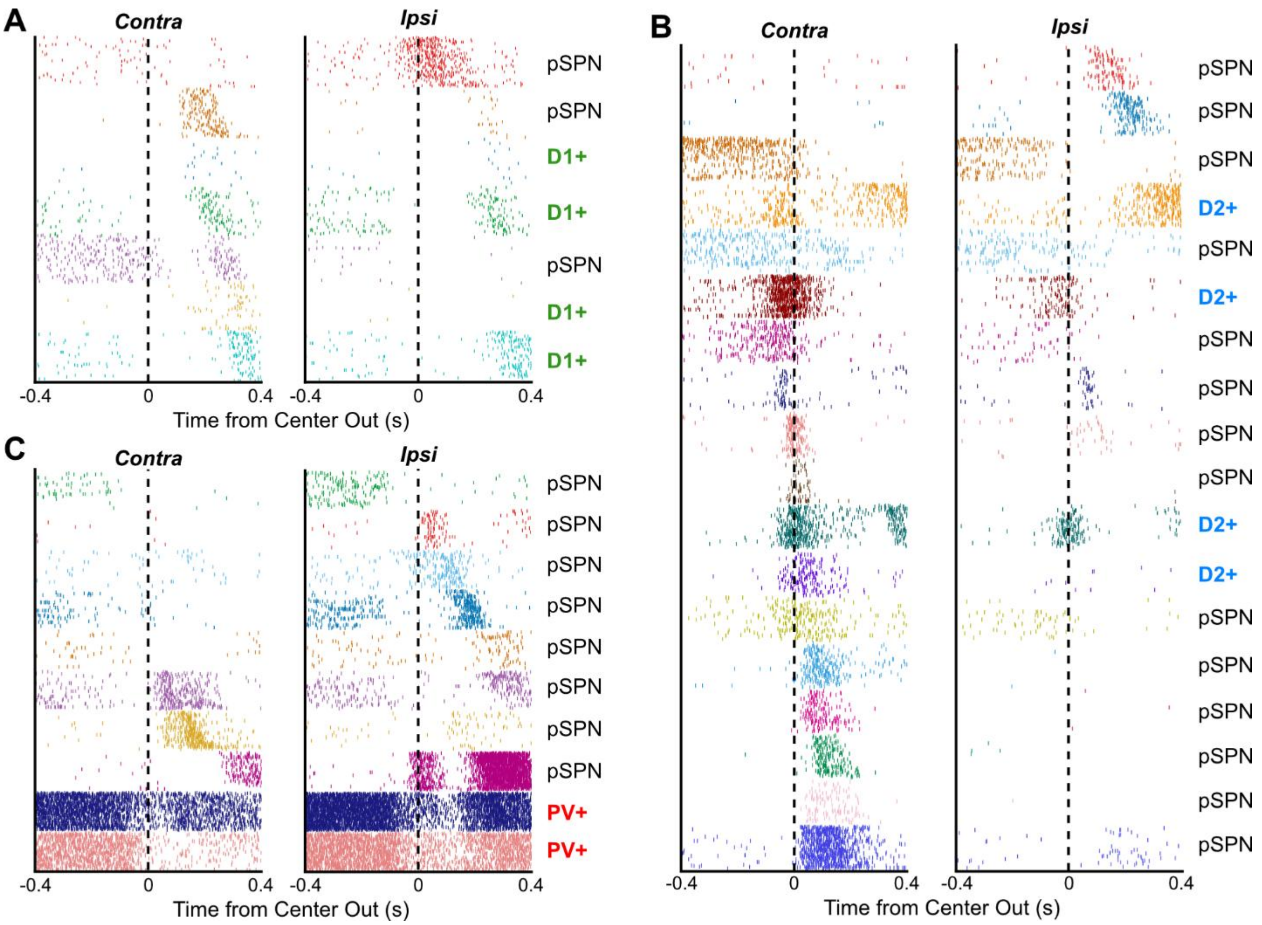


**Fig. S4. Example sessions comparing *Contra* and *Ipsi* movement sequences.** Format as in Fig. 4A-C showing both movement directions. For each unit, trials are ordered by movement time duration. (**A**) Additional example D1+ session, (**B**) D2+ session; for these, neurons were organized according to their peak time in *Contra* trials. (**C**) Example PV+ recording session, neurons were sorted according to peak time during *Ipsi* trials.

Supplementary Material
Duhne et al.,2026

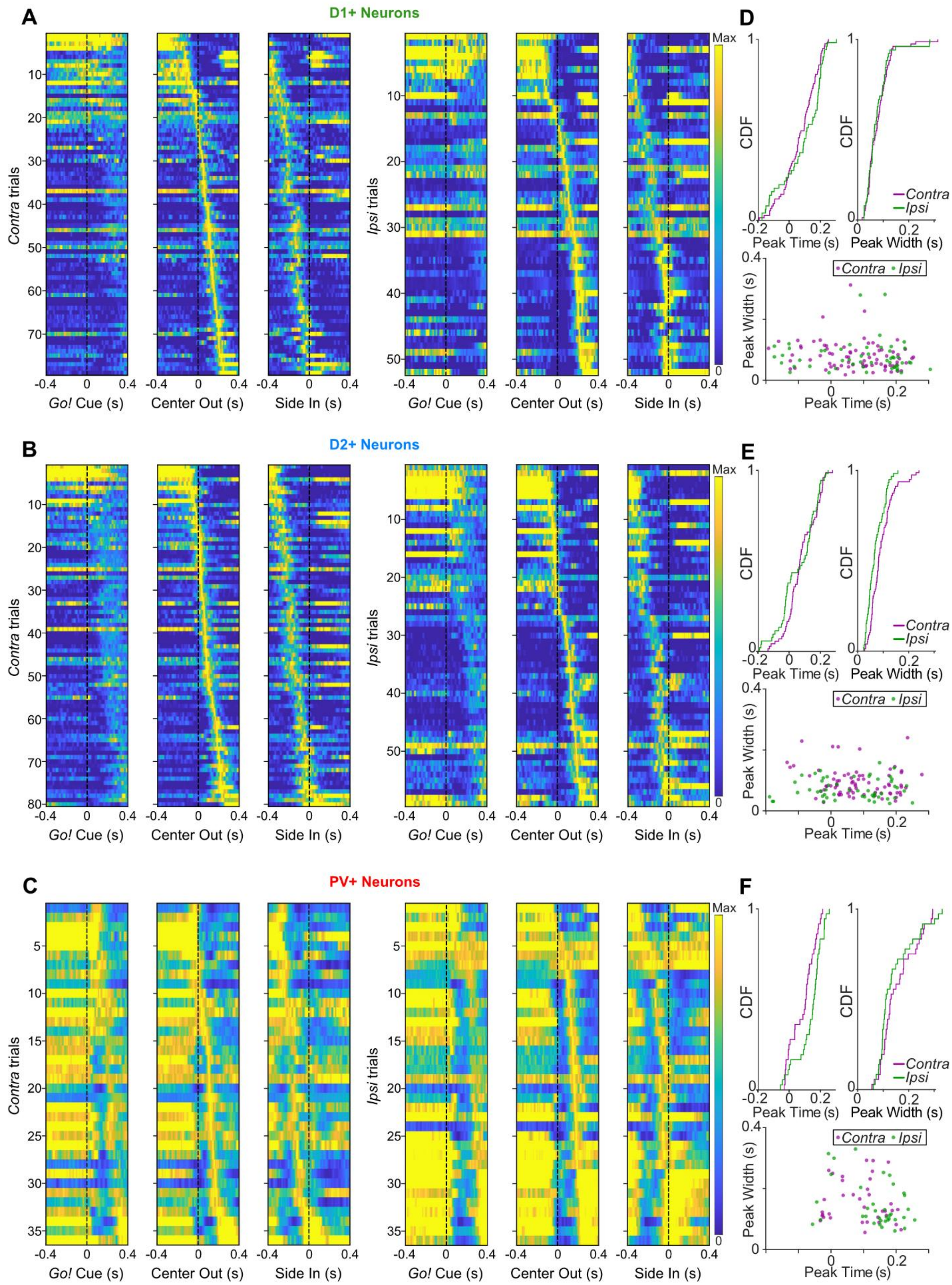

**Fig. S5. Sequences are movement-locked, not cue-locked.** Firing sequences for D1+ (**A**), D2+ (**B**), and PV+ (**C**), neurons as in Fig 4A-C showing alignment to Go!, Center out and Side In events for each side. The activity is sharper when aligned to Center out. Neurons were sorted separately for each side. Note that less SPNs participate in *Ipsi* choice trials D1+ 15.9% and D2+ 15.2% compared to *Contra* choice trials. All PVs (36/36) participated in both *Ipsi* and *Contra* trials. (**D-F**) Distributions and Scatter of firing peak times and peak widths in *Contra* and *Ipsi* trials. There was no significant difference between the peak width and peak time distributions within neuronal type comparing *Contra* and *Ipsi* trials.

Supplementary Material
Duhne et al.,2026

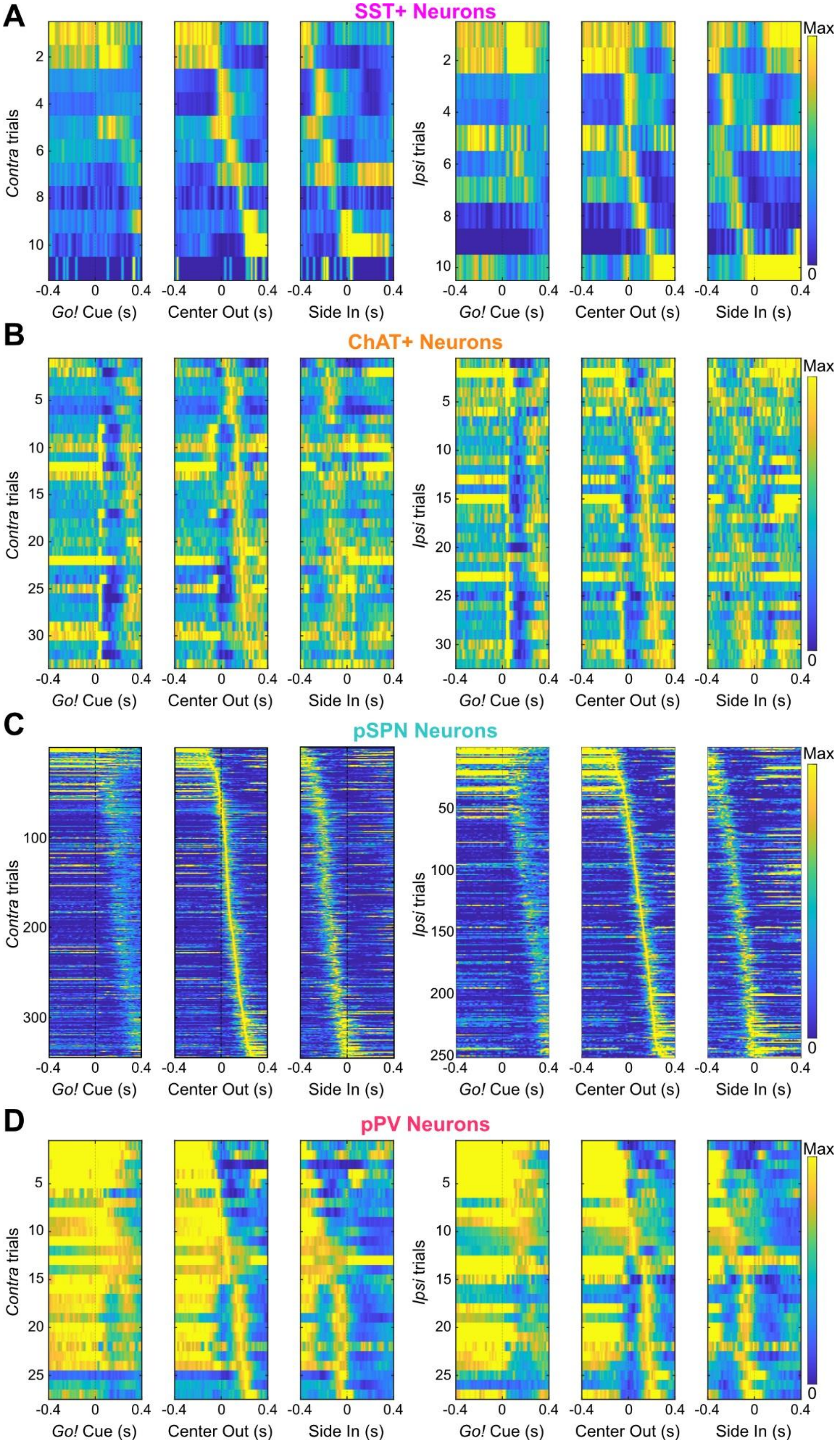

**Fig. S6. Sequential activity in other cell populations.** Firing sequences as in Figs S5A-C and 4A-C for other cell types. (**A**), SST+ neurons: only a subset of SST + neurons (11/23 47.8%) met the 3 Hz FR criterion to be considered part of the movement sequence. (**B**) ChAT+ neurons: all recorded ChAT+ (33/33) neurons participated in the movement sequence. (**C**) pSPNs recorded from D1-Cre, A2a-cre and PV-Cre rats: 335/2163 neurons (15.5%) met the 3 Hz FR firing criterion for inclusion. (**D**) pPV neurons: a large proportion of these units (27/36 75%) participated in the sequence.

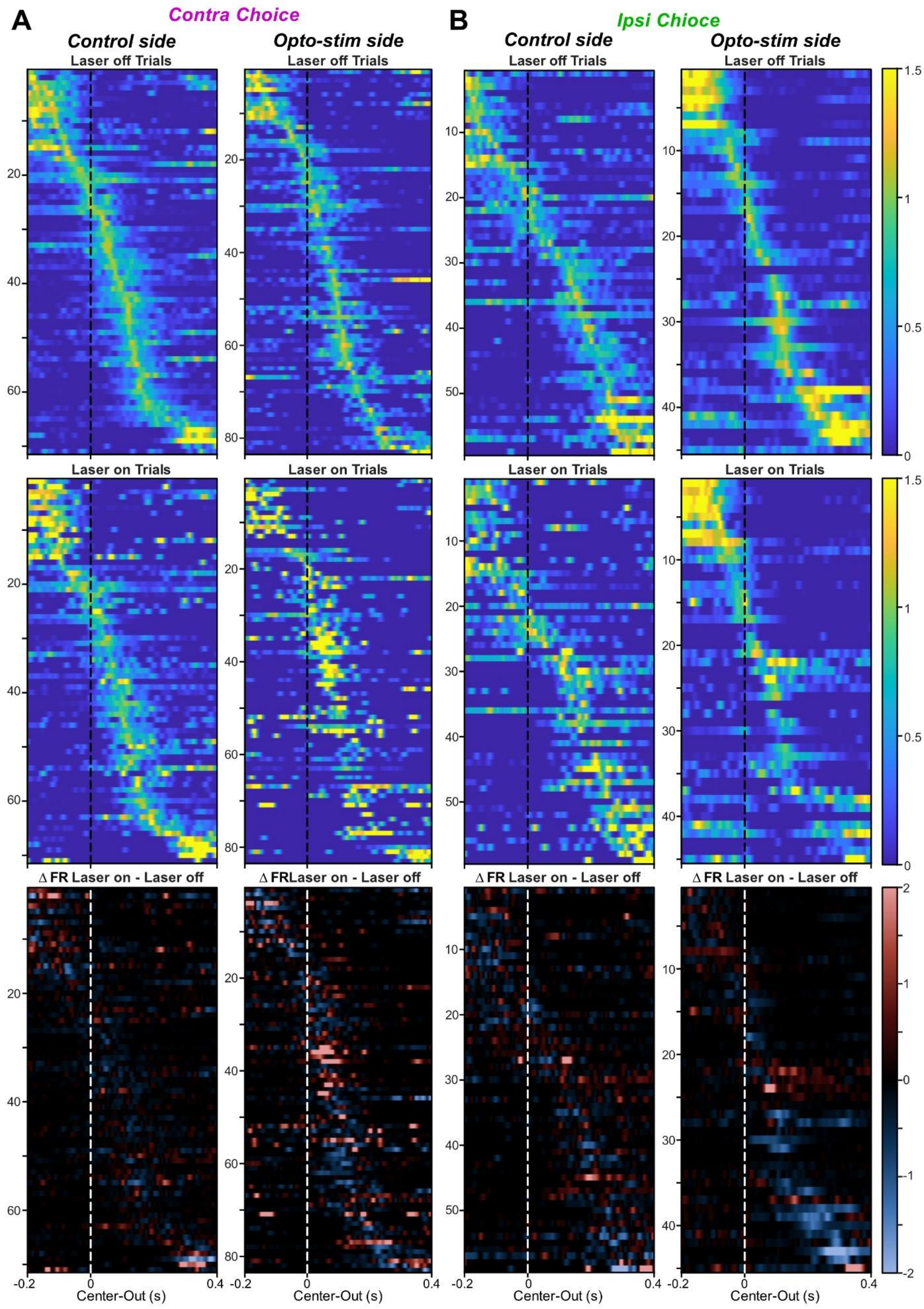
A
Contra Choice
Control side
Opto-stim side
Laser off Trials
Laser on Trials
Δ FR Laser on - Laser off
Δ FRLaser on - Laser off
B
Ipsi Chioce
Control side
Opto-stim side
Center-Out (s)
-0.2
0
0.4
1.5
1
0.5
0
2
-1
-2

**Fig. S7. Sequential activity in other cell populations.** Firing sequences as in Figs S5A-C, S6A-D and 4A-C for trials with no laser stimulation (Laser off, top), trials with laser on (middle). Change in firing rate during sequence as in Fig. 5 B,C (bottom). The units were aligned according to firing timing when averaging all trials. (**A**) Sequence all active pSPNs in trials where *Contra* to laser stimulation and unit was made. (**B**) Same as A but for trials of *Ipsi* choice.
