## Appendix for "Parvalbumin interneurons gate and shape striatal sequences"

**Appendix. Activity patterns of individual PV+ neurons.**

Top row from left to right:

- Cell ID and firing properties (as shown in Fig. 1H).
  - $FR_{RL}$: Firing Rate During Bandit task.
  - $CV_{Aw}$: Firing variation coefficient during wake period.
  - $FR_{SWS}$: Firing Rate During slow wave sleep.
  - $CV_{SWS}$: Firing variation coefficient during slow wave sleep.
  - $Proportion_{ISI>2s}$: Proportion of interspike intervals greater than 2 seconds.
  - Post Spike Supp: Time in autocorrelogram to reach half of the mean amplitude.
  - P2V: time within average spike waveform from peak to valley.
  - PeakWidth: average spike waveform peak width at half-maximum.
- Average waveform, during light delivery (red) and outside of light delivery (black) (As in Fig. 1E).
- Raster plot showing the response to 10 ms light pulse delivery at 10 mW (As in Fig. 1D).
- Latency to spike after light delivery (red); Latency to spike from random times during laser session (black)
  - Reliability: portion of laser pulses in which a spike occurs.
  - P value for Wilcoxon rank test.
- Autocorrelation of cell firing (as in Fig. 1I).
- Inter spike interval (ISI) distribution during waking (black) and slow wave sleep (SWS, gray).

Second row from left to right:

- Latency (Light On to Center-Nose-In, CNI) distribution for the behavioral session in which the cell was recorded.
- Reaction Time distribution.
- Movement Time distribution.
- Bandit task behavior summary for the same session.
  - Top: Individual trial choices. Magenta, contraversive trials; Green, ipsiversive trials. Tall ticks, rewarded trials; Short ticks, unrewarded trials.
  - Middle: Dashed lines show nominal reward probability, for contraversive (magenta) and ipsiversive (green) choices. Dotted lines show smoothed (10-trial moving boxcar moving average) rat choice probability for contraversive (magenta) and ipsiversive (green) sides.
  - Bottom: trial value and cell firing rate (1 minute bins; as in Fig. 4A). Dot colors indicate trial value in terciles (high: red; medium: brown; low: black).

Third Row:

Raster plot and peri-event time histograms during the Bandit task.

- For Light-On, trials are separated by engaged (latency < 1 s, green) and unengaged (latency > 1 s, gray; as in Fig. 2).
- For Center-In, Go Cue and Center-Out, trials are separated by choice (contraversive, magenta; ipsiversive, green, as in Fig. 3).
- Side-In and Side-Out events are separated by outcome (rewarded, red; unrewarded, blue; as in Figs. 2, 4).
- PETH for food port entry on unrewarded trials is not shown as animals did not consistently approach the food port on those trials.

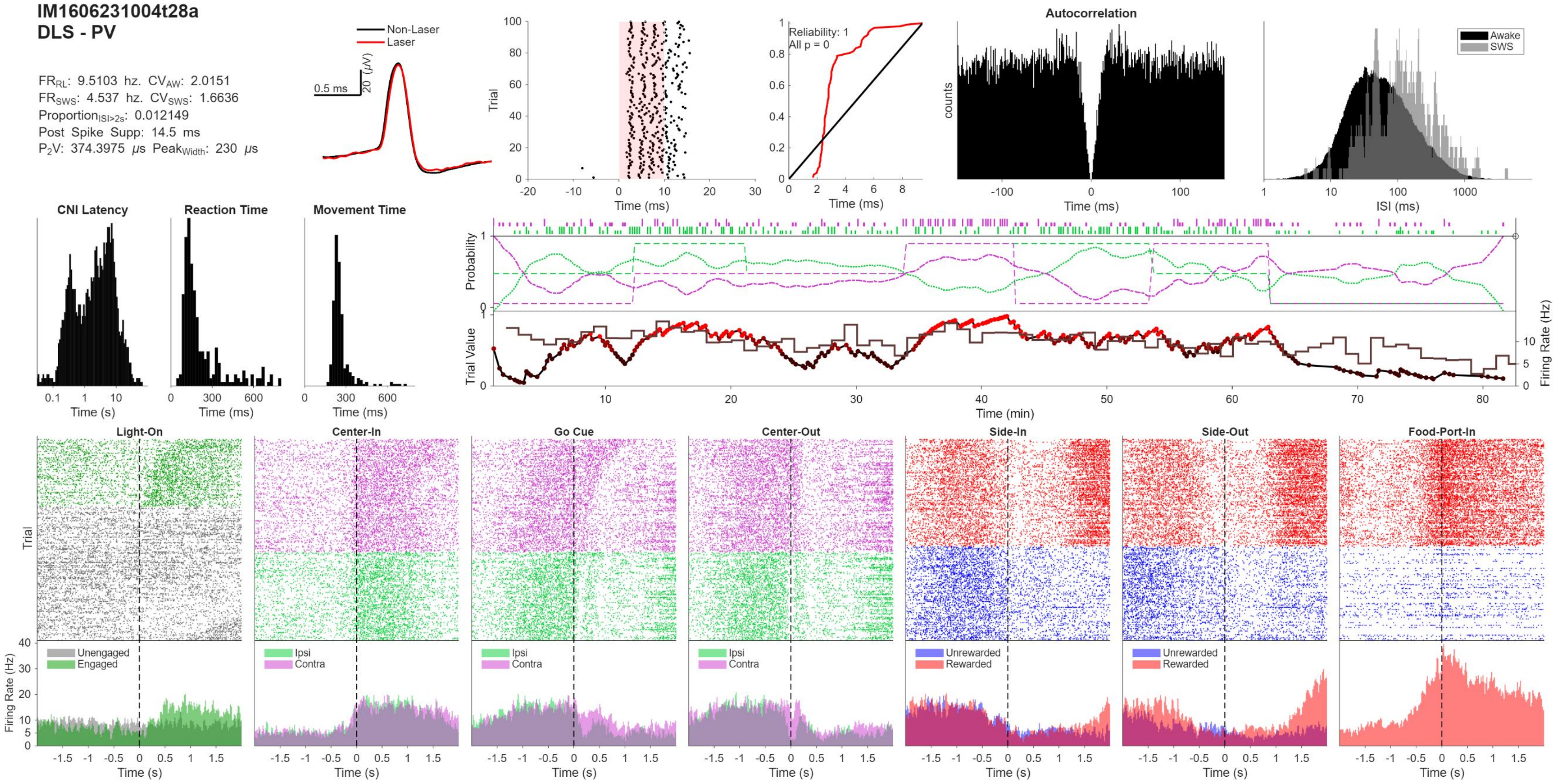
IM1606231004t28a
DLS - PV
FR_RL: 9.5103 hz. CV_AW: 2.0151
FR_SWS: 4.537 hz. CV_SWS: 1.6636
Proportion_ISI>2s: 0.012149
Post Spike Supp: 14.5 ms
P2V: 374.3975 μs Peak_Width: 230 μs
Non-Laser
Laser
0.5 ms
20 (μV)
Trial
Time (ms)
Reliability: 1
All p = 0
Time (ms)
Autocorrelation
counts
Time (ms)
Awake
SWS
ISI (ms)
CNI Latency
Time (s)
Reaction Time
Time (ms)
Movement Time
Time (ms)
Probability
Trial Value
Firing Rate (Hz)
Time (min)
Light-On
Center-In
Go Cue
Center-Out
Side-In
Side-Out
Food-Port-In
Trial
Firing Rate (Hz)
Unengaged
Engaged
Ipsi
Contra
Unrewarded
Rewarded
Time (s)

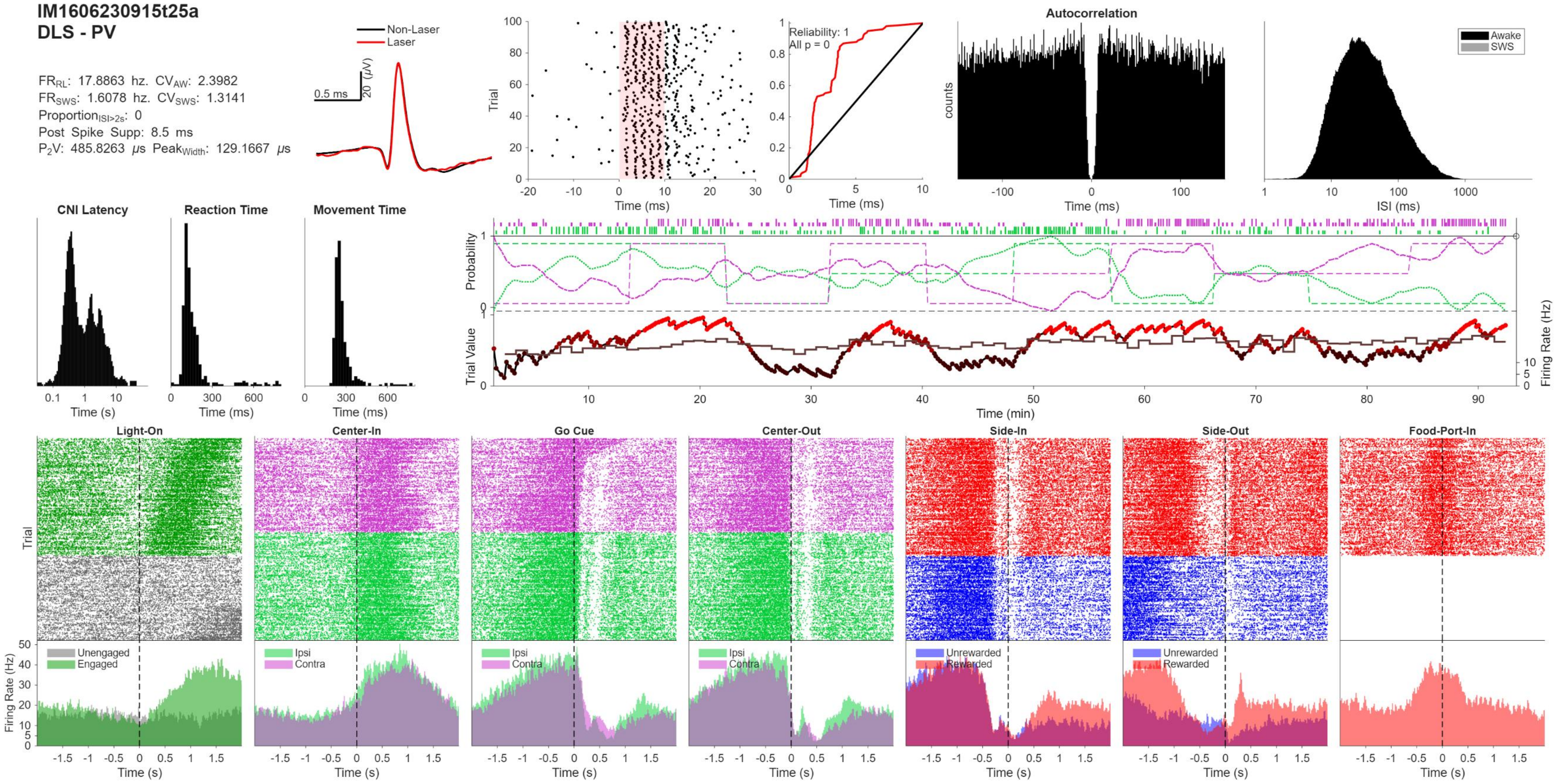
IM1606230915t25a
DLS - PV
FRRL: 17.8863 hz. CVAW: 2.3982
FRSWS: 1.6078 hz. CVSWS: 1.3141
ProportionISI>2s: 0
Post Spike Supp: 8.5 ms
P2V: 485.8263 μs PeakWidth: 129.1667 μs
Non-Laser
Laser
0.5 ms
20 (μV)
Trial
Time (ms)
Reliability: 1
All p = 0
Autocorrelation
counts
Awake
SWS
ISI (ms)
CNI Latency
Reaction Time
Movement Time
Time (s)
Probability
Trial Value
Firing Rate (Hz)
Time (min)
Light-On
Center-In
Go Cue
Center-Out
Side-In
Side-Out
Food-Port-In
Unengaged
Engaged
Ipsi
Contra
Unrewarded
Rewarded

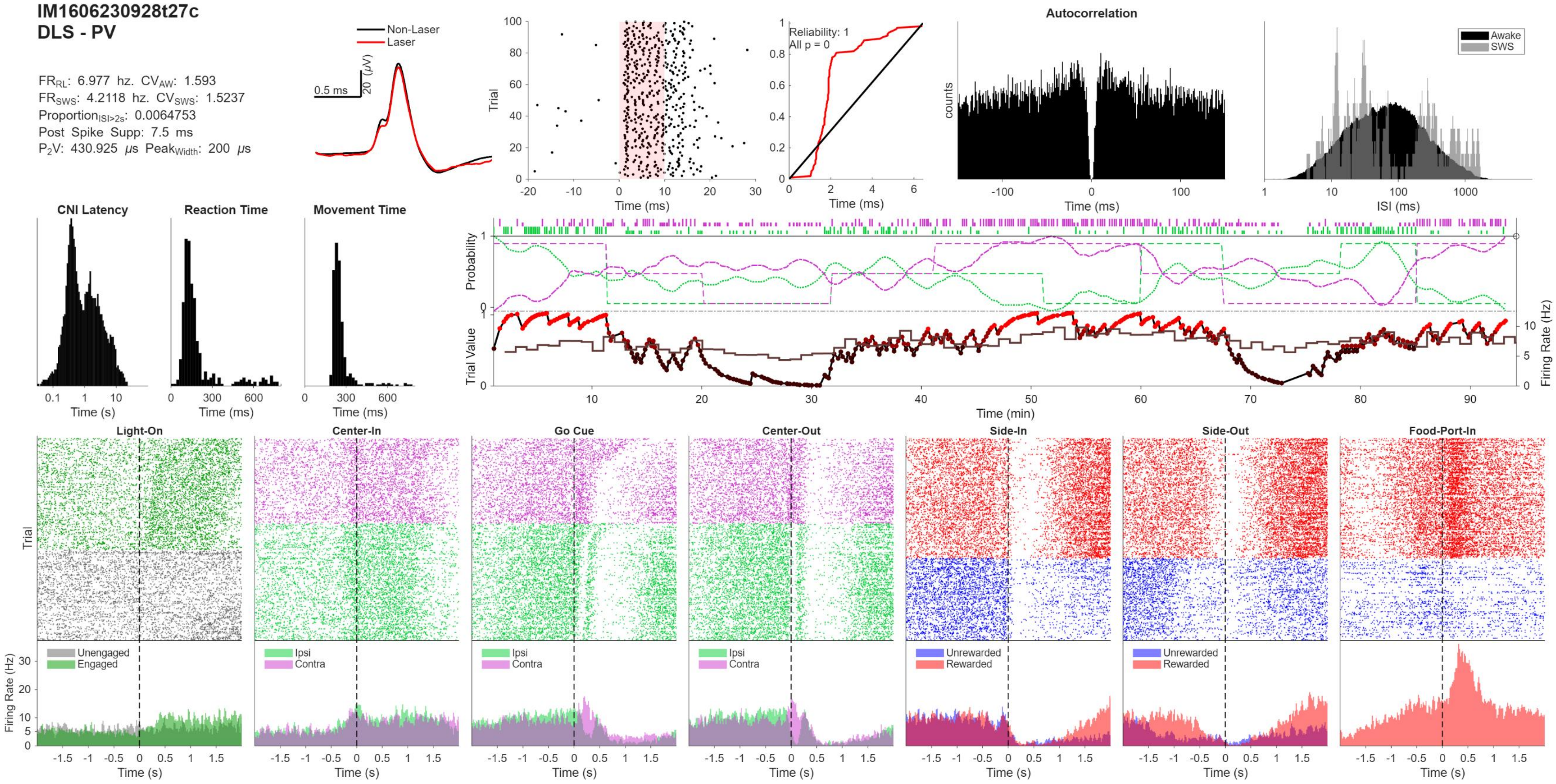
IM1606230928t27c
DLS - PV
FR_RL: 6.977 hz. CV_AW: 1.593
FR_SWS: 4.2118 hz. CV_SWS: 1.5237
Proportion_ISI>2s: 0.0064753
Post Spike Supp: 7.5 ms
P2V: 430.925 μs Peak_Width: 200 μs
Non-Laser
Laser
0.5 ms
20 (μV)
Trial
Time (ms)
Reliability: 1
All p = 0
Autocorrelation
counts
Awake
SWS
ISI (ms)
CNI Latency
Reaction Time
Movement Time
Time (s)
Probability
Trial Value
Firing Rate (Hz)
Time (min)
Light-On
Center-In
Go Cue
Center-Out
Side-In
Side-Out
Food-Port-In
Unengaged
Engaged
Ipsi
Contra
Unrewarded
Rewarded

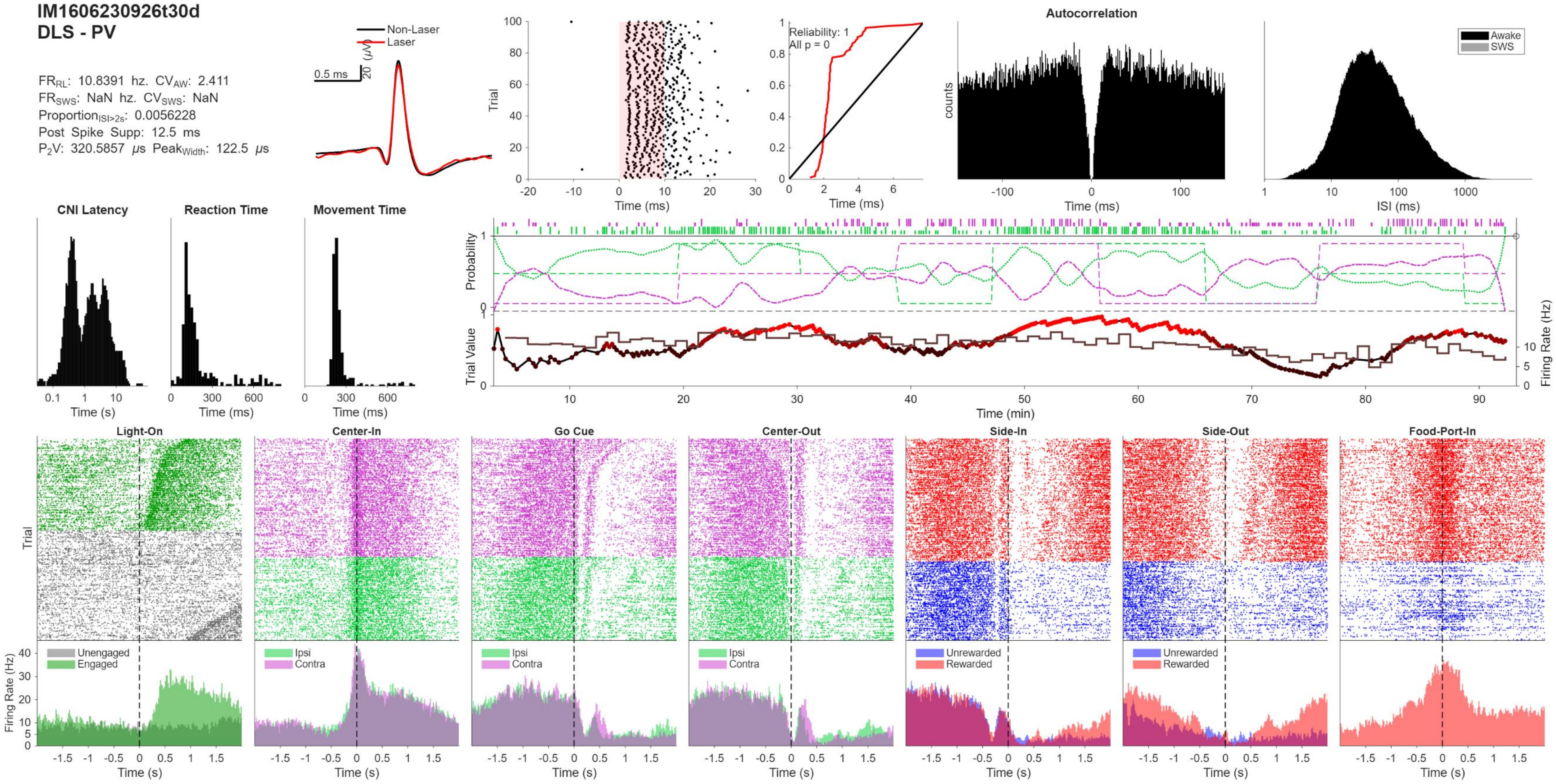
IM1606230926t30d
DLS - PV
FR_RL: 10.8391 hz. CV_AW: 2.411
FR_SWS: NaN hz. CV_SWS: NaN
Proportion_ISI>2s: 0.0056228
Post Spike Supp: 12.5 ms
P2V: 320.5857 μs Peak_Width: 122.5 μs
Non-Laser
Laser
0.5 ms
20 (μV)
Trial
Time (ms)
Reliability: 1
All p = 0
Autocorrelation
counts
Awake
SWS
ISI (ms)
CNI Latency
Reaction Time
Movement Time
Time (s)
Probability
Trial Value
Firing Rate (Hz)
Time (min)
Light-On
Center-In
Go Cue
Center-Out
Side-In
Side-Out
Food-Port-In
Unengaged
Engaged
Ipsi
Contra
Unrewarded
Rewarded
Firing Rate (Hz)

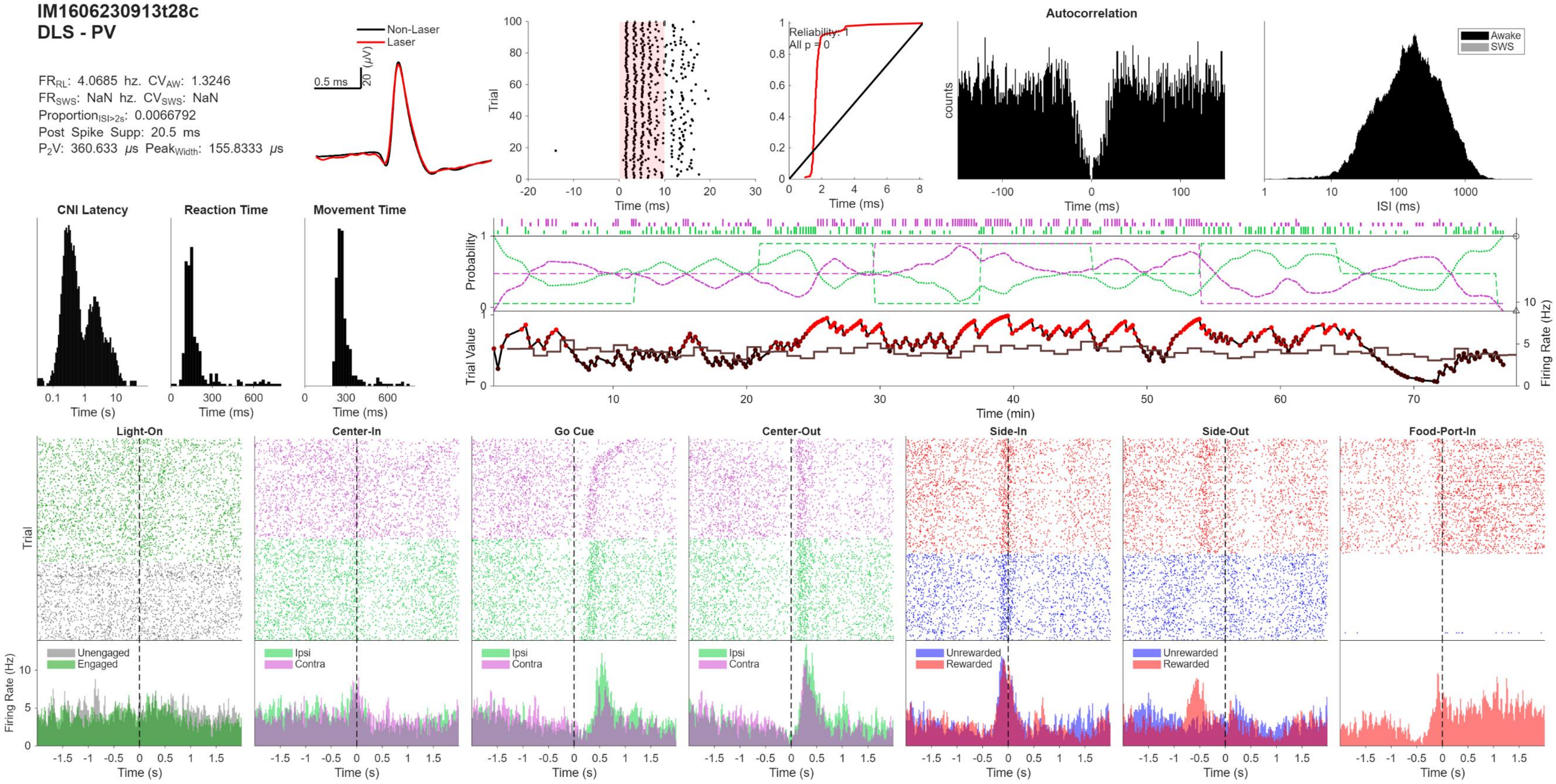
IM1606230913t28c
DLS - PV
FR$_{RL}$: 4.0685 hz. CV$_{AW}$: 1.3246
FR$_{SWS}$: NaN hz. CV$_{SWS}$: NaN
Proportion$_{ISI>2s}$: 0.0066792
Post Spike Supp: 20.5 ms
P$_2$V: 360.633 $\mu$s Peak$_{Width}$: 155.8333 $\mu$s
Non-Laser
Laser
0.5 ms
20 ($\mu$V)
Trial
Time (ms)
Reliability: 1
All p = 0
Autocorrelation
counts
Awake
SWS
ISI (ms)
CNI Latency
Reaction Time
Movement Time
Time (s)
Probability
Trial Value
Firing Rate (Hz)
Time (min)
Light-On
Center-In
Go Cue
Center-Out
Side-In
Side-Out
Food-Port-In
Unengaged
Engaged
Ipsi
Contra
Unrewarded
Rewarded

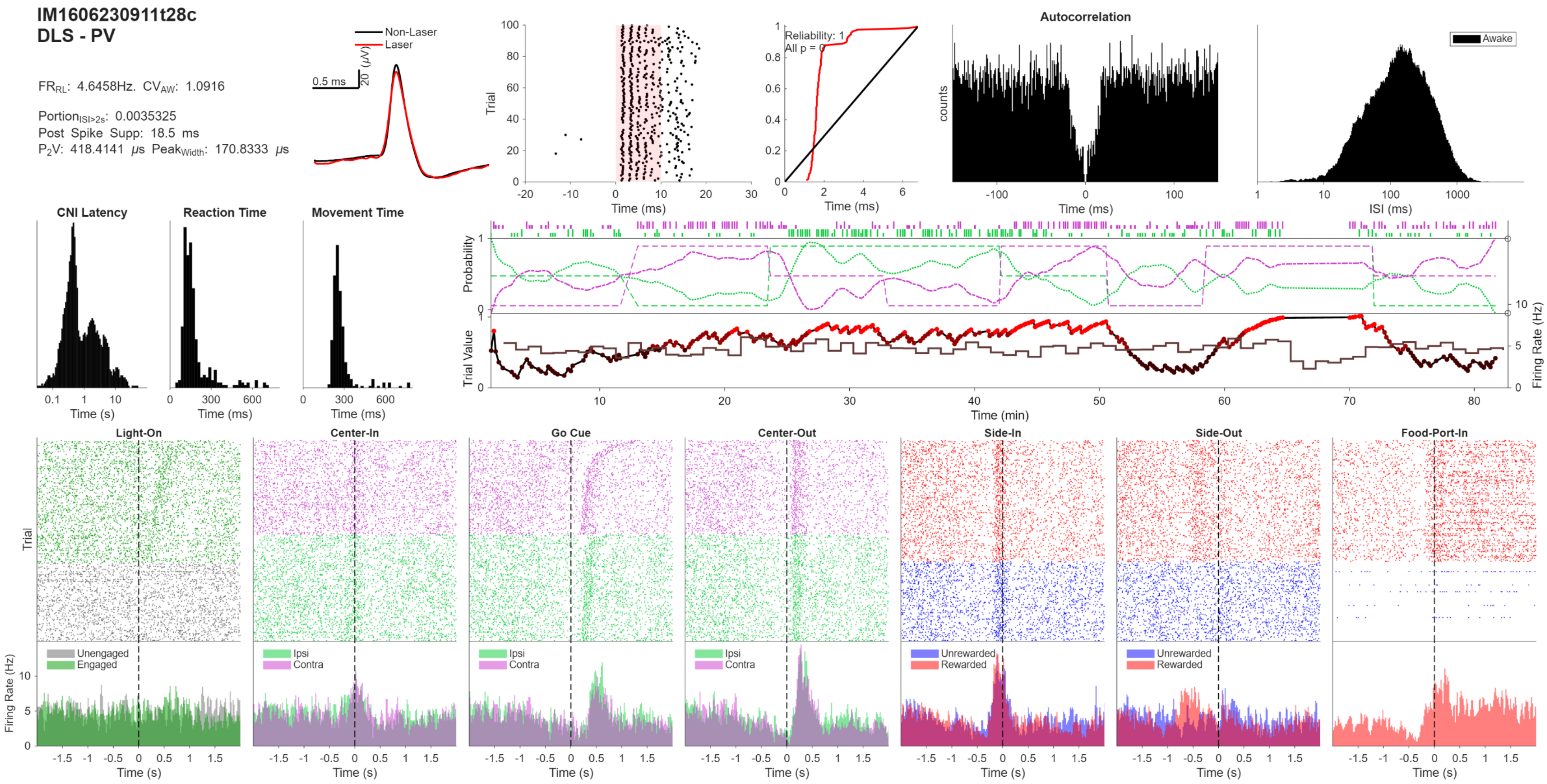

IM1606230911t28c
DLS - PV
FR_RL: 4.6458Hz. CV_AW: 1.0916
Portion_ISI>2s: 0.0035325
Post Spike Supp: 18.5 ms
P2V: 418.4141 μs Peak_Width: 170.8333 μs
Non-Laser
Laser
0.5 ms
20 (μV)
Trial
Time (ms)
Reliability: 1
All p = 0
Autocorrelation
counts
Awake
ISI (ms)
CNI Latency
Reaction Time
Movement Time
Time (s)
Probability
Trial Value
Firing Rate (Hz)
Time (min)
Light-On
Center-In
Go Cue
Center-Out
Side-In
Side-Out
Food-Port-In
Unengaged
Engaged
Ipsi
Contra
Unrewarded
Rewarded

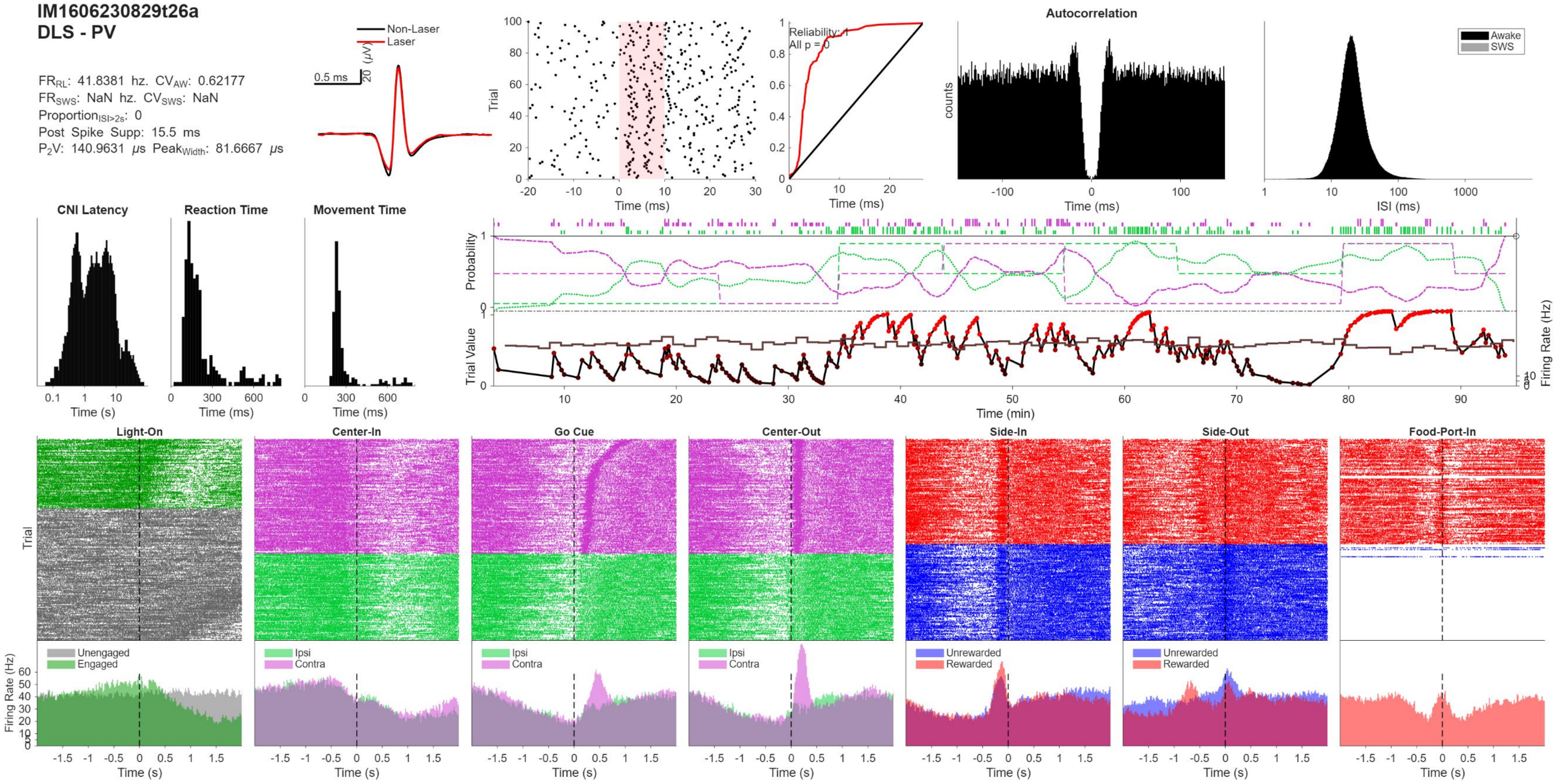
IM1606230829t26a
DLS - PV
FR_RL: 41.8381 hz. CV_AW: 0.62177
FR_SWS: NaN hz. CV_SWS: NaN
Proportion_ISI>2s: 0
Post Spike Supp: 15.5 ms
P2V: 140.9631 μs Peak_Width: 81.6667 μs
Non-Laser
Laser
0.5 ms
20 (μV)
Trial
Time (ms)
Reliability: 1
All p = 0
Autocorrelation
counts
Awake
SWS
ISI (ms)
CNI Latency
Reaction Time
Movement Time
Time (s)
Time (ms)
Probability
Trial Value
Firing Rate (Hz)
Time (min)
Light-On
Center-In
Go Cue
Center-Out
Side-In
Side-Out
Food-Port-In
Trial
Unengaged
Engaged
Ipsi
Contra
Unrewarded
Rewarded
Firing Rate (Hz)
Time (s)

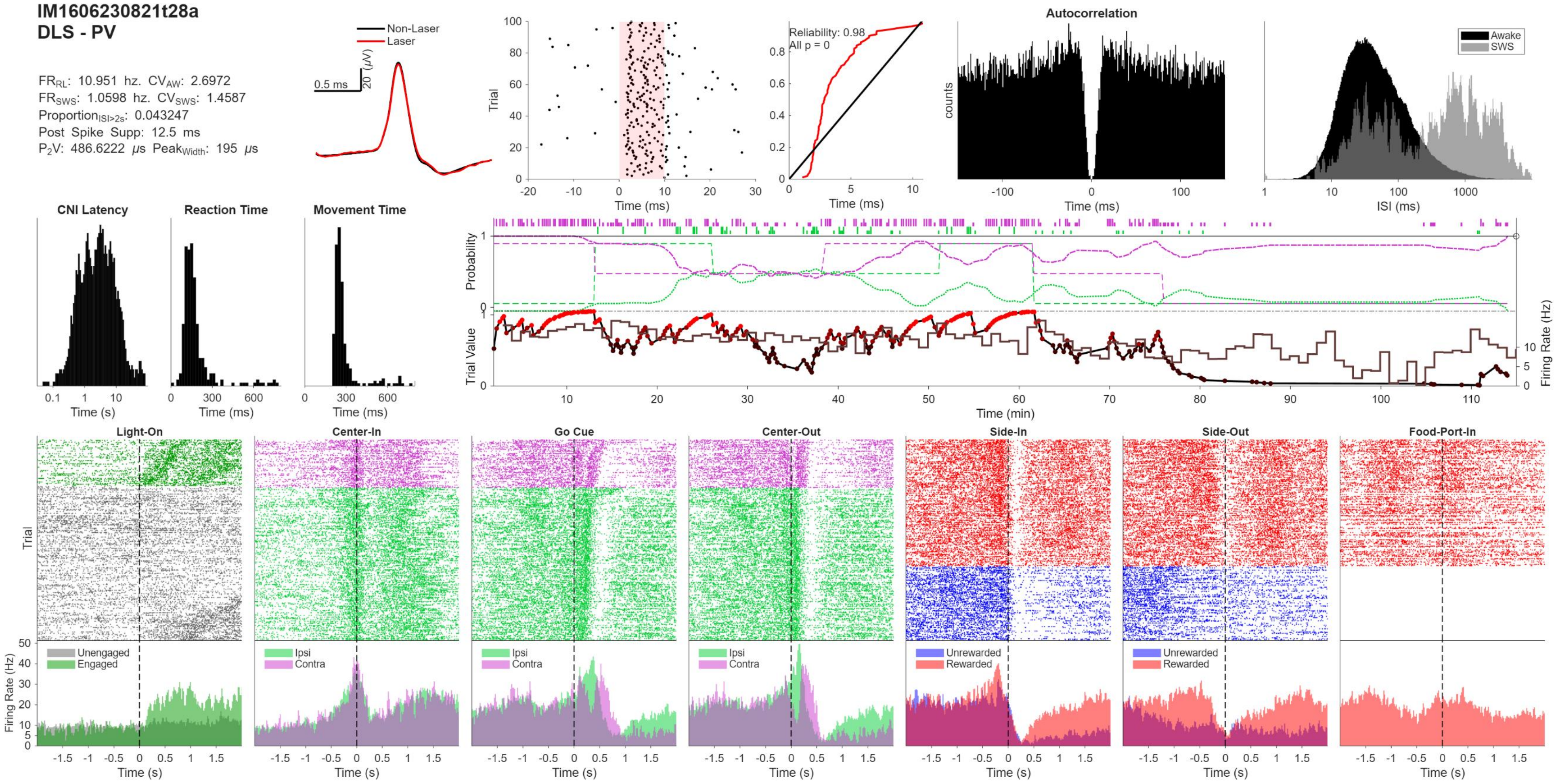
IM1606230821t28a
DLS - PV
$FR_{RL}$: 10.951 hz. $CV_{AW}$: 2.6972
$FR_{SWS}$: 1.0598 hz. $CV_{SWS}$: 1.4587
$Proportion_{ISI>2s}$: 0.043247
Post Spike Supp: 12.5 ms
$P_2V$: 486.6222 μs $Peak_{Width}$: 195 μs
Non-Laser
Laser
0.5 ms
20 (μV)
Trial
Time (ms)
Reliability: 0.98
All p = 0
Time (ms)
Autocorrelation
counts
Time (ms)
Awake
SWS
ISI (ms)
CNI Latency
Time (s)
Reaction Time
Time (ms)
Movement Time
Time (ms)
Probability
Trial Value
Firing Rate (Hz)
Time (min)
Light-On
Center-In
Go Cue
Center-Out
Side-In
Side-Out
Food-Port-In
Trial
Firing Rate (Hz)
Unengaged
Engaged
Ipsi
Contra
Unrewarded
Rewarded
Time (s)

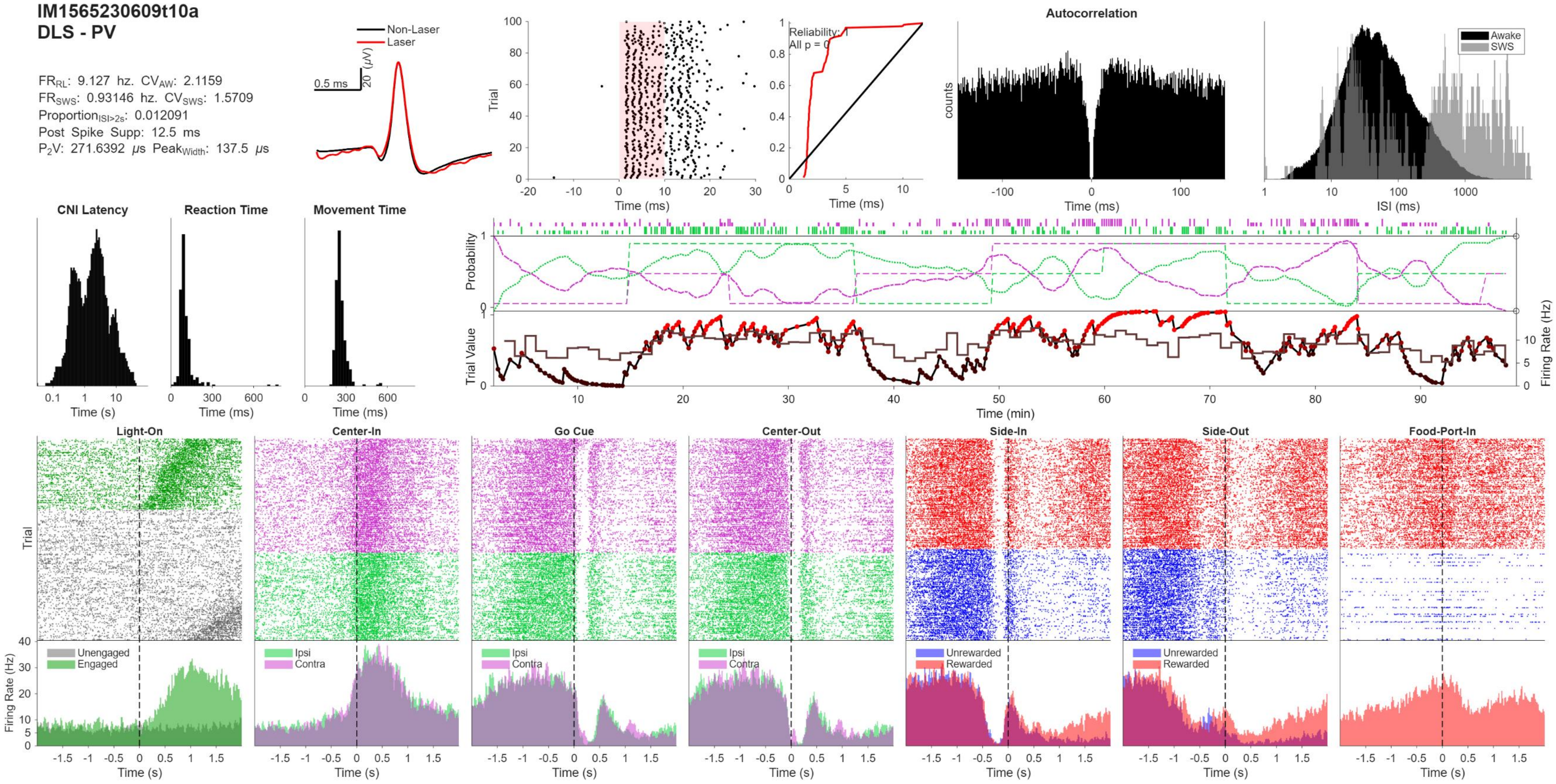
IM1565230609t10a
DLS - PV
FR$_{RL}$: 9.127 hz. CV$_{AW}$: 2.1159
FR$_{SWS}$: 0.93146 hz. CV$_{SWS}$: 1.5709
Proportion$_{ISI>2s}$: 0.012091
Post Spike Supp: 12.5 ms
P$_2$V: 271.6392 μs Peak$_{Width}$: 137.5 μs
Non-Laser
Laser
0.5 ms
20 (μV)
Trial
Time (ms)
Reliability: 1
All p = 0
Autocorrelation
counts
Awake
SWS
ISI (ms)
CNI Latency
Reaction Time
Movement Time
Time (s)
Time (ms)
Probability
Trial Value
Firing Rate (Hz)
Time (min)
Light-On
Center-In
Go Cue
Center-Out
Side-In
Side-Out
Food-Port-In
Unengaged
Engaged
Ipsi
Contra
Unrewarded
Rewarded
Firing Rate (Hz)
Time (s)

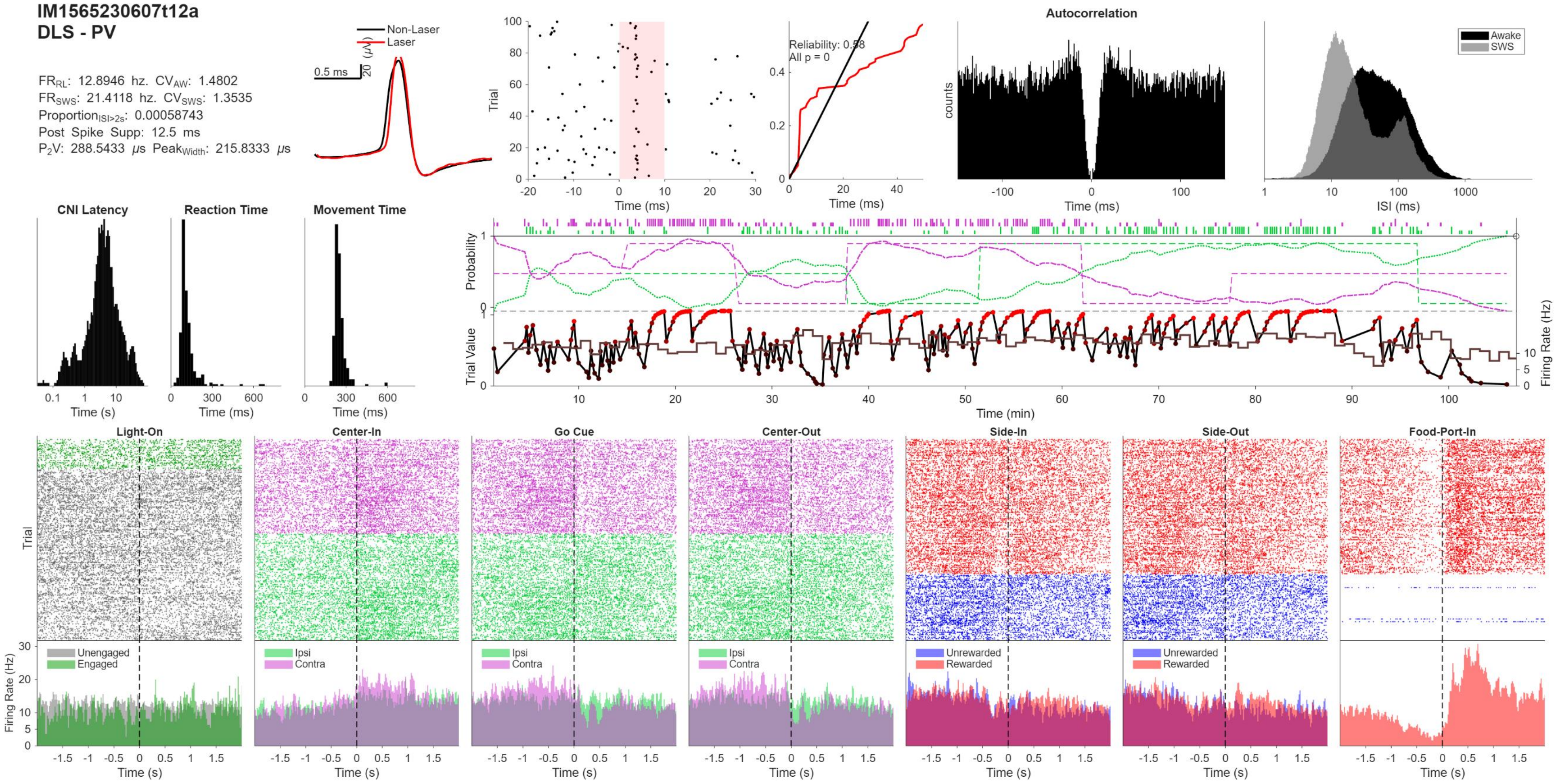
IM1565230607t12a
DLS - PV
FR$_{RL}$: 12.8946 hz. CV$_{AW}$: 1.4802
FR$_{SWS}$: 21.4118 hz. CV$_{SWS}$: 1.3535
Proportion$_{ISI>2s}$: 0.00058743
Post Spike Supp: 12.5 ms
P$_2$V: 288.5433 μs Peak$_{Width}$: 215.8333 μs
Non-Laser
Laser
0.5 ms
20 (μV)
Trial
Time (ms)
Reliability: 0.58
All p = 0
Autocorrelation
counts
Awake
SWS
ISI (ms)
CNI Latency
Reaction Time
Movement Time
Time (s)
Probability
Trial Value
Firing Rate (Hz)
Time (min)
Light-On
Center-In
Go Cue
Center-Out
Side-In
Side-Out
Food-Port-In
Unengaged
Engaged
Ipsi
Contra
Unrewarded
Rewarded

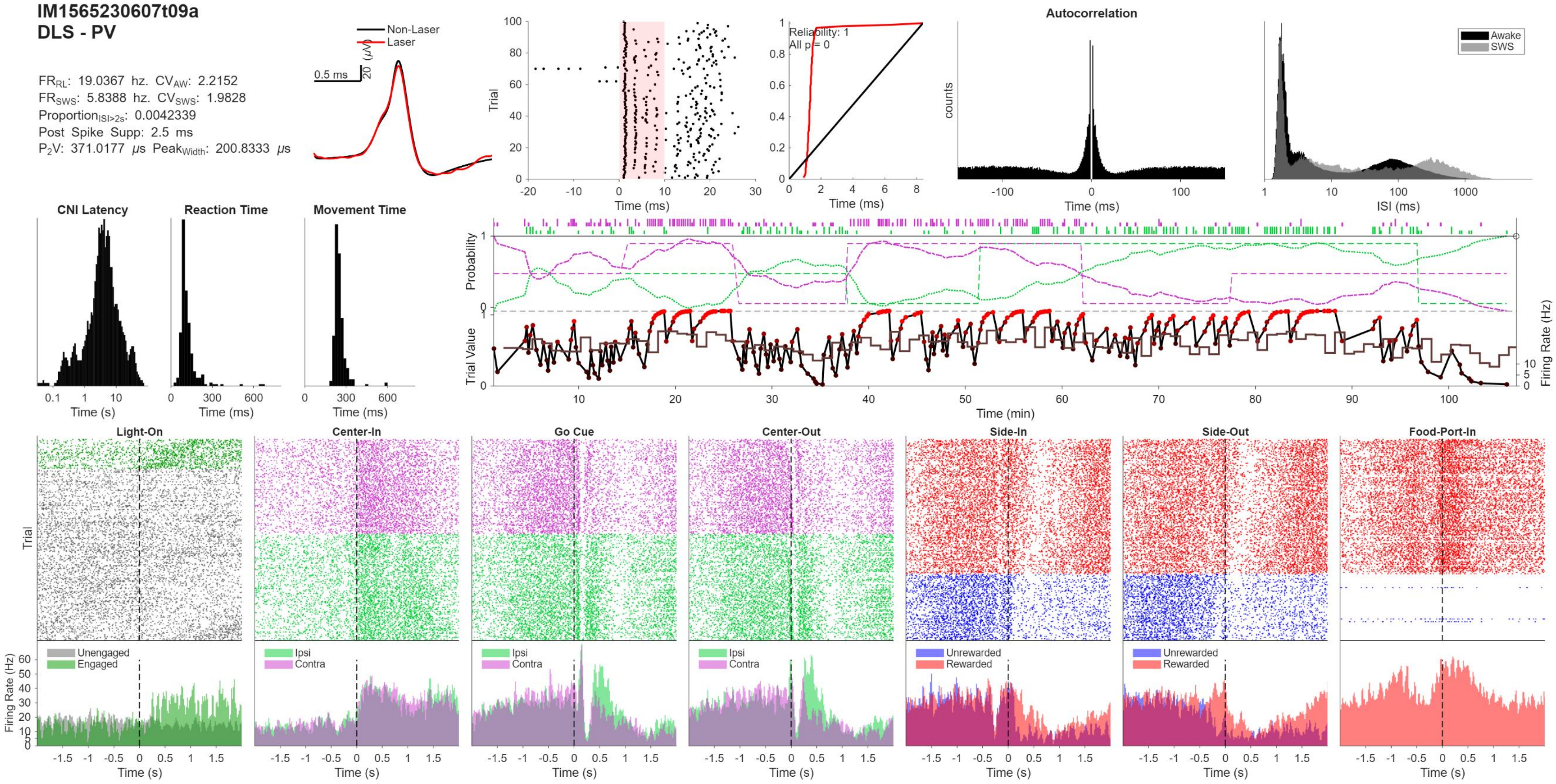
IM1565230607t09a
DLS - PV
FR_RL: 19.0367 hz. CV_AW: 2.2152
FR_SWS: 5.8388 hz. CV_SWS: 1.9828
Proportion_ISI>2s: 0.0042339
Post Spike Supp: 2.5 ms
P2V: 371.0177 μs Peak_Width: 200.8333 μs
Non-Laser
Laser
0.5 ms
20 (μV)
Trial
Time (ms)
Reliability: 1
All p = 0
Autocorrelation
counts
Awake
SWS
ISI (ms)
CNI Latency
Reaction Time
Movement Time
Time (s)
Probability
Trial Value
Firing Rate (Hz)
Time (min)
Light-On
Center-In
Go Cue
Center-Out
Side-In
Side-Out
Food-Port-In
Unengaged
Engaged
Ipsi
Contra
Unrewarded
Rewarded

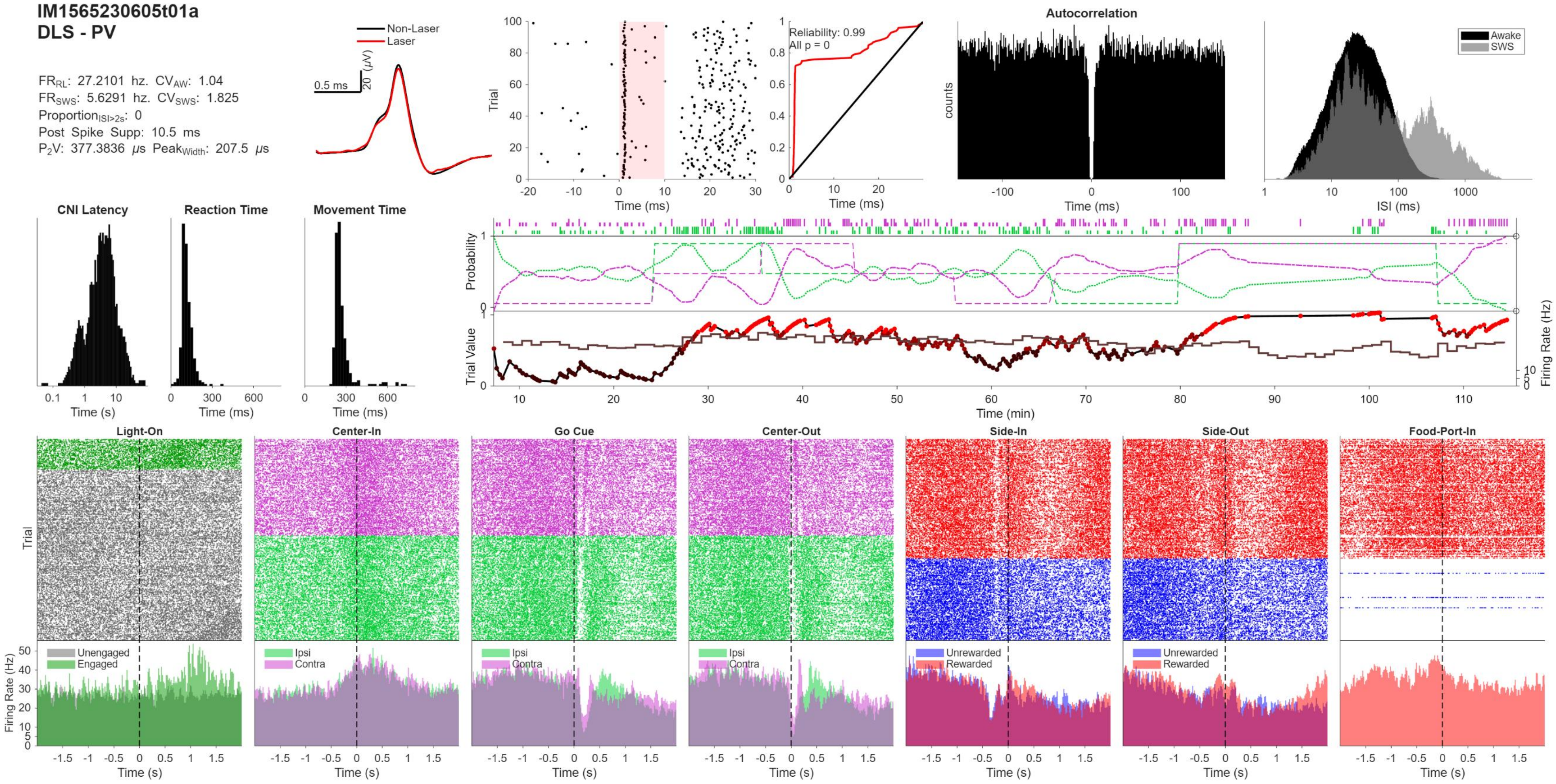
IM1565230605t01a
DLS - PV
FR_RL: 27.2101 hz. CV_AW: 1.04
FR_SWS: 5.6291 hz. CV_SWS: 1.825
Proportion_ISI>2s: 0
Post Spike Supp: 10.5 ms
P2V: 377.3836 μs Peak_Width: 207.5 μs
Non-Laser
Laser
0.5 ms
20 (μV)
Trial
Time (ms)
Reliability: 0.99
All p = 0
Autocorrelation
counts
Awake
SWS
ISI (ms)
CNI Latency
Reaction Time
Movement Time
Time (s)
Time (ms)
Probability
Trial Value
Firing Rate (Hz)
Time (min)
Light-On
Center-In
Go Cue
Center-Out
Side-In
Side-Out
Food-Port-In
Unengaged
Engaged
Ipsi
Contra
Unrewarded
Rewarded
Firing Rate (Hz)
Time (s)

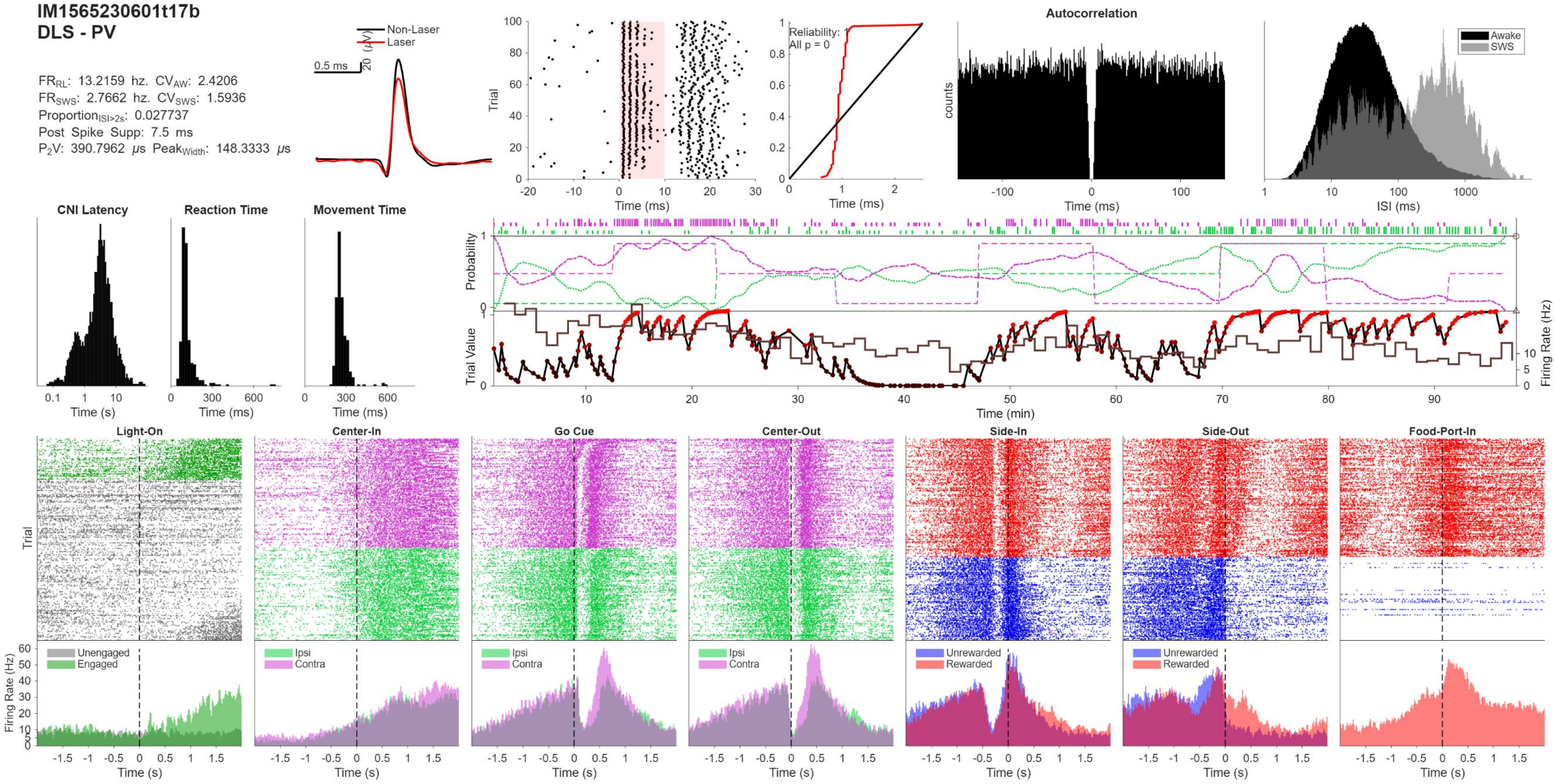
IM1565230601t17b
DLS - PV
FR_RL: 13.2159 hz. CV_AW: 2.4206
FR_SWS: 2.7662 hz. CV_SWS: 1.5936
Proportion_ISI>2s: 0.027737
Post Spike Supp: 7.5 ms
P2V: 390.7962 μs Peak_Width: 148.3333 μs
Non-Laser
Laser
0.5 ms
20 (μV)
Trial
Time (ms)
Reliability: 1
All p = 0
Time (ms)
Autocorrelation
counts
Time (ms)
Awake
SWS
ISI (ms)
CNI Latency
Reaction Time
Movement Time
Time (s)
Time (ms)
Time (ms)
Probability
Trial Value
Firing Rate (Hz)
Time (min)
Light-On
Center-In
Go Cue
Center-Out
Side-In
Side-Out
Food-Port-In
Trial
Firing Rate (Hz)
Unengaged
Engaged
Ipsi
Contra
Ipsi
Contra
Ipsi
Contra
Unrewarded
Rewarded
Unrewarded
Rewarded
Time (s)

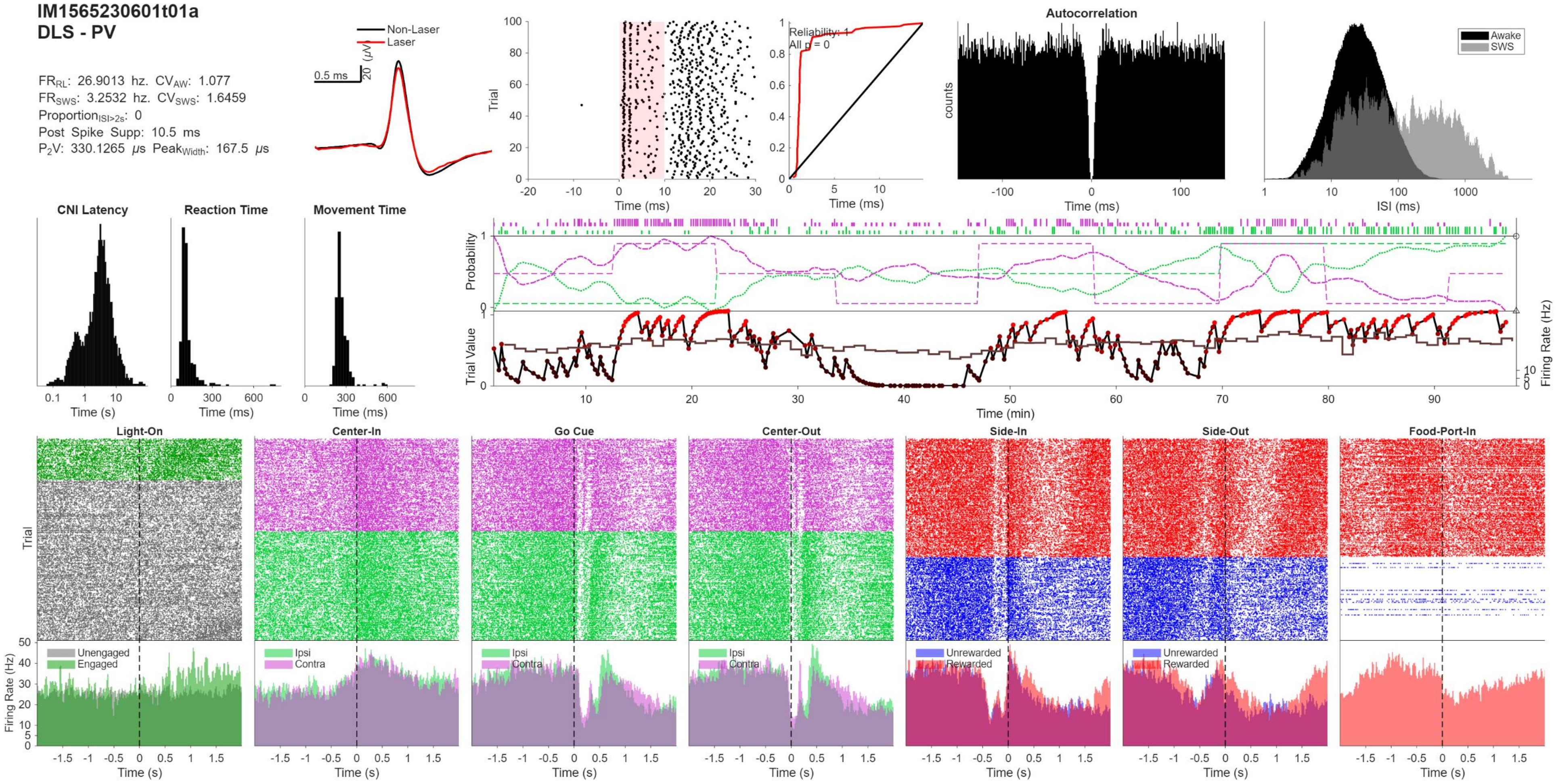
IM1565230601t01a
DLS - PV
FR_RL: 26.9013 hz. CV_AW: 1.077
FR_SWS: 3.2532 hz. CV_SWS: 1.6459
Proportion_ISI>2s: 0
Post Spike Supp: 10.5 ms
P2V: 330.1265 μs Peak_Width: 167.5 μs
Non-Laser
Laser
0.5 ms
20 (μV)
Trial
Time (ms)
Reliability: 1
All p = 0
Time (ms)
Autocorrelation
counts
Time (ms)
Awake
SWS
ISI (ms)
CNI Latency
Reaction Time
Movement Time
Time (s)
Time (ms)
Time (ms)
Probability
Trial Value
Firing Rate (Hz)
Time (min)
Light-On
Center-In
Go Cue
Center-Out
Side-In
Side-Out
Food-Port-In
Trial
Firing Rate (Hz)
Unengaged
Engaged
Ipsi
Contra
Ipsi
Contra
Ipsi
Contra
Unrewarded
Rewarded
Unrewarded
Rewarded
Time (s)

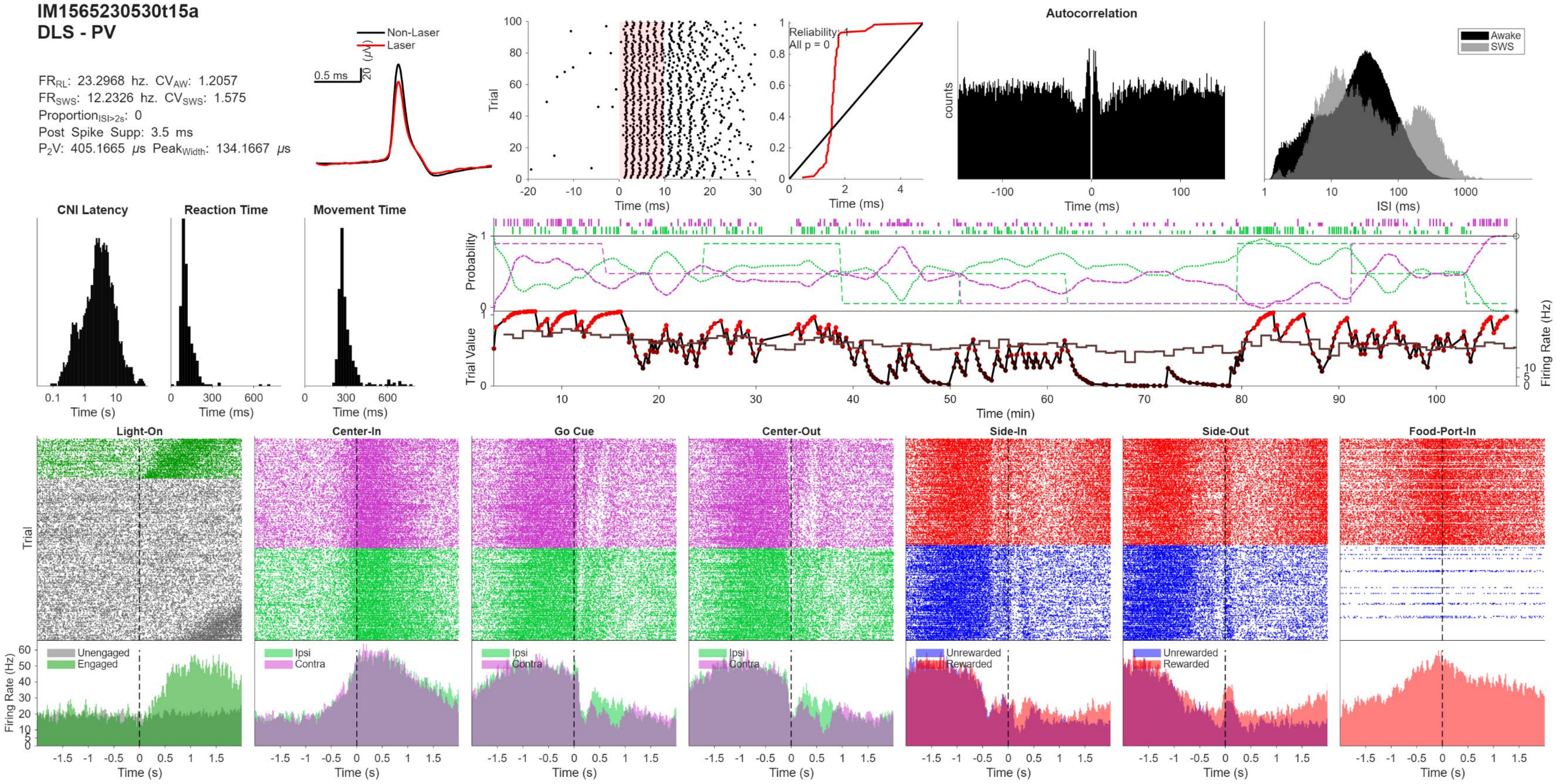

IM1565230530t15a
DLS - PV
FR_RL: 23.2968 hz. CV_AW: 1.2057
FR_SWS: 12.2326 hz. CV_SWS: 1.575
Proportion_ISI>2s: 0
Post Spike Supp: 3.5 ms
P2V: 405.1665 μs Peak_Width: 134.1667 μs
Non-Laser
Laser
0.5 ms
20 (μV)
Trial
Time (ms)
Reliability: 1
All p = 0
Autocorrelation
counts
Awake
SWS
ISI (ms)
CNI Latency
Reaction Time
Movement Time
Time (s)
Probability
Trial Value
Firing Rate (Hz)
Time (min)
Light-On
Center-In
Go Cue
Center-Out
Side-In
Side-Out
Food-Port-In
Unengaged
Engaged
Ipsi
Contra
Unrewarded
Rewarded

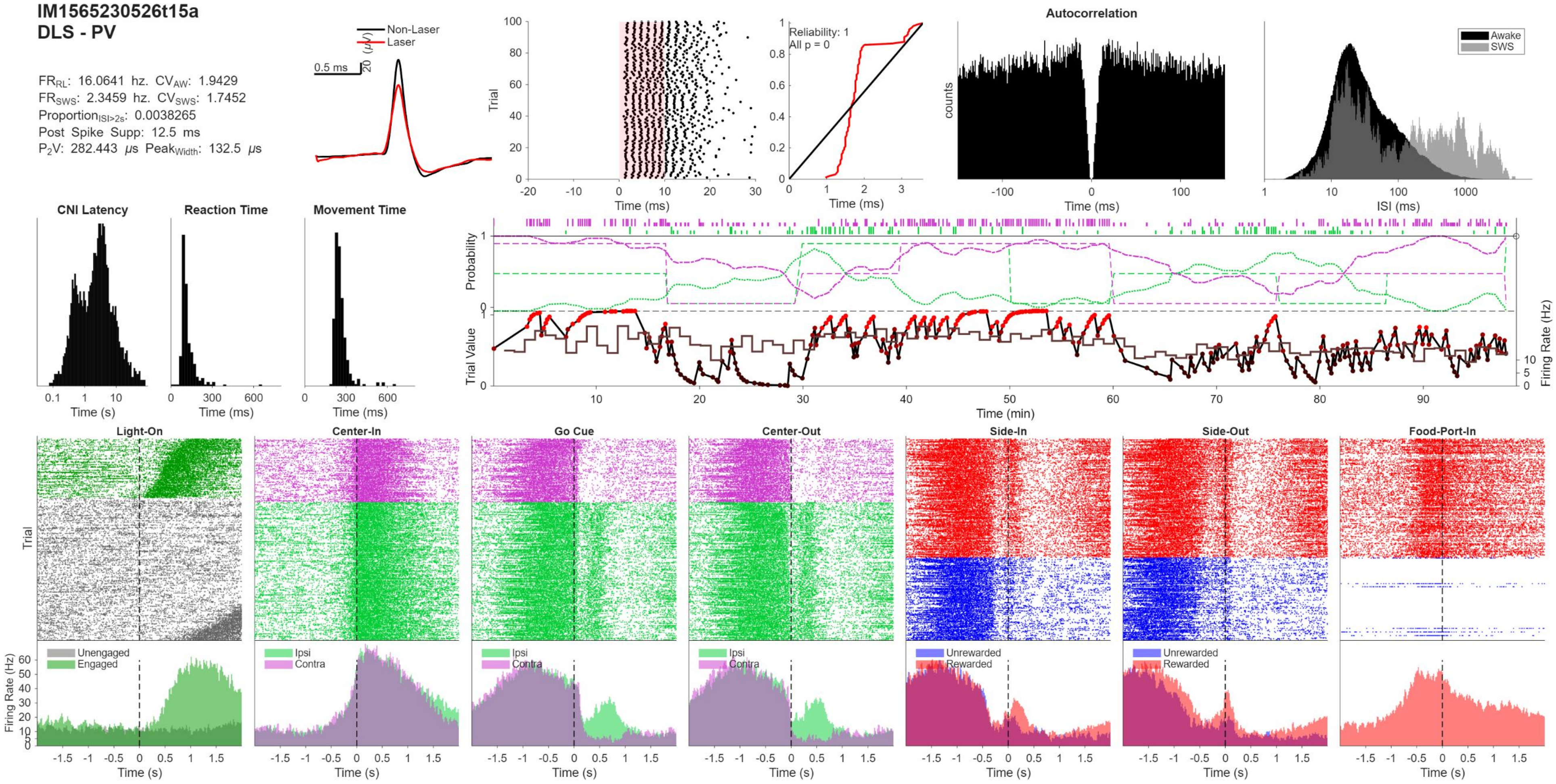
IM1565230526t15a
DLS - PV
FR_RL: 16.0641 hz. CV_AW: 1.9429
FR_SWS: 2.3459 hz. CV_SWS: 1.7452
Proportion_ISI>2s: 0.0038265
Post Spike Supp: 12.5 ms
P2V: 282.443 μs Peak_Width: 132.5 μs
Non-Laser
Laser
0.5 ms
20 (μV)
Trial
Time (ms)
Reliability: 1
All p = 0
Autocorrelation
counts
Awake
SWS
ISI (ms)
CNI Latency
Reaction Time
Movement Time
Time (s)
Probability
Trial Value
Firing Rate (Hz)
Time (min)
Light-On
Center-In
Go Cue
Center-Out
Side-In
Side-Out
Food-Port-In
Unengaged
Engaged
Ipsi
Contra
Unrewarded
Rewarded

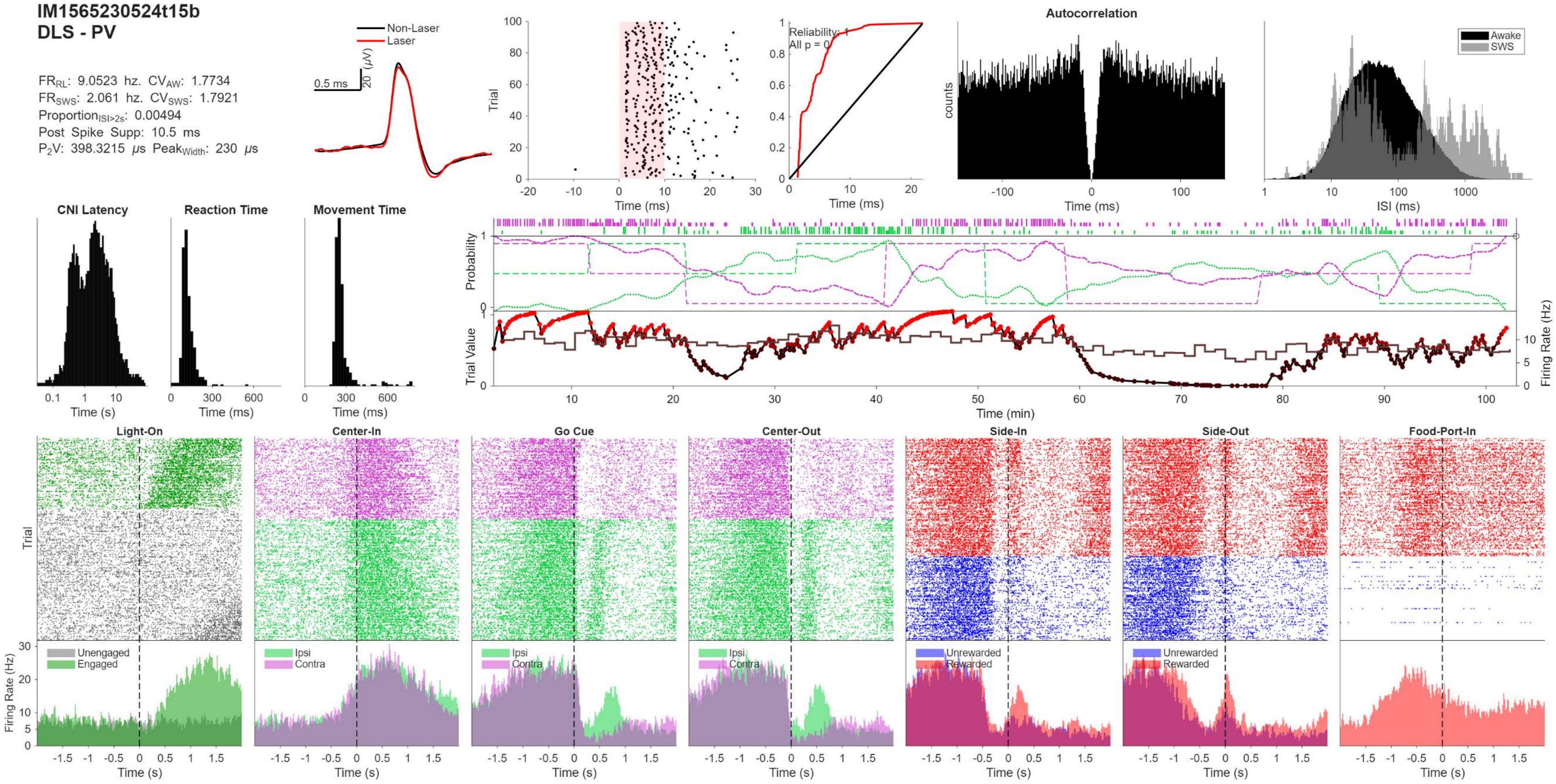
IM1565230524t15b
DLS - PV
FR_RL: 9.0523 hz. CV_AW: 1.7734
FR_SWS: 2.061 hz. CV_SWS: 1.7921
Proportion_ISI>2s: 0.00494
Post Spike Supp: 10.5 ms
P2V: 398.3215 μs Peak_Width: 230 μs
Non-Laser
Laser
0.5 ms
20 (μV)
Trial
Time (ms)
Reliability: 1
All p = 0
Autocorrelation
counts
Awake
SWS
ISI (ms)
CNI Latency
Reaction Time
Movement Time
Time (s)
Time (ms)
Probability
Trial Value
Firing Rate (Hz)
Time (min)
Light-On
Center-In
Go Cue
Center-Out
Side-In
Side-Out
Food-Port-In
Unengaged
Engaged
Ipsi
Contra
Unrewarded
Rewarded

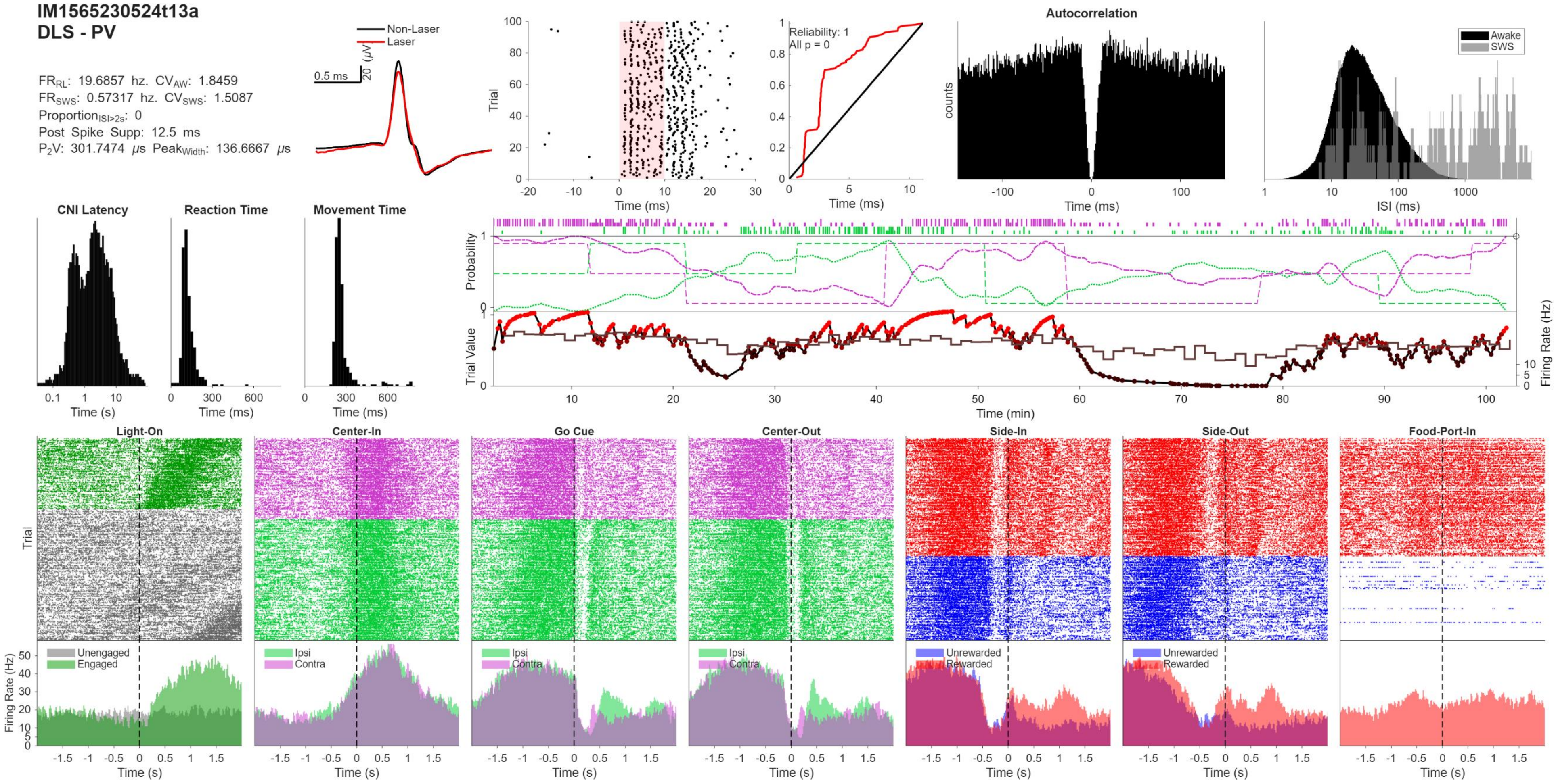

IM1565230524t13a
DLS - PV
FR_RL: 19.6857 hz. CV_AW: 1.8459
FR_SWS: 0.57317 hz. CV_SWS: 1.5087
Proportion_ISI>2s: 0
Post Spike Supp: 12.5 ms
P2V: 301.7474 μs Peak_Width: 136.6667 μs
Non-Laser
Laser
0.5 ms
20 (μV)
Trial
Time (ms)
Reliability: 1
All p = 0
Autocorrelation
counts
Awake
SWS
ISI (ms)
CNI Latency
Reaction Time
Movement Time
Time (s)
Probability
Trial Value
Firing Rate (Hz)
Time (min)
Light-On
Center-In
Go Cue
Center-Out
Side-In
Side-Out
Food-Port-In
Unengaged
Engaged
Ipsi
Contra
Unrewarded
Rewarded

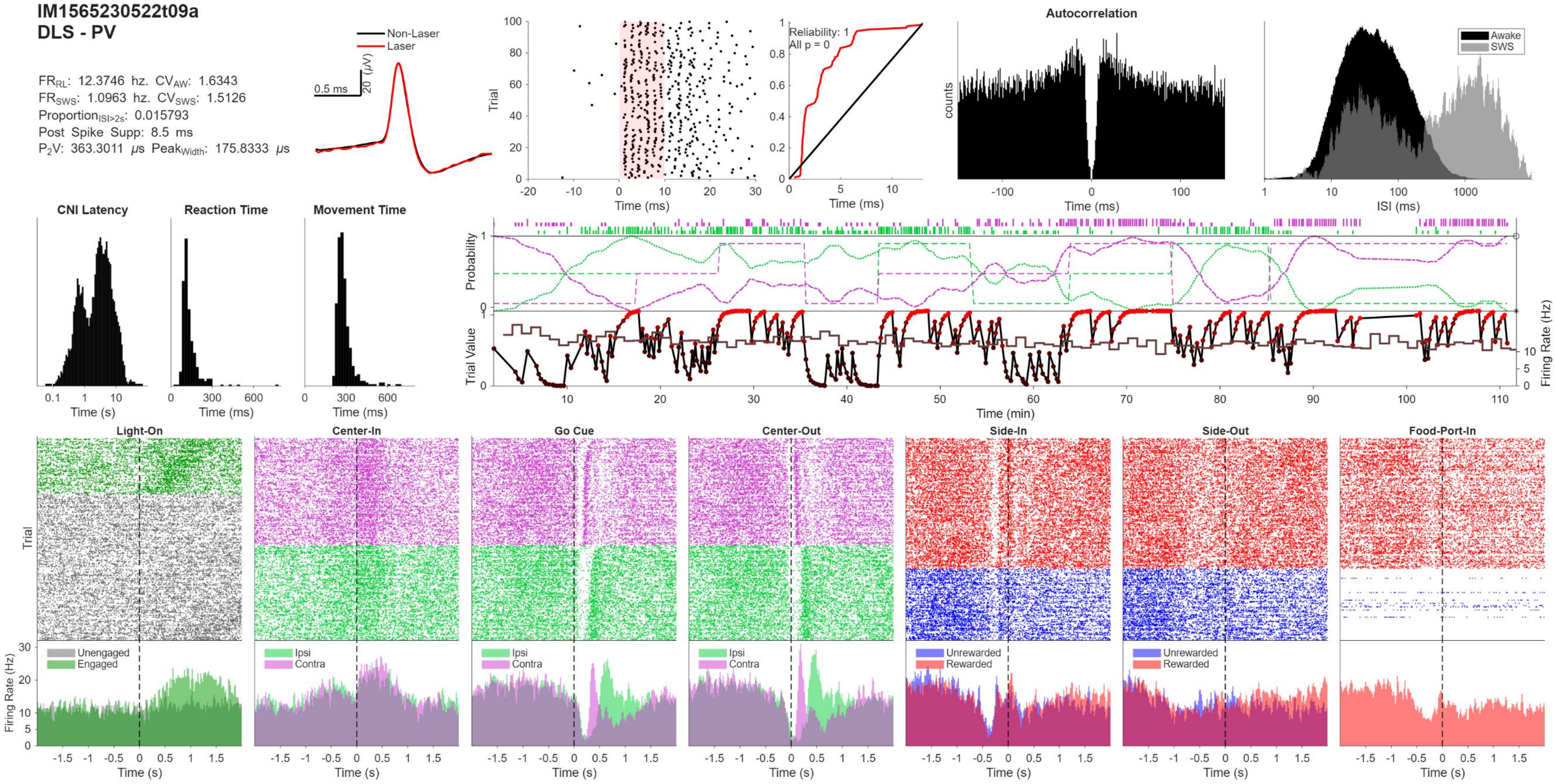
IM1565230522t09a
DLS - PV
FR_RL: 12.3746 hz. CV_AW: 1.6343
FR_SWS: 1.0963 hz. CV_SWS: 1.5126
Proportion_ISI>2s: 0.015793
Post Spike Supp: 8.5 ms
P2V: 363.3011 μs Peak_Width: 175.8333 μs
Non-Laser
Laser
0.5 ms
20 (μV)
Trial
Time (ms)
Reliability: 1
All p = 0
Autocorrelation
counts
Awake
SWS
ISI (ms)
CNI Latency
Reaction Time
Movement Time
Time (s)
Time (ms)
Probability
Trial Value
Firing Rate (Hz)
Time (min)
Light-On
Center-In
Go Cue
Center-Out
Side-In
Side-Out
Food-Port-In
Unengaged
Engaged
Ipsi
Contra
Unrewarded
Rewarded
Firing Rate (Hz)

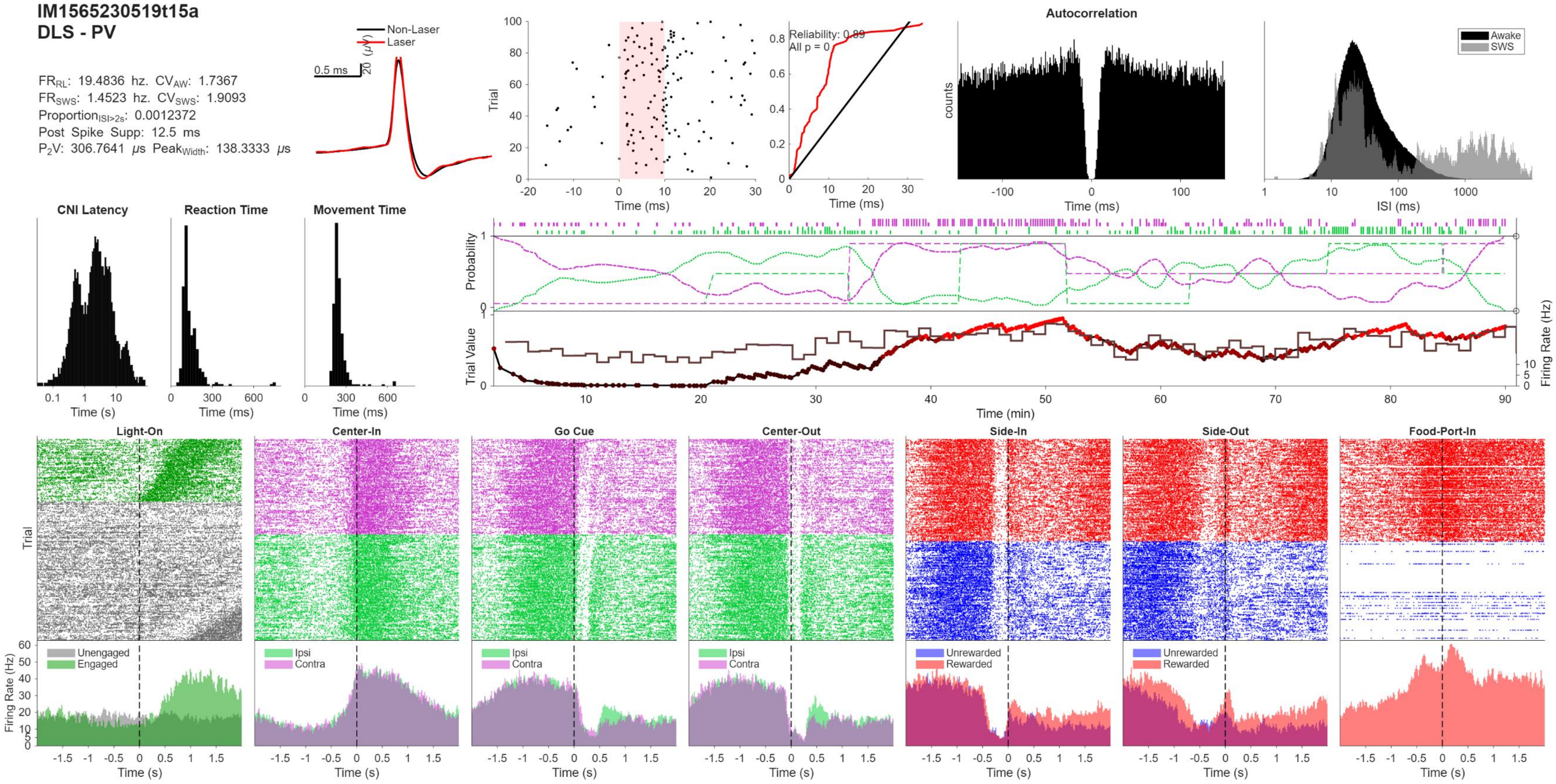
IM1565230519t15a
DLS - PV
FR_RL: 19.4836 hz. CV_AW: 1.7367
FR_SWS: 1.4523 hz. CV_SWS: 1.9093
Proportion_ISI>2s: 0.0012372
Post Spike Supp: 12.5 ms
P2V: 306.7641 μs Peak_Width: 138.3333 μs
Non-Laser
Laser
0.5 ms
20 (μV)
Trial
Time (ms)
Reliability: 0.89
All p = 0
Autocorrelation
counts
Awake
SWS
ISI (ms)
CNI Latency
Reaction Time
Movement Time
Time (s)
Probability
Trial Value
Firing Rate (Hz)
Time (min)
Light-On
Center-In
Go Cue
Center-Out
Side-In
Side-Out
Food-Port-In
Unengaged
Engaged
Ipsi
Contra
Unrewarded
Rewarded

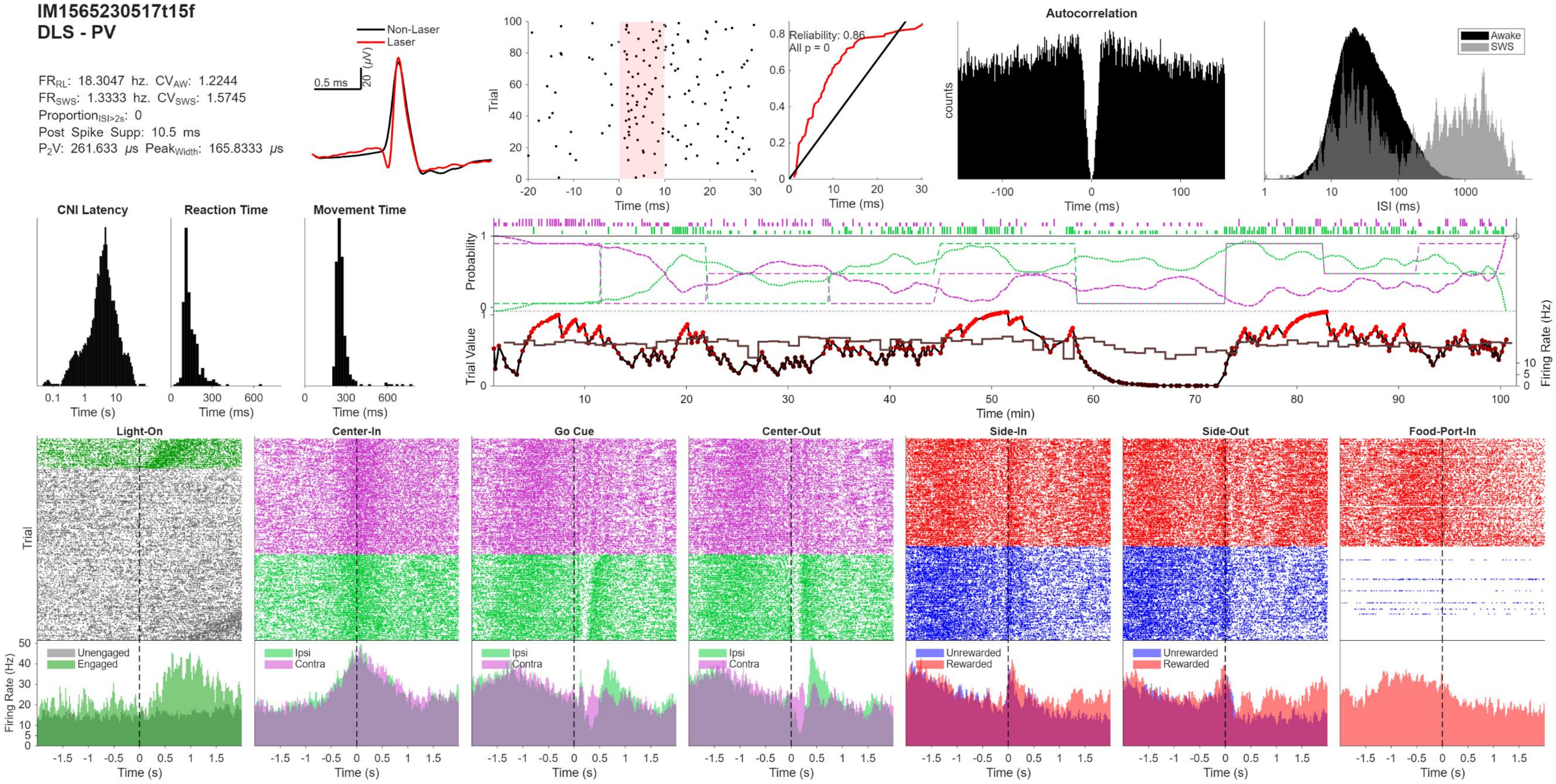
IM1565230517t15f
DLS - PV
FR$_{RL}$: 18.3047 hz. CV$_{AW}$: 1.2244
FR$_{SWS}$: 1.3333 hz. CV$_{SWS}$: 1.5745
Proportion$_{ISI>2s}$: 0
Post Spike Supp: 10.5 ms
P$_2$V: 261.633 μs Peak$_{Width}$: 165.8333 μs
Non-Laser
Laser
0.5 ms
20 (μV)
Trial
Time (ms)
Reliability: 0.86
All p = 0
Autocorrelation
counts
Awake
SWS
ISI (ms)
CNI Latency
Reaction Time
Movement Time
Time (s)
Probability
Trial Value
Firing Rate (Hz)
Time (min)
Light-On
Center-In
Go Cue
Center-Out
Side-In
Side-Out
Food-Port-In
Unengaged
Engaged
Ipsi
Contra
Unrewarded
Rewarded
Firing Rate (Hz)

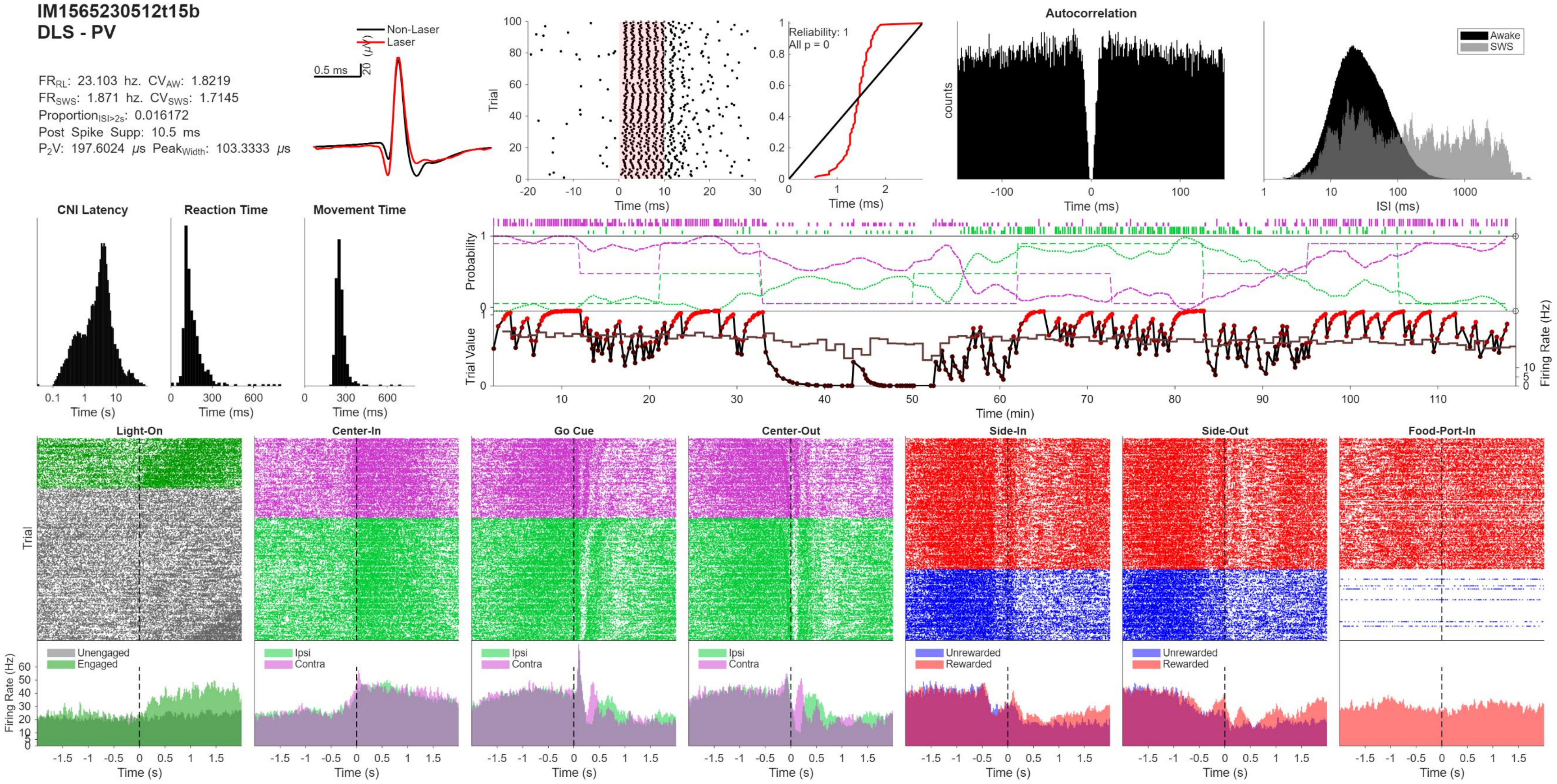
IM1565230512t15b
DLS - PV
FR_RL: 23.103 hz. CV_AW: 1.8219
FR_SWS: 1.871 hz. CV_SWS: 1.7145
Proportion_ISI>2s: 0.016172
Post Spike Supp: 10.5 ms
P2V: 197.6024 μs Peak_Width: 103.3333 μs
Non-Laser
Laser
0.5 ms
20 (μV)
Trial
Time (ms)
Reliability: 1
All p = 0
Autocorrelation
counts
Awake
SWS
ISI (ms)
CNI Latency
Reaction Time
Movement Time
Time (s)
Probability
Trial Value
Firing Rate (Hz)
Time (min)
Light-On
Center-In
Go Cue
Center-Out
Side-In
Side-Out
Food-Port-In
Unengaged
Engaged
Ipsi
Contra
Unrewarded
Rewarded

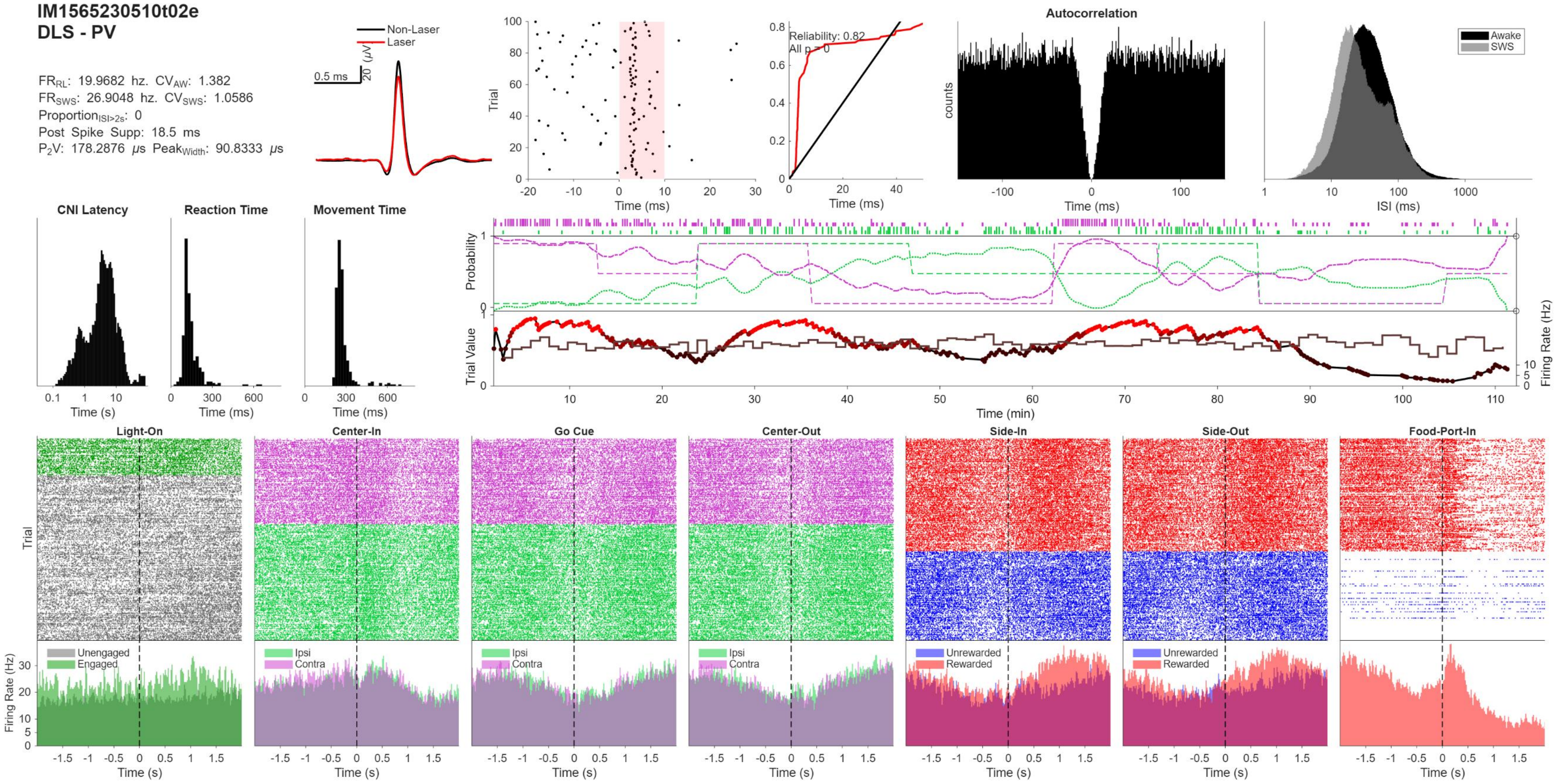
IM1565230510t02e
DLS - PV
FRRL: 19.9682 hz. CVAW: 1.382
FRSWS: 26.9048 hz. CVSWS: 1.0586
ProportionISI>2s: 0
Post Spike Supp: 18.5 ms
P2V: 178.2876 μs PeakWidth: 90.8333 μs
Non-Laser
Laser
0.5 ms
20 (μV)
Trial
Time (ms)
Reliability: 0.82
All p = 0
Autocorrelation
counts
Awake
SWS
ISI (ms)
CNI Latency
Reaction Time
Movement Time
Time (s)
Probability
Trial Value
Firing Rate (Hz)
Time (min)
Light-On
Center-In
Go Cue
Center-Out
Side-In
Side-Out
Food-Port-In
Unengaged
Engaged
Ipsi
Contra
Unrewarded
Rewarded

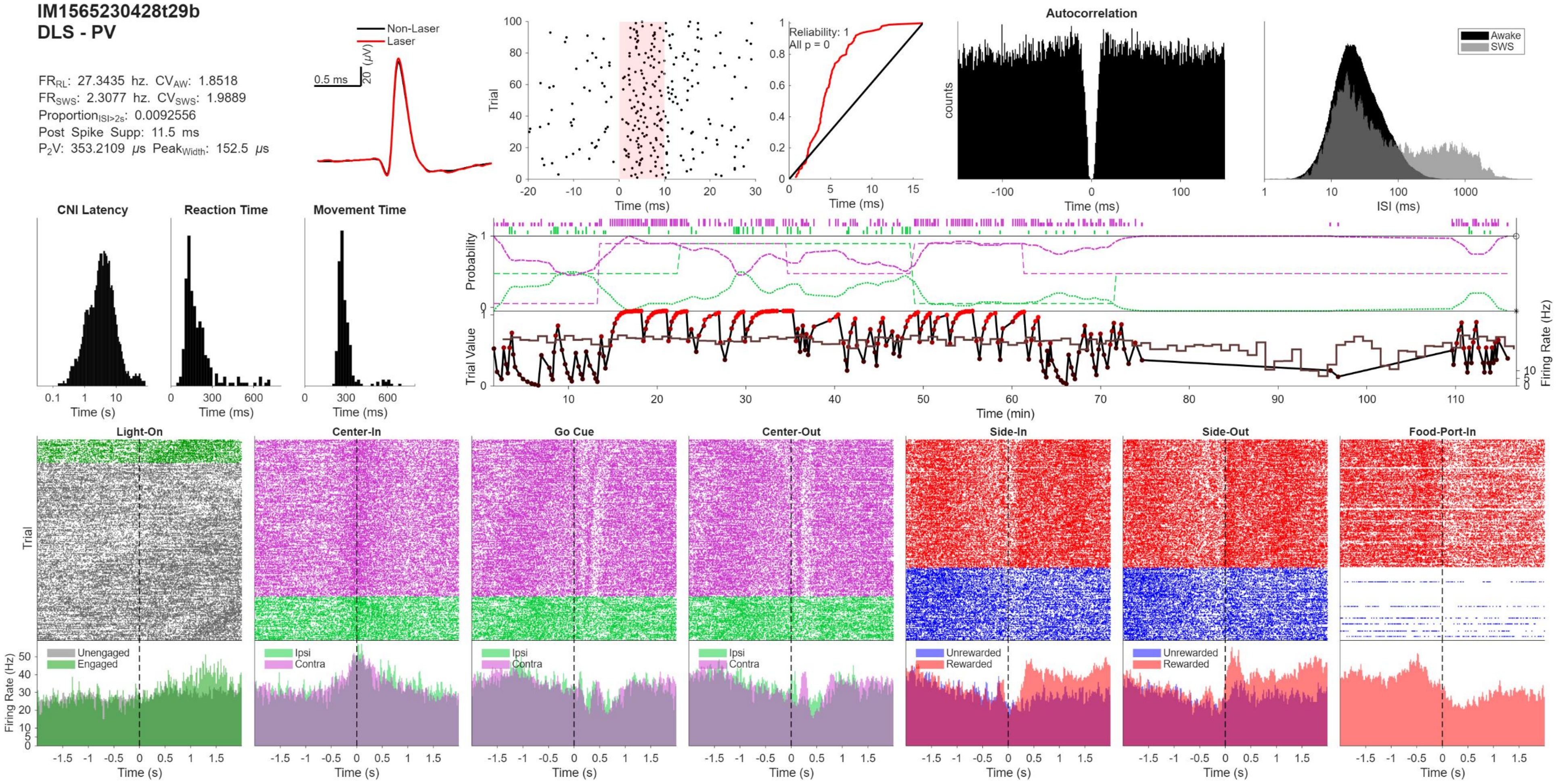
IM1565230428t29b
DLS - PV
FR_RL: 27.3435 hz. CV_AW: 1.8518
FR_SWS: 2.3077 hz. CV_SWS: 1.9889
Proportion_ISI>2s: 0.0092556
Post Spike Supp: 11.5 ms
P2V: 353.2109 μs Peak_Width: 152.5 μs
Non-Laser
Laser
0.5 ms
20 (μV)
Trial
Time (ms)
Reliability: 1
All p = 0
Autocorrelation
counts
Awake
SWS
ISI (ms)
CNI Latency
Reaction Time
Movement Time
Time (s)
Probability
Trial Value
Firing Rate (Hz)
Time (min)
Light-On
Center-In
Go Cue
Center-Out
Side-In
Side-Out
Food-Port-In
Unengaged
Engaged
Ipsi
Contra
Unrewarded
Rewarded

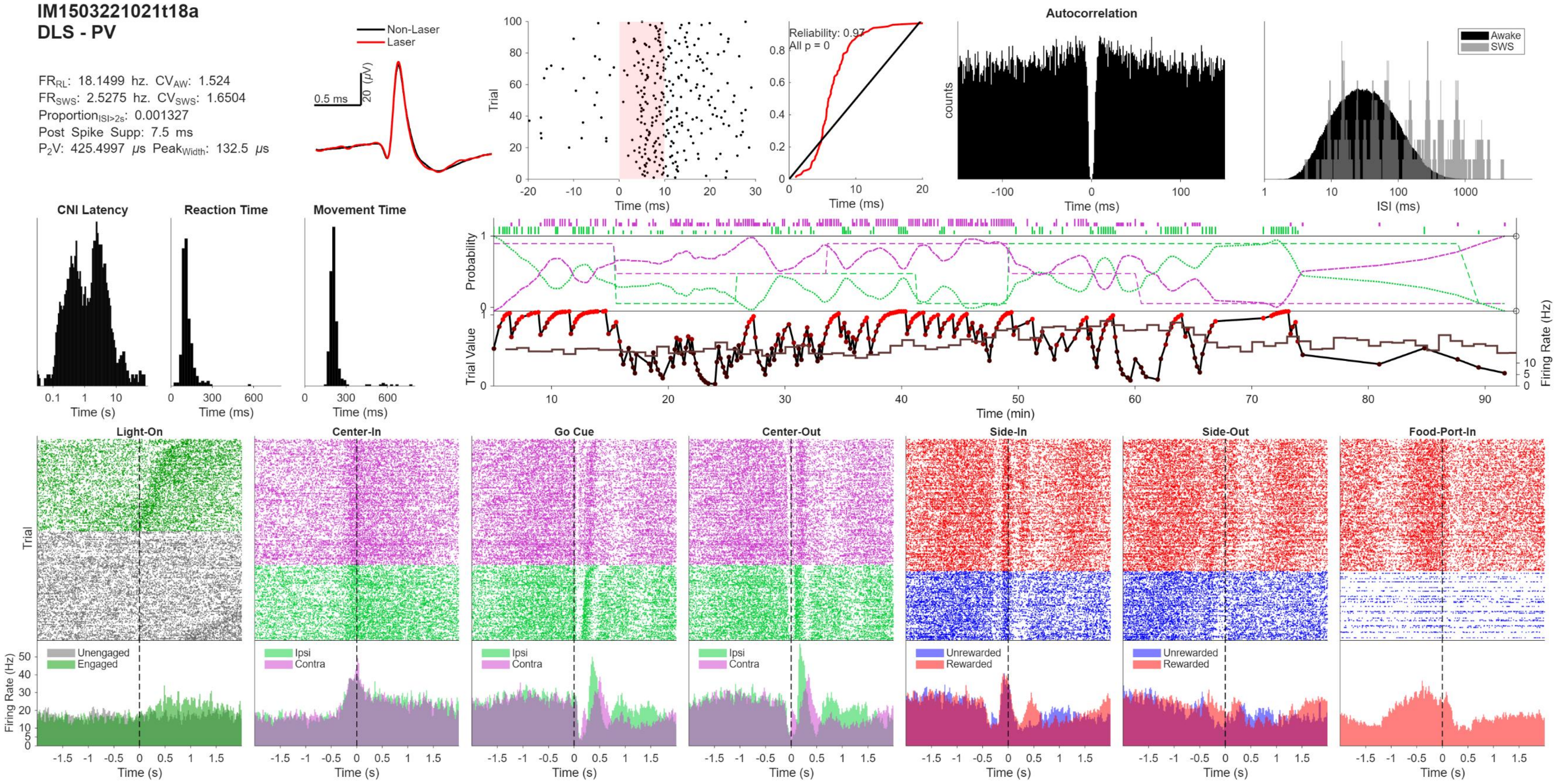
IM1503221021t18a
DLS - PV
FR_RL: 18.1499 hz. CV_AW: 1.524
FR_SWS: 2.5275 hz. CV_SWS: 1.6504
Proportion_ISI>2s: 0.001327
Post Spike Supp: 7.5 ms
P2V: 425.4997 μs Peak_Width: 132.5 μs
Non-Laser
Laser
0.5 ms
20 (μV)
Trial
Time (ms)
Reliability: 0.97
All p = 0
Autocorrelation
counts
Awake
SWS
ISI (ms)
CNI Latency
Reaction Time
Movement Time
Time (s)
Probability
Trial Value
Firing Rate (Hz)
Time (min)
Light-On
Center-In
Go Cue
Center-Out
Side-In
Side-Out
Food-Port-In
Unengaged
Engaged
Ipsi
Contra
Unrewarded
Rewarded

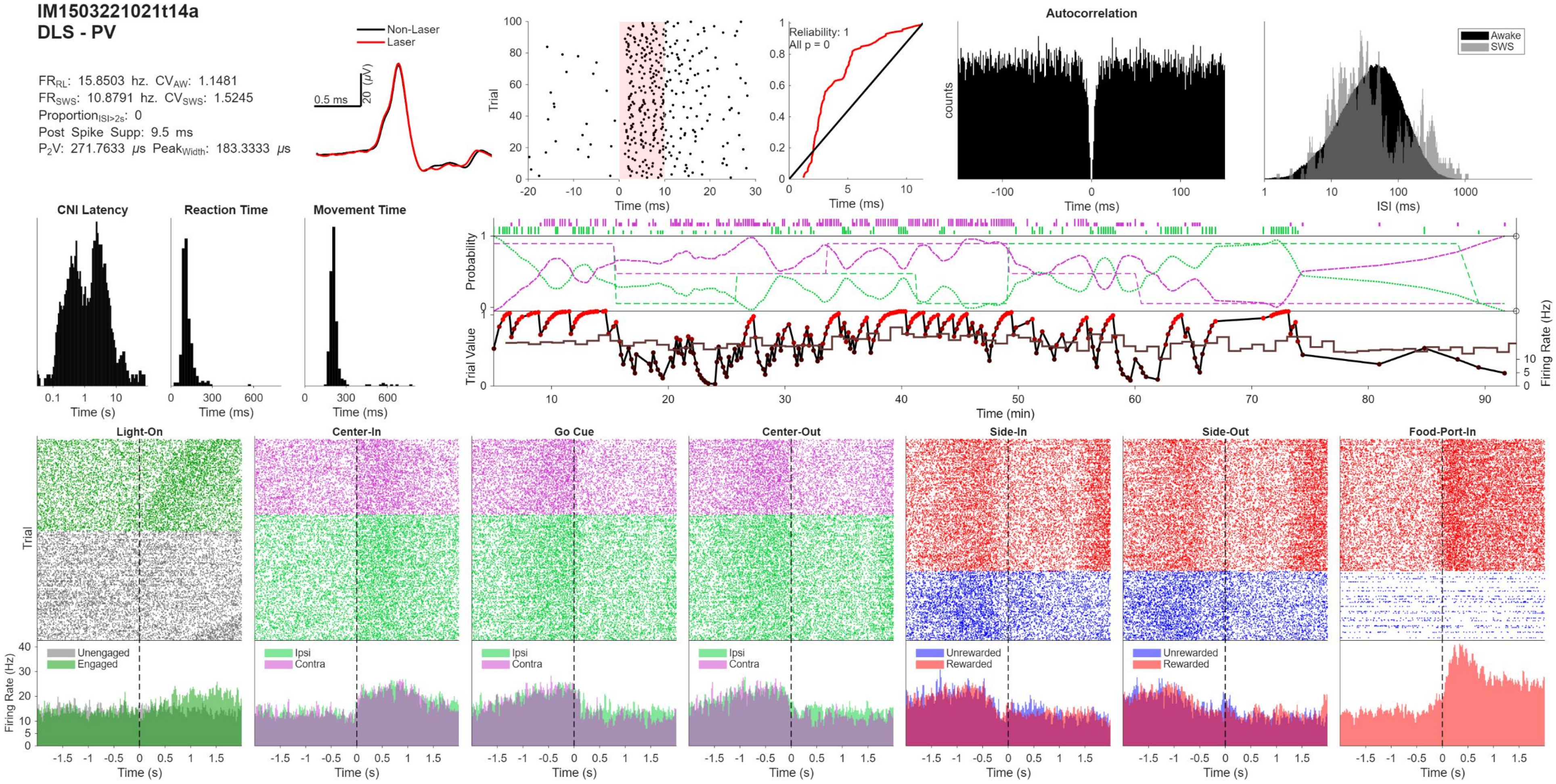
IM1503221021t14a
DLS - PV
FRRL: 15.8503 hz. CVAW: 1.1481
FRSWS: 10.8791 hz. CVSWS: 1.5245
ProportionISI>2s: 0
Post Spike Supp: 9.5 ms
P2V: 271.7633 μs PeakWidth: 183.3333 μs
Non-Laser
Laser
0.5 ms
20 (μV)
Trial
Time (ms)
Reliability: 1
All p = 0
Time (ms)
Autocorrelation
counts
Time (ms)
Awake
SWS
ISI (ms)
CNI Latency
Time (s)
Reaction Time
Time (ms)
Movement Time
Time (ms)
Probability
Trial Value
Firing Rate (Hz)
Time (min)
Light-On
Center-In
Go Cue
Center-Out
Side-In
Side-Out
Food-Port-In
Trial
Firing Rate (Hz)
Unengaged
Engaged
Ipsi
Contra
Unrewarded
Rewarded
Time (s)

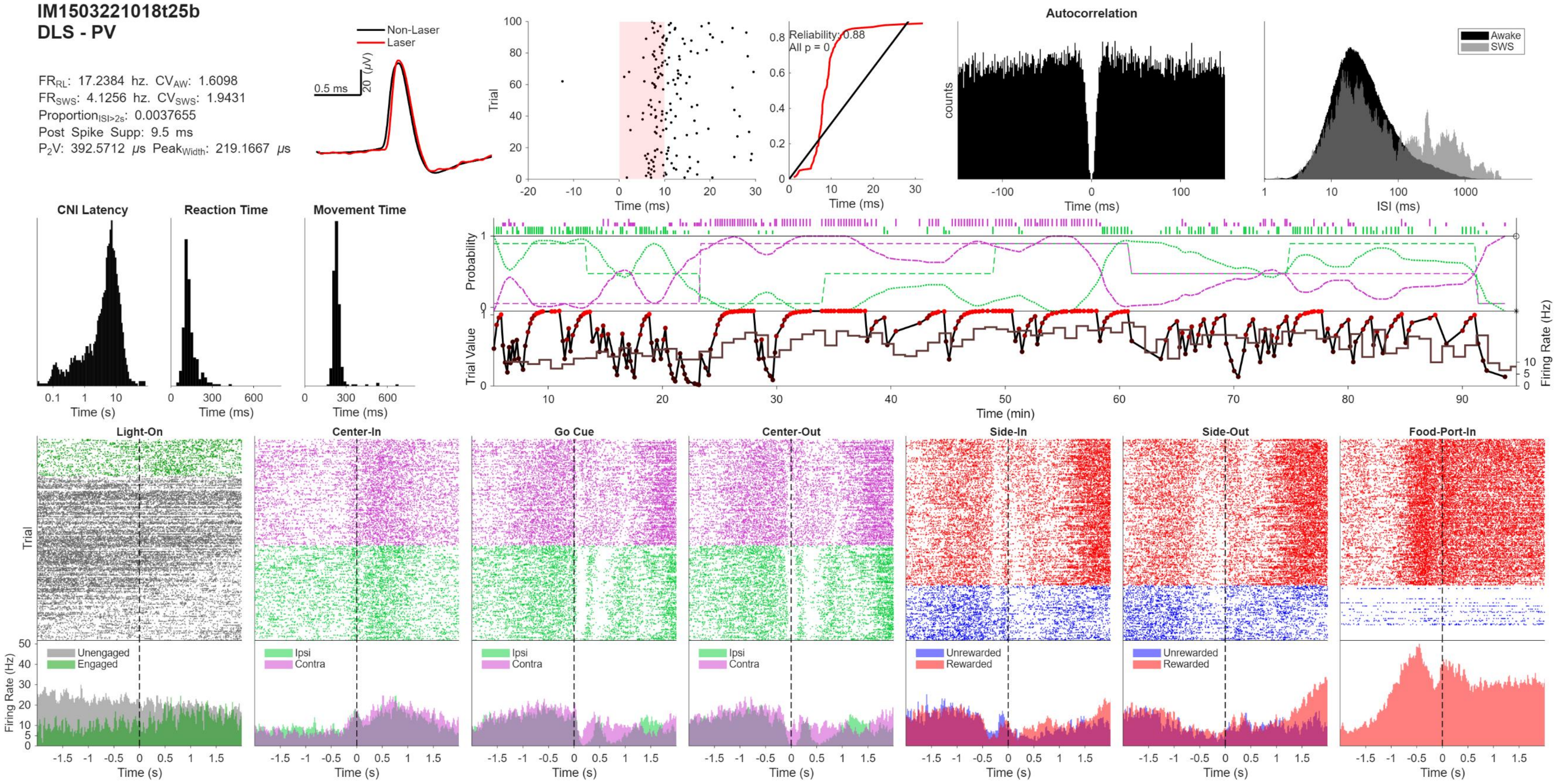
IM1503221018t25b
DLS - PV
$FR_{RL}$: 17.2384 hz. $CV_{AW}$: 1.6098
$FR_{SWS}$: 4.1256 hz. $CV_{SWS}$: 1.9431
$Proportion_{ISI>2s}$: 0.0037655
Post Spike Supp: 9.5 ms
$P_2V$: 392.5712 $\mu s$ $Peak_{Width}$: 219.1667 $\mu s$
Non-Laser
Laser
0.5 ms
20 ($\mu V$)
Trial
Time (ms)
Reliability: 0.88
All p = 0
Autocorrelation
counts
Time (ms)
Awake
SWS
ISI (ms)
CNI Latency
Time (s)
Reaction Time
Time (ms)
Movement Time
Time (ms)
Probability
Trial Value
Firing Rate (Hz)
Time (min)
Light-On
Center-In
Go Cue
Center-Out
Side-In
Side-Out
Food-Port-In
Trial
Firing Rate (Hz)
Unengaged
Engaged
Ipsi
Contra
Unrewarded
Rewarded
Time (s)

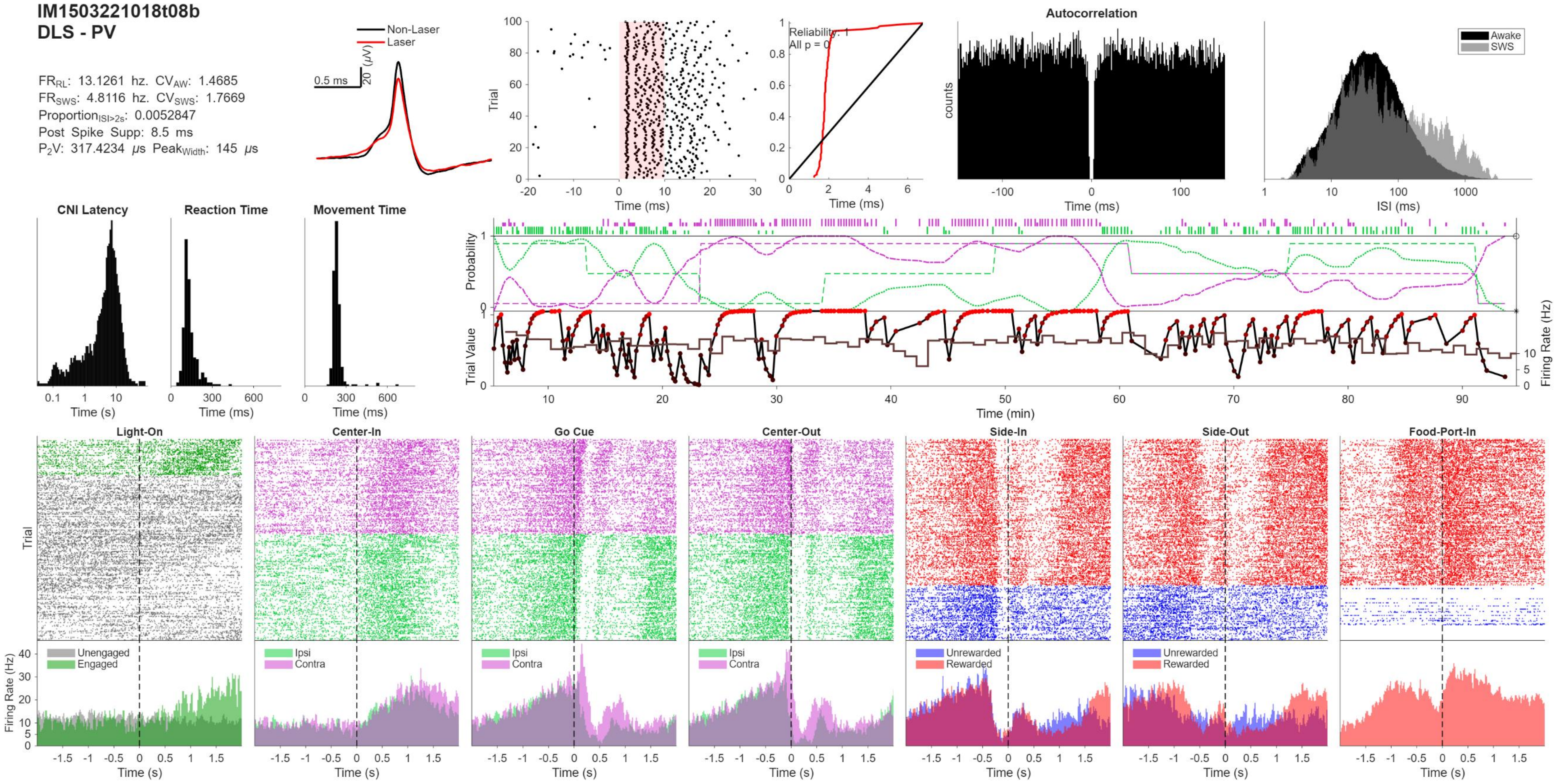
IM1503221018t08b
DLS - PV
FR_RL: 13.1261 hz. CV_AW: 1.4685
FR_SWS: 4.8116 hz. CV_SWS: 1.7669
Proportion_ISI>2s: 0.0052847
Post Spike Supp: 8.5 ms
P2V: 317.4234 μs Peak_Width: 145 μs
Non-Laser
Laser
0.5 ms
20 (μV)
Trial
Time (ms)
Reliability: 1
All p = 0
Autocorrelation
counts
Awake
SWS
ISI (ms)
CNI Latency
Reaction Time
Movement Time
Time (s)
Time (ms)
Probability
Trial Value
Firing Rate (Hz)
Time (min)
Light-On
Center-In
Go Cue
Center-Out
Side-In
Side-Out
Food-Port-In
Unengaged
Engaged
Ipsi
Contra
Unrewarded
Rewarded

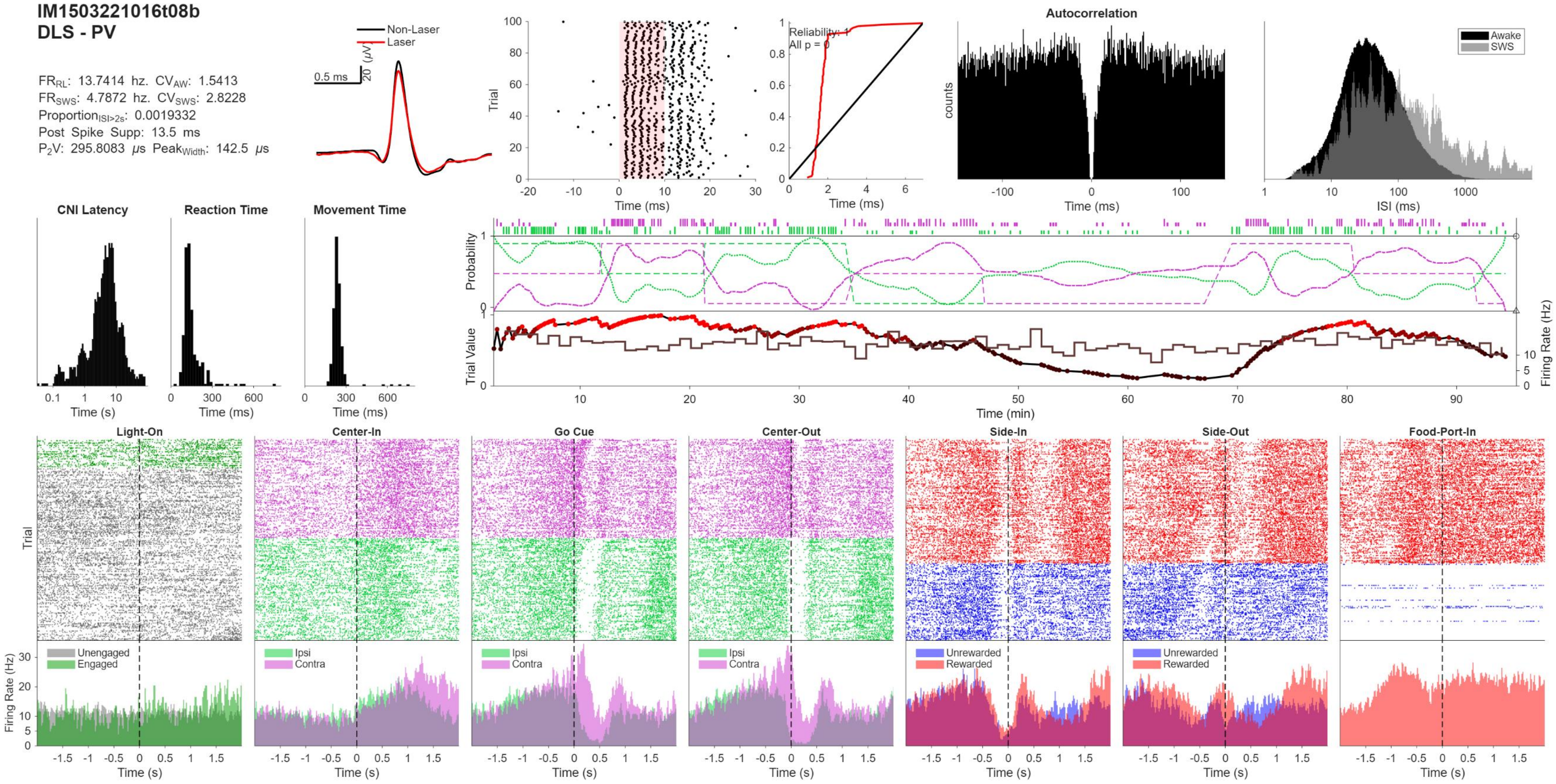
IM1503221016t08b
DLS - PV
FR_RL: 13.7414 hz. CV_AW: 1.5413
FR_SWS: 4.7872 hz. CV_SWS: 2.8228
Proportion_ISI>2s: 0.0019332
Post Spike Supp: 13.5 ms
P2V: 295.8083 μs Peak_Width: 142.5 μs
Non-Laser
Laser
0.5 ms
20 (μV)
Reliability: 1
All p = 0
Autocorrelation
Awake
SWS
CNI Latency
Reaction Time
Movement Time
Light-On
Center-In
Go Cue
Center-Out
Side-In
Side-Out
Food-Port-In
Unengaged
Engaged
Ipsi
Contra
Unrewarded
Rewarded
Trial
counts
Probability
Trial Value
Firing Rate (Hz)
Time (ms)
ISI (ms)
Time (s)
Time (min)

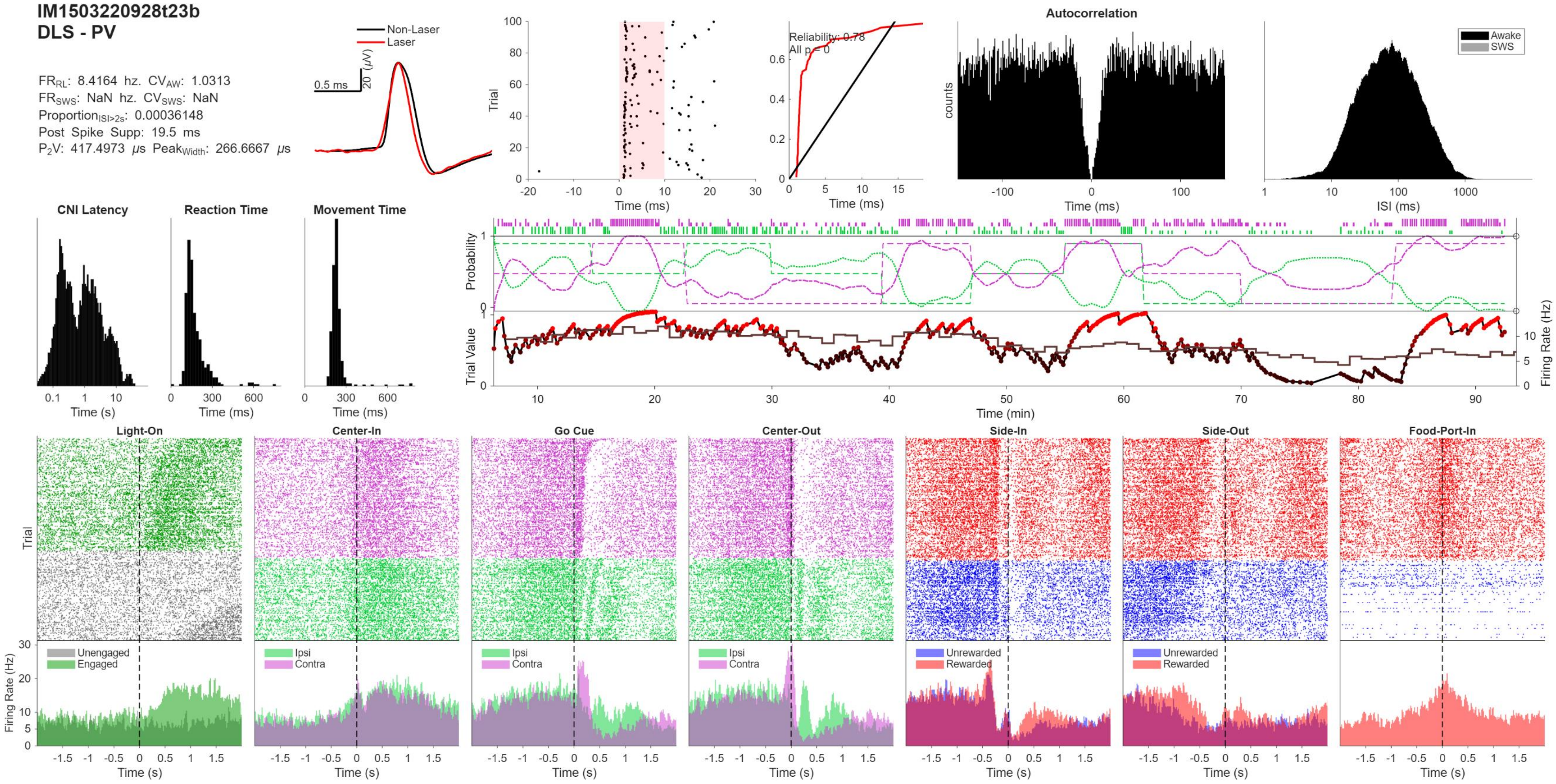

IM1503220928t23b
DLS - PV
FR_RL: 8.4164 hz. CV_AW: 1.0313
FR_SWS: NaN hz. CV_SWS: NaN
Proportion_ISI>2s: 0.00036148
Post Spike Supp: 19.5 ms
P2V: 417.4973 μs Peak_Width: 266.6667 μs
Non-Laser
Laser
0.5 ms
20 (μV)
Trial
Time (ms)
Reliability: 0.78
All p = 0
Autocorrelation
counts
Awake
SWS
ISI (ms)
CNI Latency
Reaction Time
Movement Time
Time (s)
Time (ms)
Probability
Trial Value
Firing Rate (Hz)
Time (min)
Light-On
Center-In
Go Cue
Center-Out
Side-In
Side-Out
Food-Port-In
Unengaged
Engaged
Ipsi
Contra
Unrewarded
Rewarded

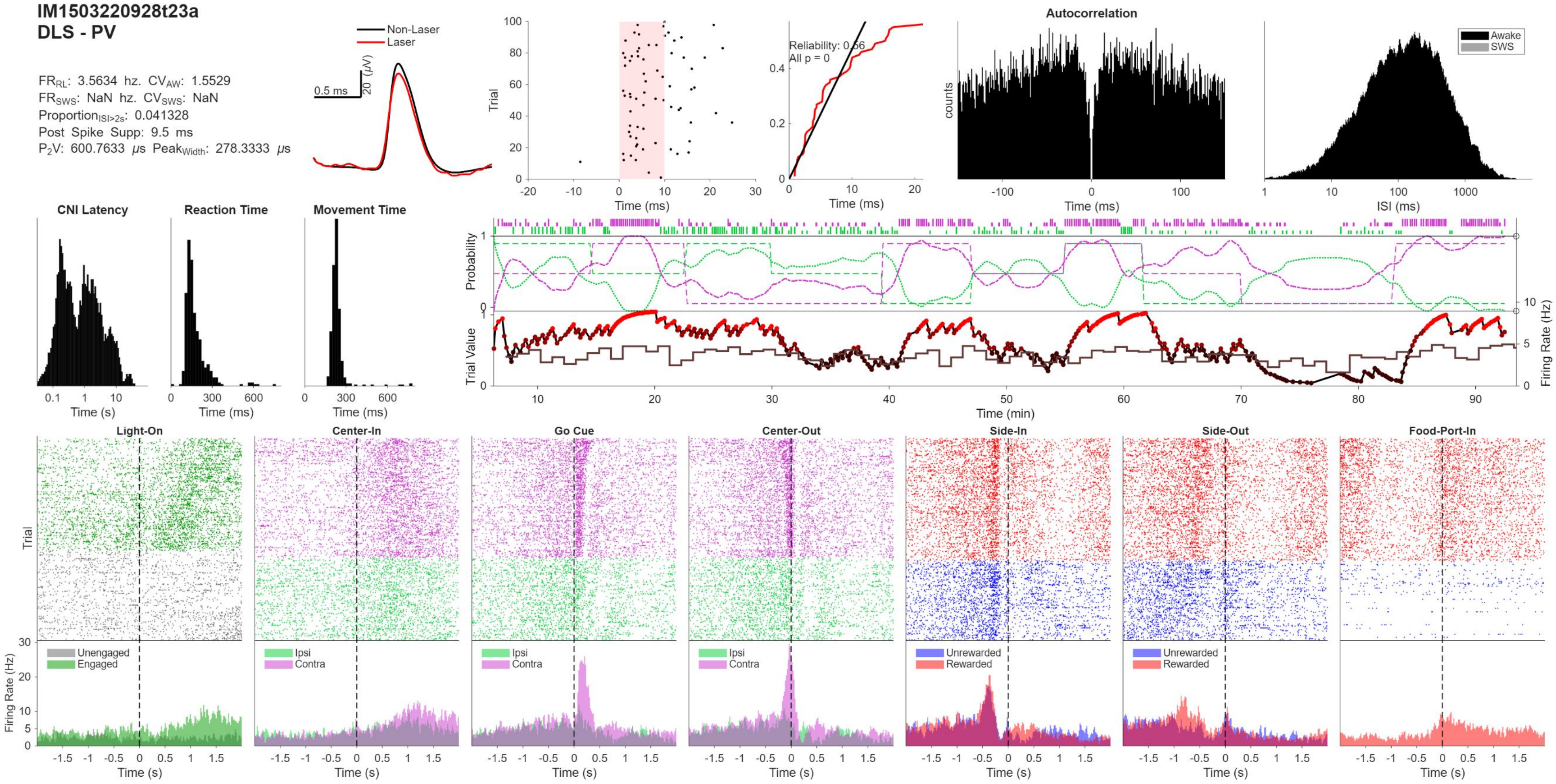

IM1503220928t23a
DLS - PV
FRRL: 3.5634 hz. CVAW: 1.5529
FRSWS: NaN hz. CVSWS: NaN
ProportionISI>2s: 0.041328
Post Spike Supp: 9.5 ms
P2V: 600.7633 μs PeakWidth: 278.3333 μs
Non-Laser
Laser
0.5 ms
20 (μV)
Trial
Time (ms)
Reliability: 0.56
All p = 0
Time (ms)
Autocorrelation
counts
Time (ms)
Awake
SWS
ISI (ms)
CNI Latency
Reaction Time
Movement Time
Time (s)
Time (ms)
Time (ms)
Probability
Trial Value
Firing Rate (Hz)
Time (min)
Light-On
Center-In
Go Cue
Center-Out
Side-In
Side-Out
Food-Port-In
Trial
Firing Rate (Hz)
Unengaged
Engaged
Ipsi
Contra
Ipsi
Contra
Ipsi
Contra
Unrewarded
Rewarded
Unrewarded
Rewarded
Time (s)

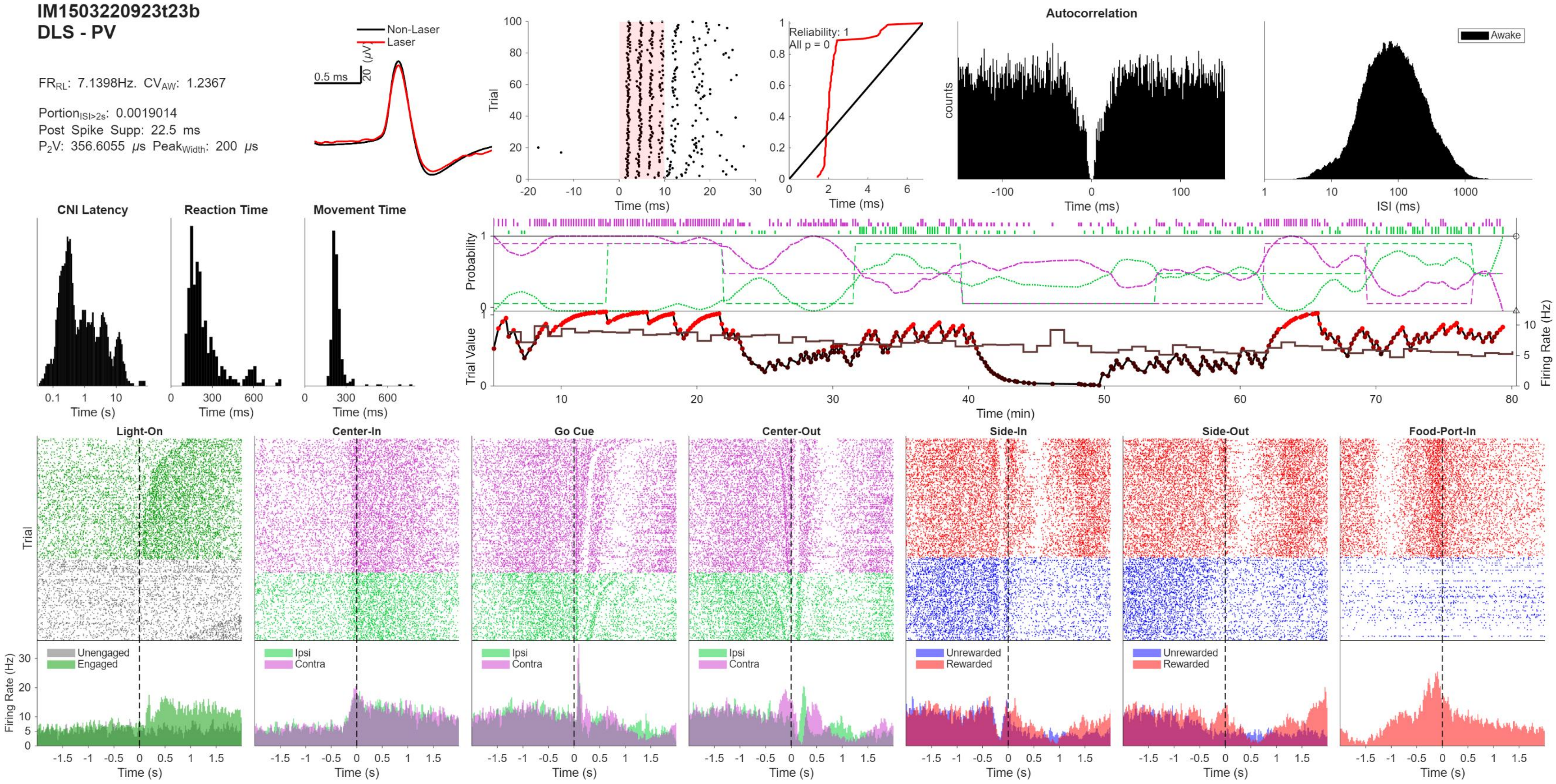
IM1503220923t23b
DLS - PV
FR_RL: 7.1398Hz. CV_AW: 1.2367
Portion_ISI>2s: 0.0019014
Post Spike Supp: 22.5 ms
P2V: 356.6055 μs Peak_Width: 200 μs
Non-Laser
Laser
0.5 ms
20 (μV)
Trial
Time (ms)
Reliability: 1
All p = 0
Time (ms)
Autocorrelation
counts
Time (ms)
Awake
ISI (ms)
CNI Latency
Time (s)
Reaction Time
Time (ms)
Movement Time
Time (ms)
Probability
Trial Value
Firing Rate (Hz)
Time (min)
Light-On
Center-In
Go Cue
Center-Out
Side-In
Side-Out
Food-Port-In
Trial
Firing Rate (Hz)
Unengaged
Engaged
Ipsi
Contra
Ipsi
Contra
Ipsi
Contra
Unrewarded
Rewarded
Unrewarded
Rewarded
Time (s)

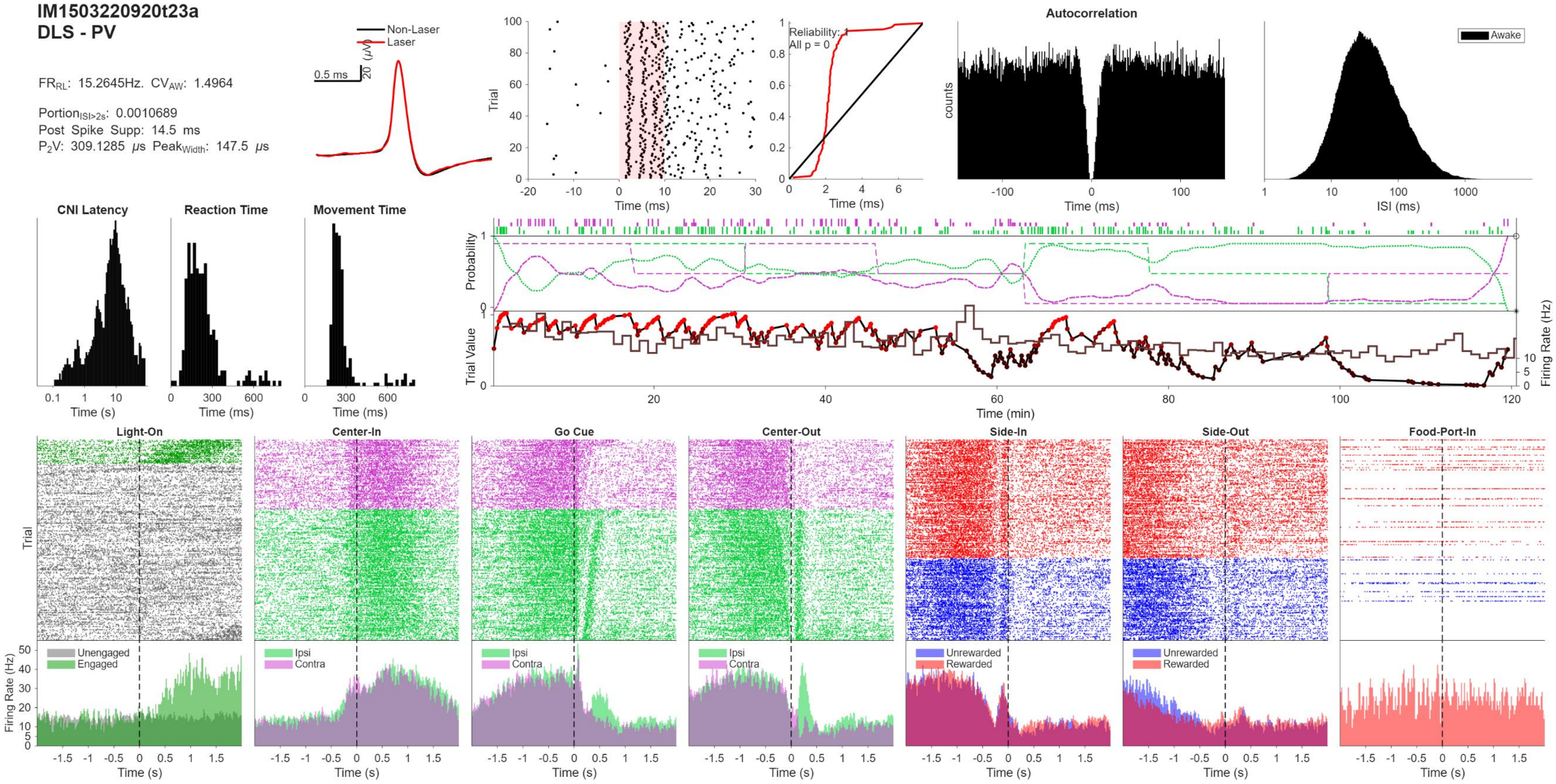
IM1503220920t23a
DLS - PV
FR_RL: 15.2645Hz. CV_AW: 1.4964
Portion_ISI>2s: 0.0010689
Post Spike Supp: 14.5 ms
P2V: 309.1285 μs Peak_Width: 147.5 μs
Non-Laser
Laser
0.5 ms
20 (μV)
Trial
Time (ms)
Reliability: 1
All p = 0
Autocorrelation
counts
Awake
ISI (ms)
CNI Latency
Reaction Time
Movement Time
Time (s)
Probability
Trial Value
Firing Rate (Hz)
Time (min)
Light-On
Center-In
Go Cue
Center-Out
Side-In
Side-Out
Food-Port-In
Unengaged
Engaged
Ipsi
Contra
Unrewarded
Rewarded

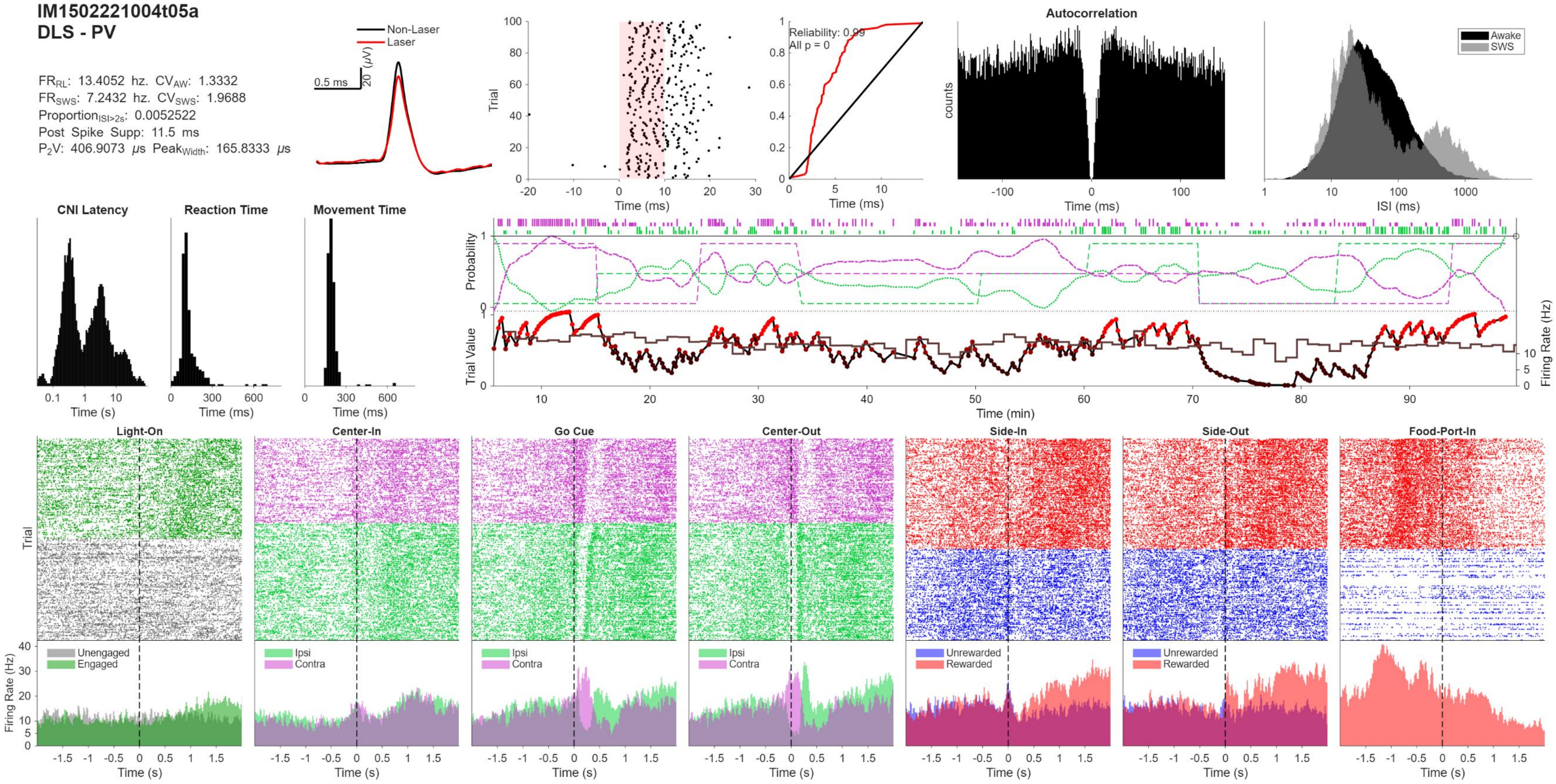
IM1502221004t05a
DLS - PV
FR$_{RL}$: 13.4052 hz. CV$_{AW}$: 1.3332
FR$_{SWS}$: 7.2432 hz. CV$_{SWS}$: 1.9688
Proportion$_{ISI>2s}$: 0.0052522
Post Spike Supp: 11.5 ms
P$_2$V: 406.9073 μs Peak$_{Width}$: 165.8333 μs
Non-Laser
Laser
0.5 ms
20 (μV)
Trial
Time (ms)
Reliability: 0.99
All p = 0
Time (ms)
Autocorrelation
counts
Time (ms)
Awake
SWS
ISI (ms)
CNI Latency
Time (s)
Reaction Time
Time (ms)
Movement Time
Time (ms)
Probability
Trial Value
Firing Rate (Hz)
Time (min)
Light-On
Center-In
Go Cue
Center-Out
Side-In
Side-Out
Food-Port-In
Trial
Firing Rate (Hz)
Unengaged
Engaged
Ipsi
Contra
Unrewarded
Rewarded
Time (s)

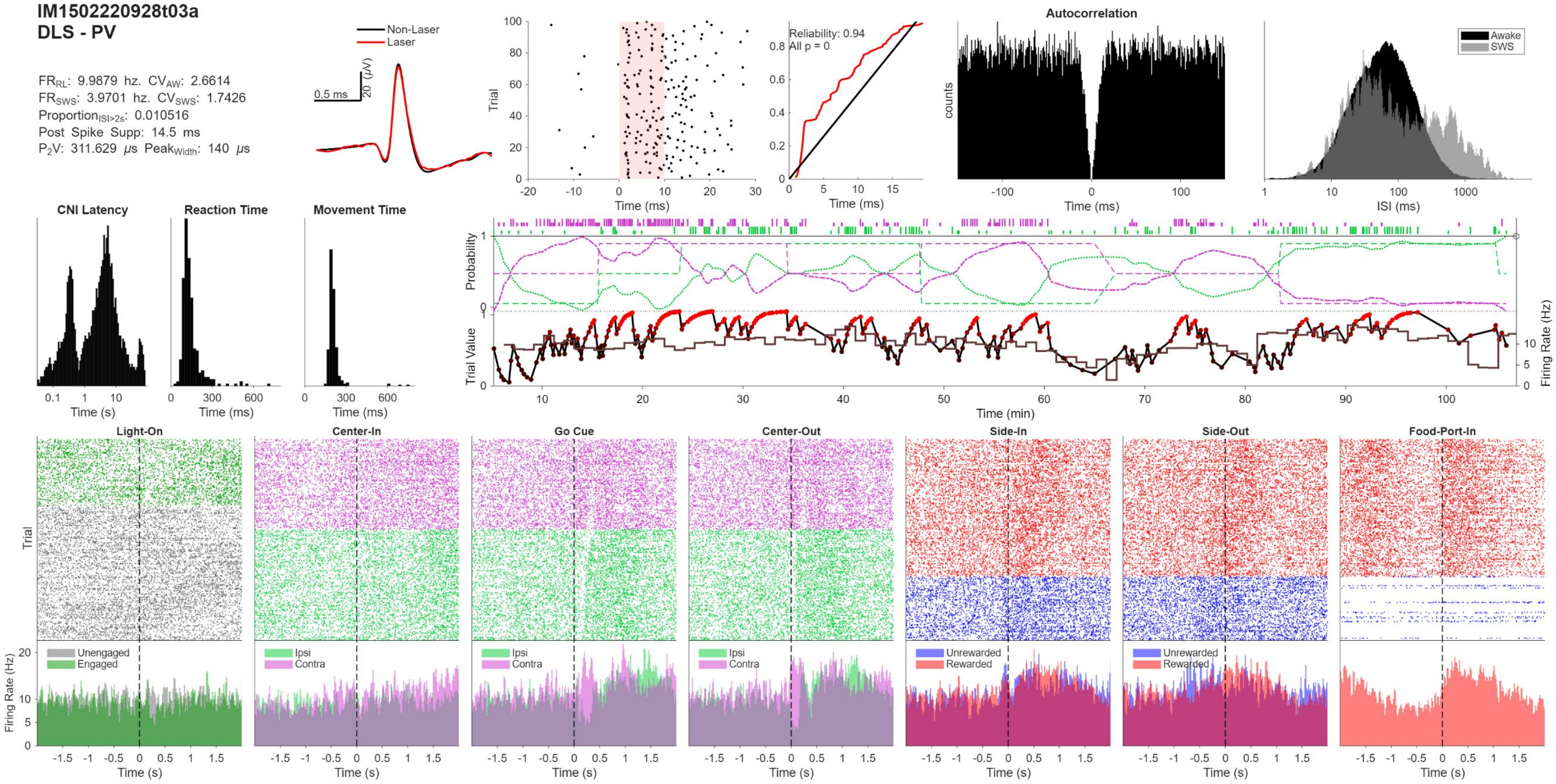
IM1502220928t03a
DLS - PV
FRRL: 9.9879 hz. CVAW: 2.6614
FRSWS: 3.9701 hz. CVSWS: 1.7426
ProportionISI>2s: 0.010516
Post Spike Supp: 14.5 ms
P2V: 311.629 μs PeakWidth: 140 μs
Non-Laser
Laser
0.5 ms
20 (μV)
Trial
Time (ms)
Reliability: 0.94
All p = 0
Time (ms)
Autocorrelation
counts
Time (ms)
Awake
SWS
ISI (ms)
CNI Latency
Reaction Time
Movement Time
Time (s)
Time (ms)
Time (ms)
Probability
Trial Value
Firing Rate (Hz)
Time (min)
Light-On
Center-In
Go Cue
Center-Out
Side-In
Side-Out
Food-Port-In
Trial
Firing Rate (Hz)
Unengaged
Engaged
Ipsi
Contra
Unrewarded
Rewarded
Time (s)

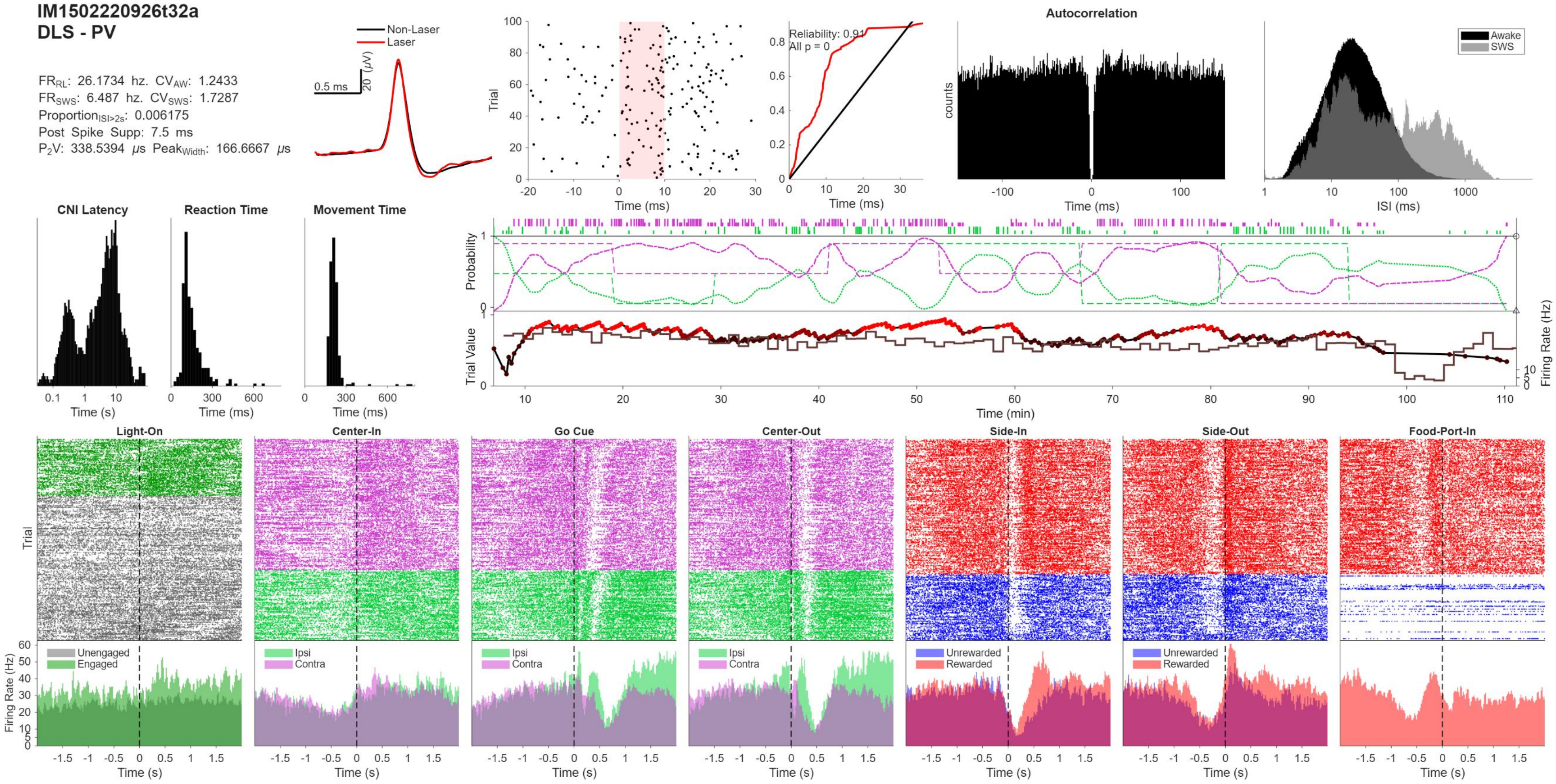
IM1502220926t32a
DLS - PV
FR$_{RL}$: 26.1734 hz. CV$_{AW}$: 1.2433
FR$_{SWS}$: 6.487 hz. CV$_{SWS}$: 1.7287
Proportion$_{ISI>2s}$: 0.006175
Post Spike Supp: 7.5 ms
P$_2$V: 338.5394 μs Peak$_{Width}$: 166.6667 μs
Non-Laser
Laser
0.5 ms
20 (μV)
Trial
Time (ms)
Reliability: 0.91
All p = 0
Time (ms)
Autocorrelation
counts
Time (ms)
Awake
SWS
ISI (ms)
CNI Latency
Time (s)
Reaction Time
Time (ms)
Movement Time
Time (ms)
Probability
Trial Value
Firing Rate (Hz)
Time (min)
Light-On
Center-In
Go Cue
Center-Out
Side-In
Side-Out
Food-Port-In
Trial
Firing Rate (Hz)
Unengaged
Engaged
Ipsi
Contra
Unrewarded
Rewarded
Time (s)
